# Cell transcriptomic atlas of the non-human primate *Macaca fascicularis*

**DOI:** 10.1101/2021.12.13.472311

**Authors:** Lei Han, Xiaoyu Wei, Chuanyu Liu, Giacomo Volpe, Zhenkun Zhuang, Xuanxuan Zou, Zhifeng Wang, Taotao Pan, Yue Yuan, Xiao Zhang, Peng Fan, Pengcheng Guo, Yiwei Lai, Ying Lei, Xingyuan Liu, Feng Yu, Shuncheng Shangguan, Guangyao Lai, Qiuting Deng, Ya Liu, Liang Wu, Quan Shi, Hao Yu, Yunting Huang, Mengnan Cheng, Jiangshan Xu, Yang Liu, Mingyue Wang, Chunqing Wang, Yuanhang Zhang, Duo Xie, Yunzhi Yang, Yeya Yu, Huiwen Zheng, Yanrong Wei, Fubaoqian Huang, Junjie Lei, Waidong Huang, Zhiyong Zhu, Haorong Lu, Bo Wang, Xiaofeng Wei, Fengzhen Chen, Tao Yang, Wensi Du, Jing Chen, Shibo Xu, Juan An, Carl Ward, Zongren Wang, Zhong Pei, Chi-Wai Wong, Xiaolei Liu, Huafeng Zhang, Mingyuan Liu, Baoming Qin, Axel Schambach, Joan Isern, Liqiang Feng, Yan Liu, Xiangyu Guo, Zhen Liu, Qiang Sun, Patrick H. Maxwell, Nick Barker, Pura Muñoz-Cánoves, Ying Gu, Jan Mulder, Mathias Uhlen, Tao Tan, Shiping Liu, Huanming Yang, Jian Wang, Yong Hou, Xun Xu, Miguel A. Esteban, Longqi Liu

**Affiliations:** BGI-ShenZhen, Shenzhen 518103, China; Shenzhen Key Laboratory of Single-Cell Omics, BGI-Shenzhen, Shenzhen 518120, China; Shenzhen Bay Laboratory, Shenzhen 518000, China; College of Life Sciences, University of Chinese Academy of Sciences, Beijing 100049, China; Hematology and Cell Therapy Unit, IRCCS-Istituto Tumori ‘Giovanni Paolo II’, Bari 70124, Italy; School of Biology and Biological Engineering, South China University of Technology, Guangzhou 510006, China; State Key Laboratory for Zoonotic Diseases, Key Laboratory for Zoonosis Research of Ministry of Education, Institute of Zoonosis, College of Veterinary Medicine, Jilin University, Changchun 130062, China; Laboratory of Integrative Biology, Guangzhou Institutes of Biomedicine and Health, Chinese Academy of Sciences, Guangzhou 510530, China; Joint School of Life Sciences, Guangzhou Institutes of Biomedicine and Health and Guangzhou Medical University, Guangzhou 510530, China; Department of Biology, University of Copenhagen, Copenhagen DK-2200, Denmark; China National GeneBank, BGI-Shenzhen, Shenzhen 518120, China; BGI College and Henan Institute of Medical and Pharmaceutical Sciences, Zhengzhou University, Zhengzhou 450000, China; Institute for Stem Cells and Neural Regeneration, School of Pharmacy, State Key Laboratory of Reproductive Medicine, Nanjing Medical University, Nanjing 211166, China; University of Science and Technology of China, Hefei 230026, China; Department of Urology, First Affiliated Hospital, Sun Yat-sen University, Guangzhou 510000, China; Department of Neurology, First Affiliated Hospital, Sun Yat-sen University, Guangzhou 510000, China; Huazhen Biosciences, Guangzhou 510900, China; Department of Orthopedics, Tianjin Medical University General Hospital, Tianjin 300052, China; Laboratory of Metabolism and Cell Fate, Guangzhou Institutes of Biomedicine and Health, Chinese Academy of Sciences, Guangzhou 510530, China; Institute of Experimental Hematology, Hannover Medical School, Hannover 30625, Germany; Division of Hematology/Oncology, Harvard Medical School, Boston MA 02115, USA; Spanish National Center on Cardiovascular Research (CNIC), Madrid E-28029, Spain; State Key Laboratory of Respiratory Diseases, Guangzhou Institutes of Biomedicine and Health, Chinese Academy of Sciences, Guangzhou 510530, China; Jinan University, Guangzhou 510632, China; Institute of Neuroscience, State Key Laboratory of Neuroscience, CAS Key Laboratory of Primate Neurobiology, CAS Center for Excellence in Brain Science and Intelligence Technology, Chinese Academy of Sciences, Shanghai 200031, China; Cambridge Institute for Medical Research, Department of Medicine, University of Cambridge, Cambridge CB2 0XY, United Kingdom; A*STAR Institute of Molecular and Cell Biology, Singapore 138648, Singapore; Department of Experimental and Health Sciences, Pompeu Fabra University (UPF), ICREA and CIBERNED, Barcelona E-08003, Spain; Department of Protein Science, Science for Life Laboratory, KTH-Royal Institute of Technology, Stockholm 17121, Sweden; Department of Neuroscience, Karolinska Institute, Stockholm 17177, Sweden; State Key Laboratory of Primate Biomedical Research, Institute of Primate Translational Medicine, Kunming University of Science and Technology, Kunming 650500, China; James D. Watson Institute of Genome Sciences, Hangzhou 310058, China; Guangdong Provincial Key Laboratory of Genome Read and Write, Shenzhen 518120, China; Institute of Stem Cells and Regeneration, Chinese Academy of Sciences, Beijing 100101, China

## Abstract

Studying tissue composition and function in non-human primates (NHP) is crucial to understand the nature of our own species. Here, we present a large-scale single-cell and single-nucleus transcriptomic atlas encompassing over one million cells from 43 tissues from the adult NHP *Macaca fascicularis*. This dataset provides a vast, carefully annotated, resource to study a species phylogenetically close to humans. As proof of principle, we have reconstructed the cell-cell interaction networks driving Wnt signalling across the body, mapped the distribution of receptors and co-receptors for viruses causing human infectious diseases and intersected our data with human genetic disease orthologous coordinates to identify both expected and unexpected associations. Our *Macaca fascicularis* cell atlas constitutes an essential reference for future single-cell studies in human and NHP.

## MAIN TEXT

Global initiatives such as the Human Cell Atlas are aiming to chart the cell types and cell states of all tissues in the human body using high-throughput single-cell/nucleus RNA-sequencing (sc/snRNA-seq) and other technologies^1–5^. The ultimate goal of these efforts is to create complete reference maps across different ethnic groups, ages, environmental conditions and pathologies. A major obstacle in this endeavour is that accessing a wide range of ‘high quality’ human samples and obtaining enough sample size is complicated by relevant practical and ethical considerations. Model animals (e.g., mouse and rat) are a useful resource to fill knowledge gaps^6–8^, in particular the effects of experimental perturbation, but due to profound phylogenetic differences many developmental, physiological and pathological aspects are not mimicked in humans. Given the evolutionary proximity, NHP present an excellent alternative (the nearest-to-human) when no other suitable models exist. Generating a NHP cell atlas will produce an extensive catalogue of human disease and age-related features that can be modelled in NHP. It will also provide unique insights into the evolutionary and adaptative mechanisms underlying changes in body function between the two species. In this regard, it could for example discover tissue regenerative capacities selectively maintained in NHP and potential ways to boost them in human.

NHP encompass a large and very diverse group of species with major ecological, dietary, locomotor and behavioural differences^9–11^. Because of their close evolutionary proximity to humans among NHP, overall characteristics and wider availability, macaques are primarily employed for research purposes worldwide including human disease modelling and preclinical safety assessment studies^12,13^. Here, we have used adult *Macaca fascicularis* (cynomolgus monkey) to generate the largest single-cell transcriptomic NHP dataset to date, encompassing over 1 million individual cells/nuclei from 43 tissues covering all major systems (nervous, immune, endocrine, cardiovascular, respiratory, digestive, skeletal, reproductive and urinary), all performed with the same droplet-based approach^14^. To facilitate the exploration of this dataset, we have created the first version of the Non-Human Primate Cell Atlas or NHPCA, an open and interactive database (https://db.cngb.org/nhpca/) that will be regularly updated with subsequent sc/snRNA-seq *Macaca fascicularis* datasets focused on development, aging, disease and drug responses, as well as other omics datasets and data from other NHP species.

### Generation of an adult monkey single-cell transcriptomic atlas

We isolated cells/nuclei from 43 different tissue samples from three male and three female six-year-old *Macaca fascicularis* monkeys (**Fig. 1a** **and Supplementary Table 1a**). Bladder (two), cerebellum (two), diaphragm (two), gallbladder (two), kidney (two), liver (three), lung (two), salivary gland (two), subcutaneous (two) and visceral adipose tissue (two) were analyzed as biological replicates to assess individual and gender variability, observing good overlap in all cases (**Extended Data Fig. 1**). Most of the tissues were profiled by snRNA-seq^15–17^, which allows both to circumvent complications associated with stressful dissociation protocols that can alter the cell transcriptome and to profile cells from frozen tissues for removing the need of sample processing immediately after tissue acquisition. However, due to technical limitations in obtaining high quality nuclei, scRNA-seq was performed for colon, duodenum, spleen, stomach, lymph node and bone marrow. Peripheral blood mononuclear cells (PBMC) were also profiled using scRNA-seq. All experiments used the DNBelab C4 droplet-based platform for library generation^14^. To ensure quality, all cells with a gene count lower than 500 and/or mitochondrial content higher than 10% were excluded. We also applied DoubletFinder to detect and remove doublets, which accounted for roughly 5% of the estimated total cell/nuclei. Overall, we retained transcriptomic data for a total of 1,084,164 cells/nuclei (**Fig. 1a**), with numbers ranging from 99,123 in the cerebellum to 2,039 in the duodenum (**Supplementary Table 1a**). Global visualization of cell clustering using Uniform Manifold Approximation and Projection (UMAP) showed that each tissue clusters separately, with tissues from the same system generally clustering closer (**Fig. 1a, b** **and Extended Data Fig. 2-6**). We then performed individual UMAP representations for each tissue and applied unbiased graph-based Seurat clustering, which identified 463 cell clusters among all tissues (**Extended Data Fig. 7-10**). Based on the expression levels of cell type-specific markers (**Extended Data Fig. 11**), we identified 106 cell types in the global UMAP view of all tissues (**Fig. 1c** **and Supplementary Table 1b, c**). These were roughly categorized into epithelial cells (40 clusters), immune cells (13 clusters), endocrine cells (11 clusters), muscle cells (9 clusters), stromal cells (7 clusters), endothelial cells (7 clusters), neurons (7 clusters), glia (7 clusters), mesothelial cells (3 clusters), adipocytes (1 cluster) and unknown cells (1 cluster from carotid). On average, we detected 1,368 genes and 3,024 unique molecular identifiers (UMI) per cell. The median gene count per tissue varied between 3,016 in the neocortex and 736 in the case of PBMC, while UMI ranged between 8,015 for the neocortex and 1,313 for the prostate (**Extended Data Fig. 12**). The number of cells for each of these 106 cell types ranged from 87,890 granule cells in the cerebellum to 37 bone marrow stromal cells (**Extended Data Fig. 13**). Reassuringly, many of the 106 clusters were largely composed of a cell type belonging to a specific tissue, such as cerebellar granule cells in cluster 45, hepatocytes in clusters 87 and 88, epididymis stereociliated cells in cluster 29 and salivary acinar cells in cluster 83 (**Fig. 1c** **and Extended Data Fig. 14a**). However, cell types such as endothelial, stromal and various immune cells were shared between different tissues, as expected (**Extended Data Fig. 14b**). A detailed annotation of all cell populations detected in every tissue is provided in **Extended Data Figure 7-10 and Supplementary Table 1d, e**. Our *Macaca fascicularis* atlas is the largest NHP single-cell transcriptome dataset to date and can be explored interactively by tissue, cell type and gene through our NHPCA database.

**Figure 1.**
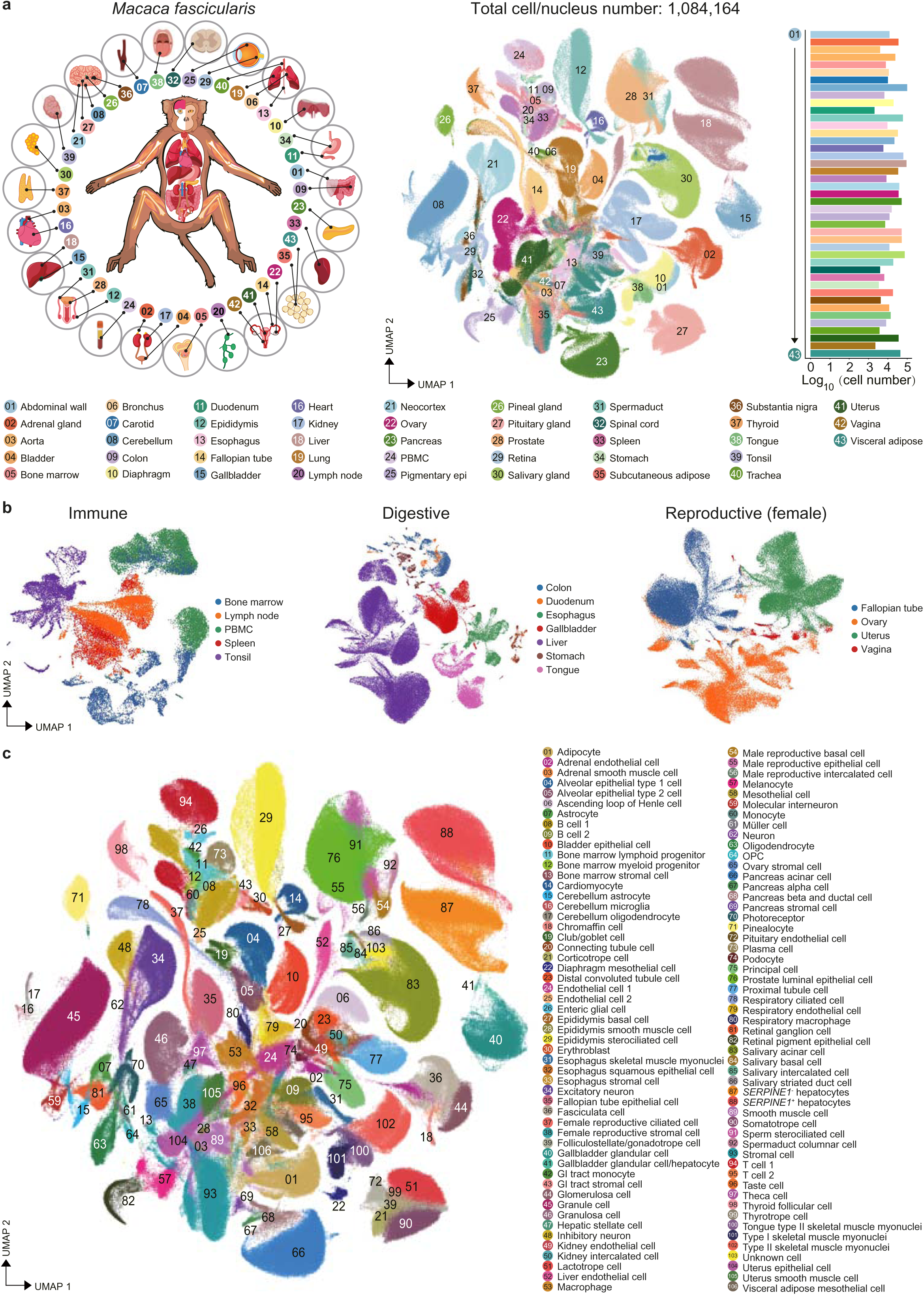
Generation of a single-cell atlas across 43 tissues of *Macaca fascicularis* monkey. **(a)** Schematic representation of monkey tissues analyzed in this study (top left panel). A total of 43 tissues were collected from three male and three female 6-year-old monkeys. UMAP visualization of the global clustering indicating all single cells from the dataset colored by tissue (top middle panel) and bar plot showing the number of cells/nuclei profiled for every tissue after passing the quality control (top right panel). N = 1,084,164 individual nuclei/cells analyzed. **(b)** UMAP visualization of tissues grouped by specific systems such as immune system (bone marrow, peripheral blood, spleen, tonsil and lymph node), digestive system (colon, duodenum, esophagus, gallbladder, liver, stomach and tongue) and female reproductive system (fallopian tube, ovary, uterus and vagina). **(c)** UMAP visualization of all clusters colored by major cell types. A total of 106 cell clusters were identified in the dataset. Cell type annotation for all major clusters is provided in the right-hand side legend. *SERPINE1* was used to discriminate two distinct cluster of hepatocytes.

### Common cell types across monkey tissues

We inspected whether common cell types distributed throughout different tissues in the monkey body display tissue-specific transcriptional programs^3,18–20^. First, we selectively clustered stromal cells, macrophages (including microglia), endothelial cells and smooth muscle cells from all sequenced tissues. While observing a considerable diversity, many cell clusters grouped together on the basis of tissue origin, such as stromal cells from the female reproductive system, microglia from the central nervous system, endothelial cells from the respiratory system and smooth muscle cells from the male reproductive system (**Extended Data Fig. 15a-d**). We also performed differentially expressed gene (DEG) analysis to obtain tissue-specific signatures, revealing a substantial heterogeneity among these common cell types across all tissues (**Extended Data Fig. 15e-h and Supplementary Table 2a-d**).

Our transcriptomic profiling of single nuclei offers the possibility of studying cell populations that cannot be characterized by conventional scRNA-seq analysis, such as myonuclei from multinucleated skeletal muscle fibers. We grouped and re-clustered cells from tissues in our atlas known to contain skeletal muscle cells (diaphragm, tongue, esophagus and abdominal wall). This showed two distant populations in abdominal wall and diaphragm, whereas nuclei from esophagus and tongue where more concentrated (**Fig. 2a**). The separation of nuclei in abdominal wall and diaphragm corresponded to *MYH7^+^* type I (slow-twitch) and *MYH2^+^* type II (fast-twitch) myofibers^21^ (**Fig. 2b, c** **and Supplementary Table 2e-g**). In contrast, type I and type II tongue myonuclei were in close vicinity, which may be related to the tongue being a highly innervated muscle^22^. Differential threshold of *MYH2* and *GPD2* further subdivided type II myonuclei into type IIa (*MYH2*^high^) and type IIb (*MYH2*^low^ *GPD2*^+^). In addition, we discriminated, albeit at low proportions, *NAV3^+^* neuromuscular junction (NMJ) nuclei in the diaphragm and *ETV5*^+^ myotendinous junction (MTJ) nuclei in both tongue and diaphragm (**Fig. 2b-d**). Moreover, we detected *PAX7*^+^ nuclei from satellite cells (the stem cells from the skeletal muscle lineage), and a small cluster of *LVRN*^+^ fibroadipogenic progenitors (FAP) could be annotated in the diaphragm. Skeletal muscle nuclei displayed subtype-specific and tissue-specific gene expression signatures and gene ontology (GO) terms (**Fig. 2e, f and Extended Data Fig 16a-c**). We also noticed substantial myonuclei heterogeneity within the same subtype and tissue (**Fig. 2f**).

**Figure 2.**
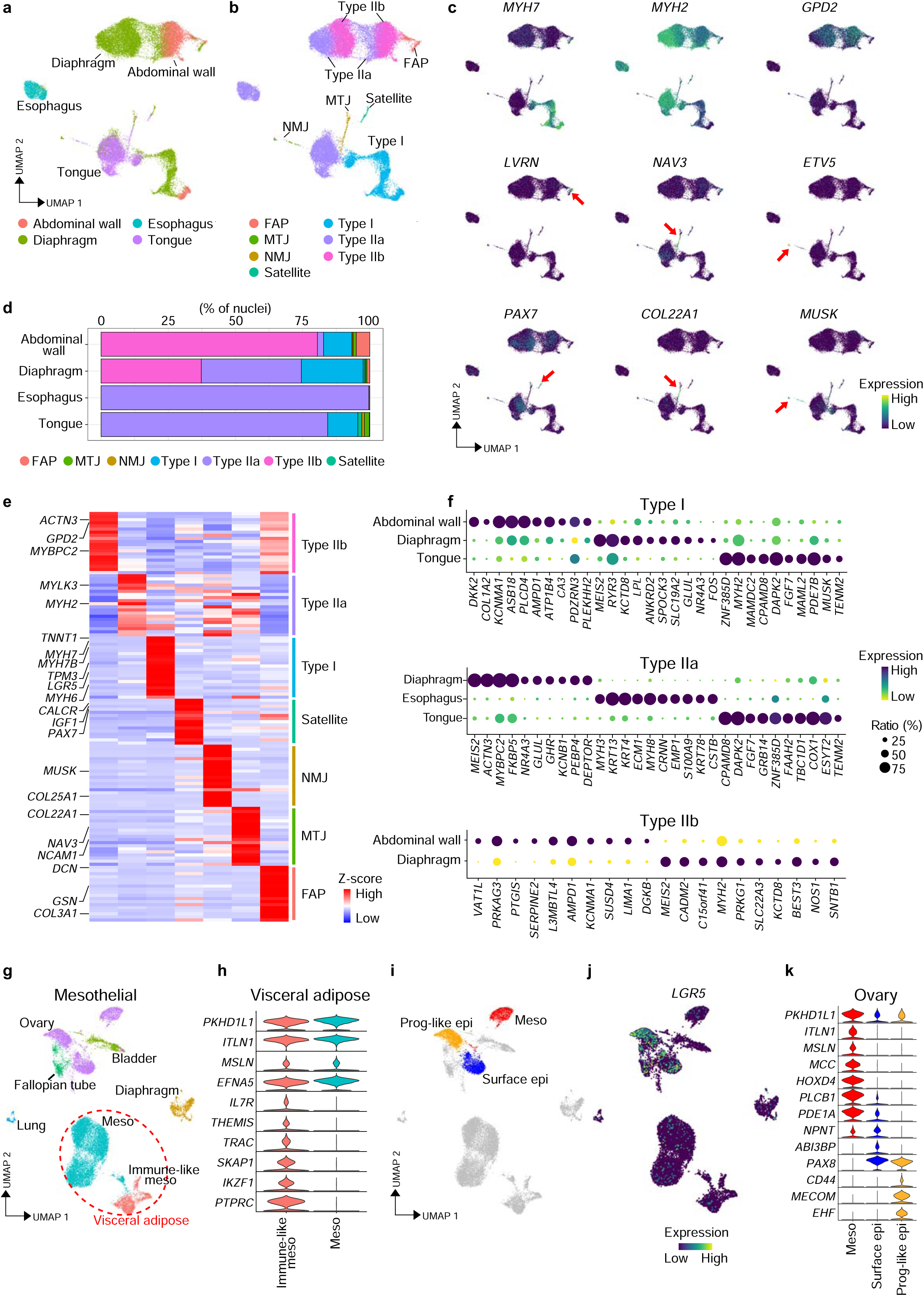
Characterization of skeletal myofibers and mesothelial cells. **(a)** UMAP visualization of the global clustering of skeletal muscle cells annotated in our dataset. Clusters are colored by tissue (abdominal wall, diaphragm, esophagus and tongue). Due to their low number, fallopian tube, vagina and tonsil skeletal cells were excluded from this analysis. Endothelial and immune cells were not included in this analysis. **(b)** UMAP representation of all re-clustered skeletal muscle cells colored by subtype. **(c)** UMAP visualization of specific markers used to identify type I (*MYH7*), type IIa (*MYH2*) and type IIb myonuclei (*GPD2*), FAP (*LVRN*), MTJ (*NAV3* and *COL22A1*), NMJ (*ETV5* and *MUSK*) and satellite cells (*PAX7*), as shown in **b**. Due to their small proportions, the latter three populations are highlighted by a red arrow. **(d)** Stacked bar plot representing the proportion of skeletal muscle nuclei (myonulcei subtypes type I, type IIa, type IIb, MTJ and NMJ, and also satellite cells and FAP) in the indicated tissues. **(e)** Heatmap showing DEG among the skeletal muscle populations highlighted in **d**. **(f)** Bubble plot showing DEG for each of the myonuclei subtypes comparing different tissues. **(g)** UMAP visualization of mesothelial cells from the selected tissues (bladder, diaphragm, fallopian tube, lung, ovary and visceral adipose tissue). Two different clusters of mesothelial cells in visceral adipose tissue are indicated by the red dotted line. **(h)** Violin plot showing the differential expression of mesothelial and immune markers in the two visceral adipose tissue clusters highlighted by the red dotted line in panel **g**. **(i)** UMAP visualization of three different clusters of mesothelial cells from the ovary (left panel). Mesothelial cells (Meso), surface epithelial (Surface epi) and progenitor-like epithelial (Prog-like epi) cells are highlighted in red, blue and yellow, respectively. **(j)** UMAP visualization of *LGR5* expression in ovarian cells. **(k)** Violin plot showing the DEG among the three populations of ovarian cells highlighted in the UMAP.

Next, to explore the heterogeneity between different types of adipocytes, we grouped and re-clustered cells from subcutaneous and visceral adipose tissues, resulting in 10 major clusters (**Extended Data Fig. 17a**). We observed a marked distinction between mature adipocytes and adipocyte progenitors, as reflected by the differential expression of *ADIPOQ* and *CD34* (**Extended Data Fig. 17b**). Visceral mature adipocytes and adipocyte progenitors displayed enriched expression of *ITLN1*, in agreement with visceral adipocytes having mesothelial origin^23^, and also high mitochondrial activity exemplified by high expression of *ND4*, *ATP6* and *COX3* ^24,25^ (**Extended Data Fig. 17c, d**). In contrast, subcutaneous mature adipocytes and adipocyte progenitors were enriched in *FOS*. Likewise, *SLC11A1* and *SPOCK3* marked mature subcutaneous and visceral adipocytes, respectively. Adipocyte progenitors contained two populations for visceral tissue (*WT1*^+^ and *CFD*^high^), three for subcutaneous tissue (*ESR1*^+^, *CXCL14*^+^*APOD*^+^ and *DPP4*^+^) and one shared between both tissues (*NOX4*^+^) (**Extended Data Fig. 17a, c and d**). Within the subcutaneous *CXCL14*^+^*APOD*^+^ progenitor cluster, we observed a population of *CFD*^high^ cells that also co-expressed *DPP4*, a marker of highly proliferative adipocyte progenitors in both mouse and human^26^. However, we did not detect significant proliferation in any of the monkey adipocyte progenitor populations based on the expression of the pan-cycling marker *MKI67*^27^ (**Extended Data Fig. 17c**). *NOX4*^+^ is an NAPDH oxidase that acts as a switch from insulin-induced proliferation to adipocyte differentiation, suggesting that the shared cluster is a converging route for both adipose tissues towards adipocytic maturation^28^.

Finally, we grouped and re-clustered all tissues that contain mesothelial cells, a type of specialized epithelial cells. Mesothelial cells from bladder, ovary and fallopian tube were in close proximity while those from other tissues clustered more separately (**Fig. 2g**). We also observed intra-tissue heterogeneity, in particular for visceral adipose tissue and ovary. In the former, we observed a cluster of immune-like mesothelial cells that, aside from the expression of the typical mesothelial markers (*MSLN*, *ITLN1* and *PKHD1L1*), express high levels of immune cell markers (e.g., *PTPRC*, *IL7R* and *TRAC*) (**Fig. 2h**). This is in agreement with the emerging concept that structural cells display immune cell properties^3,18^ and the known immunomodulatory role of visceral adipose tissue in responses to bacteria in the gut^29^. Interestingly, in the ovary, we identified a classical mesothelial population and two close *PAX8*^+^ epithelial-like populations (one mature and one progenitor-like) of mesothelial origin^30^ (**Fig. 2i-k**). Progenitor-like ovarian epithelial cells expressed well-known stem cell markers such as *LGR5*, *MECOM* and *CD44*^31^.

These findings add up to the growing understanding of common cell type heterogeneity and tissue-specific molecular signatures^3,18–20^. Our data provide a new resource for further dissecting these differences, clarifying the underlying mechanisms and studying interspecies differences^32^.

### Analysis of Wnt signaling components identifies potential stem cell populations

A single-cell body atlas of large dimensions like ours is ideal for the systematic investigation of multifaceted cell-cell interactions including those occurring in cytokine or growth factor-mediated signaling pathways such as the Wnt (wingless-related integration site) pathway^33,34^. Besides playing essential roles in embryonic development, Wnt factors control growth and maintenance of numerous tissues throughout life. Consistently, Wnt signaling effects are associated with the regulation of adult stem cell function^35^. To exert this role, Wnt factors bind to specific receptors (FZD, frizzled) and co-receptors (LRP, low-density lipoprotein receptor related protein). In addition, LGR (leucine rich repeat containing G protein-coupled receptor) proteins (LGR4, 5 and act as amplifiers of Wnt signals by inhibiting negative regulators^36^. Accordingly, LGR5 and 6 often mark and regulate adult homeostatic and facultative stem cells, mostly of epithelial origin, in multiple mammalian tissues, whereas LGR4 has a widespread distribution and less clear function. We thus performed a survey of LGR proteins throughout the monkey body to thoroughly dissect cells targeted by the Wnt pathway and identify previously unappreciated stem cell populations. In this regard, it is worth noting that the majority of reports of LGR5-expressing cells to date have been performed with genetically engineered mouse models due to the lack of specific tools and reagents to study other mammals^36^.

*LGR5* was detected across several monkey tissues, unexpectedly with the highest expression in type I skeletal muscle myonuclei, epithelial cells of the uterus and fallopian tube, oligodendrocyte progenitor cells (OPC) and renal distal convoluted tubule cells (DCTC) (**Fig. 3a**). With the exception of epithelial cells in the uterus and fallopian tube^36^, these tissues have not previously been reported to contain LGR5^+^ cells in mammalian adulthood. The expression of *LGR6* appeared to be more restricted (**Extended Data Fig. 18a**), with higher abundance in cardiomyocytes, thyroid follicular cells, folliculostellate cells of the pituitary gland and the previously reported smooth muscle cells^37^ **(Extended Data Fig. 19-22)**. We also detected *LGR5*^+^ or *LGR6*^+^ cells in selected cell populations of numerous other tissues including both previously reported (e.g., ovary epithelial cells^31^, hepatocytes^38^ and colon enterocytes^39^) and unreported (e.g., *LGR5*^+^ cells in bipolar cells of the retina^40^) (**Fig. 3a, Extended Data Fig. 18a and 19-22**). In general, *LGR5* and *LGR6* did not overlap, apart from fallopian tube epithelial cells and vagina smooth muscle cells (**Extended Data Fig. 18b)**. Moreover, we observed little overlap between *LGR5*^+^ or *LGR6*^+^ cells with those expressing *MKI67*, apart from epithelial cells of the fallopian tube and uterus and basal cells of the salivary gland **(Extended Data Fig. 19-22 and Supplementary Table 3a-c)**. In contrast to *LGR5* and *6*, *LGR4* was ubiquitously expressed across most tissues, with the highest expression in pancreatic acinar, beta and ductal cells, Müller cells of the retina and adipocytes (**Extended Data Fig. 18c**).

**Figure 3.**
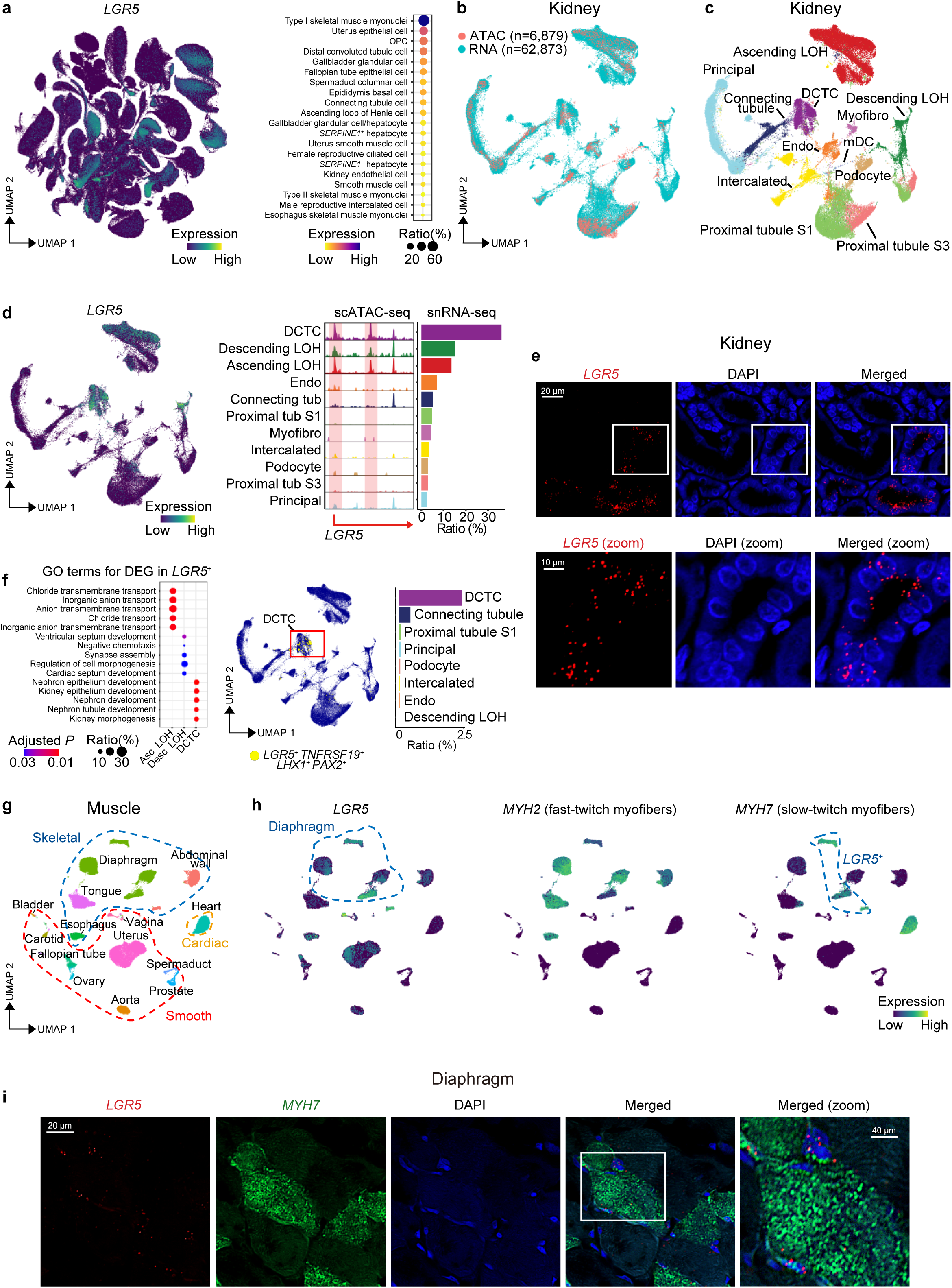
Analysis of *LGR5*^+^ cells across all monkey tissues. **(a)** UMAP visualization of *LGR5* expression across all tissues profiled in this study. The bubble plot on the right shows the *LGR5* expression ratio in the indicated cell types. **(b)** Co-embedding of kidney snRNA-seq (highlighted in blue) and scATAC-seq (highlighted in red) datasets. **(c)** UMAP visualization of integrated kidney snRNA- and scATAC-seq data. Cell clusters are colored according to cell identity. Abbreviations: DCTC, distal convoluted tubule cells; Endo, endothelial cells; LOH, loop of Henle; mDC, myeloid-derived dendritic cells; Myofibro, myofibroblasts. **(d)** UMAP visualization of *LGR5* across kidney cell types and ArchR track visualization of aggregate scATAC-seq signals on the *LGR5* locus in each cell type annotated in **c**. The bar plot on the right side indicates the ratio (%) of *LGR5*^+^ cells in each cell type of kidney. **(e)** Representative image of smFISH detection for *LGR5* expression in DCTCs (scale bar 20 μm). The bottom panel represents a magnification of the area indicated by the white box in the top panel. **(f)** GO analysis showing the pathways associated to the DEGs obtained by comparing *LGR5*^+^ cells from DCTC, ascending and descending LOH. The UMAP and the barplot on the right highlight the presence and the percentage of *LGR5*^+^ cells co-expressing the progenitor markers *PAX2*, *TNFRSF19* and *LHX2*. **(g)** UMAP visualization of all muscle cell types annotated in our dataset clustered by tissue (abdominal wall, aorta bladder, carotid, diaphragm, esophagus, fallopian tube, heart, ovary, prostate, spermaduct, tongue, uterus and vagina). The dotted lines group clusters of cells belonging to a specific muscle type (cardiac, skeletal and smooth muscle). **(h)** UMAP visualization of *LGR5*, *MYH2* and *MYH7* across all skeletal muscle cell types. The blue dotted line in the left panel indicates all clusters belonging to the diaphragm while the one in the right panel indicates *LGR5*^+^ cells. **(i)** Representative image of smFISH detection for *LGR5*, *MYH7* and their co-expression in skeletal myonuclei of the diaphragm (scale bar 20 μm). The panel of the right is a magnification of the area indicated by the white box.

In the kidney, *LGR5*^+^ cells were mostly enriched in the DCTC and to a lesser extent in the descending and ascending loop of Henle (**Fig. 3a** **and Extended Data Fig. 20**). To support this observation, we performed single-cell Assay for Transposase Accessible Chromatin sequencing (scATAC-seq) of monkey kidney and integrated the results with our kidney snRNA-seq data dataset (N = 6,879) (**Fig. 3b, c and Extended Data Fig. 23a, b**). The analysis showed peaks of open chromatin at the *LGR5* promoter and a putative enhancer open in the same cell types expressing *LGR5* (**Fig. 3d**). As validation, we performed single-molecule fluorescence *in-situ* hybridization (smFISH) for *LGR5*, which showed strong expression in selected kidney tubules (**Fig. 3e**). Moreover, GO analysis of DEG comparing the *LGR5*^+^ fractions of DCTC, ascending and descending loop of Henle revealed the enrichment of pathways involved in kidney development in DCTC (**Fig. 3f**), suggesting the possibility that these are progenitor cells. This was strengthened by the observation that DCTC *LGR5*^+^ cells co-express renal progenitor cell markers such as *PAX2*, *LHX1* and *TNFRSF19*^41,42^. We also integrated our data with available human^43^ and mouse^44^ kidney snRNA-seq datasets. Despite observing good integration, we noticed very little, or no, *LGR5* expression in those adult human or mouse kidney datasets^45^ (**Extended Data Fig. 24a-c**).

In the neocortex, integration of available human^46^ and our own mouse snRNA-seq data with our monkey data pointed as well at differential *LGR5* expression patterns between species. *LGR5* expression was highest in OPC in monkey and in oligodendrocytes in human, whereas in mouse it was higher in inhibitory neurons than OPC and oligodendrocytes (**Extended Data Fig. 25a-c**). Pseudotime ordered by Monocle 2 of the OPC maturation trajectory towards oligodendrocyte showed concentration of *LGR5* in monkey OPC (**Extended Data Fig. 25d, e**). Likewise, double immunofluorescence for the OPC marker PDGFRA and LGR5 confirmed their co-expression in OPC from monkey neocortex (**Extended Data Fig. 25f**). The observation that type I skeletal myonuclei and cardiomyocytes ranked first in expression of *LGR5* and *LGR6* in monkey tissues, respectively, was intriguing (**Fig. 3a** **and Extended Data Fig. 18a**). To inspect this further, we grouped and re-clustered all types of muscle cells (skeletal, smooth and cardiac) in our atlas (**Fig. 3g**). *LGR5* was more enriched in *MYH7*^+^ slow-twitch myonuclei of the abdominal wall and diaphragm (**Fig. 3h**), whereas *LGR6* was higher in cardiomyocytes and smooth muscle cells (aorta, ovary, carotid and vagina) (**Extended Data Fig. 26a**). *LGR5* and *LGR6* expression in slow-twitch skeletal myonuclei and in cardiomyocytes, respectively, were validated by smFISH (**Fig. 3i** **and Extended Data Fig. 26b**). In mouse, *LGR5* is known to be expressed in NMJ myonuclei^47^ and a subset of satellite cells activated upon injury^48^, but we did not detect *LGR5* enrichment in either cell type in our monkey dataset (**Extended Data Fig. 19**). The lack of *LGR5* enrichment in monkey satellite cells is unsurprising given that we did not apply any injury to the skeletal muscle tissues profiled. Yet, we could detect *LGR6* in cardiomyocytes using previously reported mouse and human snRNA-seq datasets^49,50^ (**Extended Data Fig. 26c, d**). Similarly, *LGR6* was enriched in several monkey pituitary cell populations, being most highly expressed in folliculostellate cells, which have been reported to be pituitary gland stem cells^51^ (**Extended Data Fig. 26e**). Consistently, those cells also expressed other progenitor markers such as *SOX2*, *PAX6*, *CD44* and *CXCR4* (**Extended Data Fig. 26f**). Moreover, GO analysis of DEG specific to this *LGR5*^+^ population compared to other pituitary cells showed enrichment of terms related to development (**Extended Data Fig. 26g**).

Next, we profiled the genes encoding Wnt factors and the R-spondin family (RSPO1-4) of ligands for LGR proteins^35,36^ in a panel of monkey tissues containing cells with high *LGR5* (kidney, epididymis, fallopian tube, liver, ovary, neocortex and diaphragm) and *LGR6* (heart and pituitary gland) expression (**Extended Data Fig. 27a, b and 28-31**). This allowed us to dissect the potential cell-cell interaction networks driving Wnt signalling throughout the monkey body. Notably, RSPO cytokines were widely distributed but displayed higher expression in mesenchymal-like cells (e.g., smooth muscle cells of epididymis, hepatic stellate cells and folliculostellate cells of the pituitary gland) and mesothelial cells (e.g., of diaphragm, fallopian tube and ovary) of different tissues. Interestingly, *RSPO2* was also high in inhibitory neurons of the neocortex (**Extended Data Fig. 30**). The expression of Wnt factors was more limited and in general lower than RSPO cytokines but we noticed high levels of *WNT9B* in principal cells of the collecting duct in kidney (**Extended Data Fig. 27a, c**), *WNT2B* in mesothelial cells of the fallopian tube (**Extended Data Fig. 29a**) and ovary (**Extended Data Fig. 30c**), and as expected *WNT2* in endothelial cells of the liver^52^ (**Extended Data Fig. 29c**). Wnt9b is an essential regulator of kidney embryonic development in multiple species and of kidney regeneration in lower vertebrates^53^. Supporting the snRNA-seq data, scATAC-seq analysis of the *WNT9B* locus revealed increased enhancer accessibility in monkey principal cells compared to other kidney cell types (**Extended Data Fig. 27d**). In contrast, we detected low *WNT9B* expression in available mouse^44^ and human^46^ snRNA-seq datasets (**Extended Data Fig. 27e**). WNT9B may be responsible for inducing *LGR5* (a Wnt pathway target) in a fraction of DCTC, potentially creating a feedback loop that amplifies WNT9B signals to keep those cells in a progenitor state. In fact, Wnt factors are known to act predominantly on neighbouring cells^33,35^, and cells of the collecting duct and DCTC are in closer proximity than other nephron structures (**Extended Data Fig. 27f**). We further included Wnt receptors and other co-receptors^54^ in the analysis, and also the TCF family of transcription factors bound by β-catenin^55^, as a resource for additional exploration in these tissues (**Extended Data Fig. 27a,b and 28-31**).

Therefore, we have reconstructed the Wnt signaling network in monkey tissues and identified cell types with potential progenitor or homeostatic characteristics. Additional signaling pathways and/or ligand-receptor interactions can be explored through our NHPCA database.

### Prediction of viral infection vulnerability in monkey tissues

To demonstrate the utility of our atlas for advancing the knowledge of disease pathogenesis, we first mapped the expression of the main viral receptors/co-receptors for a panel of 126 viruses including respiratory ones across all monkey tissues. As expected, *NCAM1* (cytomegalovirus receptor) was enriched in astrocytes, oligodendrocytes and neurons, consistent with the knowledge of this virus attacking the central nervous system^56^. In contrast, *CD46*^57^ (receptor for Measles and Herpes viruses) was enriched in epithelial cells from bladder, female and male reproductive system, and liver endothelial cells (**Fig. 4a, Extended Data Fig. 32 and Supplementary Table 4a**). Given the emergency state of the current COVID-19 pandemic caused by SARS-CoV-2^58^, we focused on its receptor *ACE2* and co-receptor *TMPRSS2*^59^ to assess how widespread and homogeneous their expression is in monkey tissues. This offers the major advantage of studying COVID-19 pathogenesis in a species phylogenetically close to humans^60^, and also provides the possibility of profiling cell types and/or tissues that have not been studied in human. In this regard, although the lung is the predominantly affected tissue in COVID-19, it is important to clarify what other tissues are targeted to better understand the disease course and its transmissibility^61^. *TMPRSS2* displayed a broad expression across multiple monkey tissues, whereas *ACE2* had a more restricted pattern. The highest *ACE2* expression was found in epithelial cells from gallbladder (glandular cells), kidney (mostly proximal tubule cells), lung (ciliated, club and alveolar type 2 [AT2] cells) and liver (hepatocytes and cholangiocytes) (**Fig. 4b, Extended Data Fig. 33, 34 and Supplementary Table 4b**). *ACE2* in these tissues was remarkably heterogeneous, suggesting that regulatory mechanisms fine-tune its expression levels. Notably, double positive (*ACE2*^+^ *TMPRSS2*^+^) cells have a higher risk of infection by SARS-CoV-2^59^ but it remains unclear what tissues and cell types throughout the human body co-express these genes. We noticed the largest overlap between *ACE2* and *TMPRSS2* in monkey gallbladder cells in agreement with reports of COVID-19 patients developing acute cholecystitis^62^. Significant co-expression was also observed in ciliated and club cells of the lung, as expected^63,64^, and, interestingly, proximal and connecting tubule cells of the kidney. A smaller overlap was observed in hepatocytes, bladder epithelial cells and pancreatic beta and ductal cells (**Fig. 4c**). Next, we performed a comparative analysis of *ACE2* and *TMPRSS2* distribution in human^3,6,43^ and monkey. A similar distribution was seen in both the gallbladder and liver in the two species, while distinct patterns were observed for proximal tubule cells of the kidney and for ciliated and AT2 cells of the lung (**Extended Data Fig. 35a**). This is important because it implys a mechanism by which the infection with SARS-CoV-2 in the two species could have different consequences.

**Figure 4.**
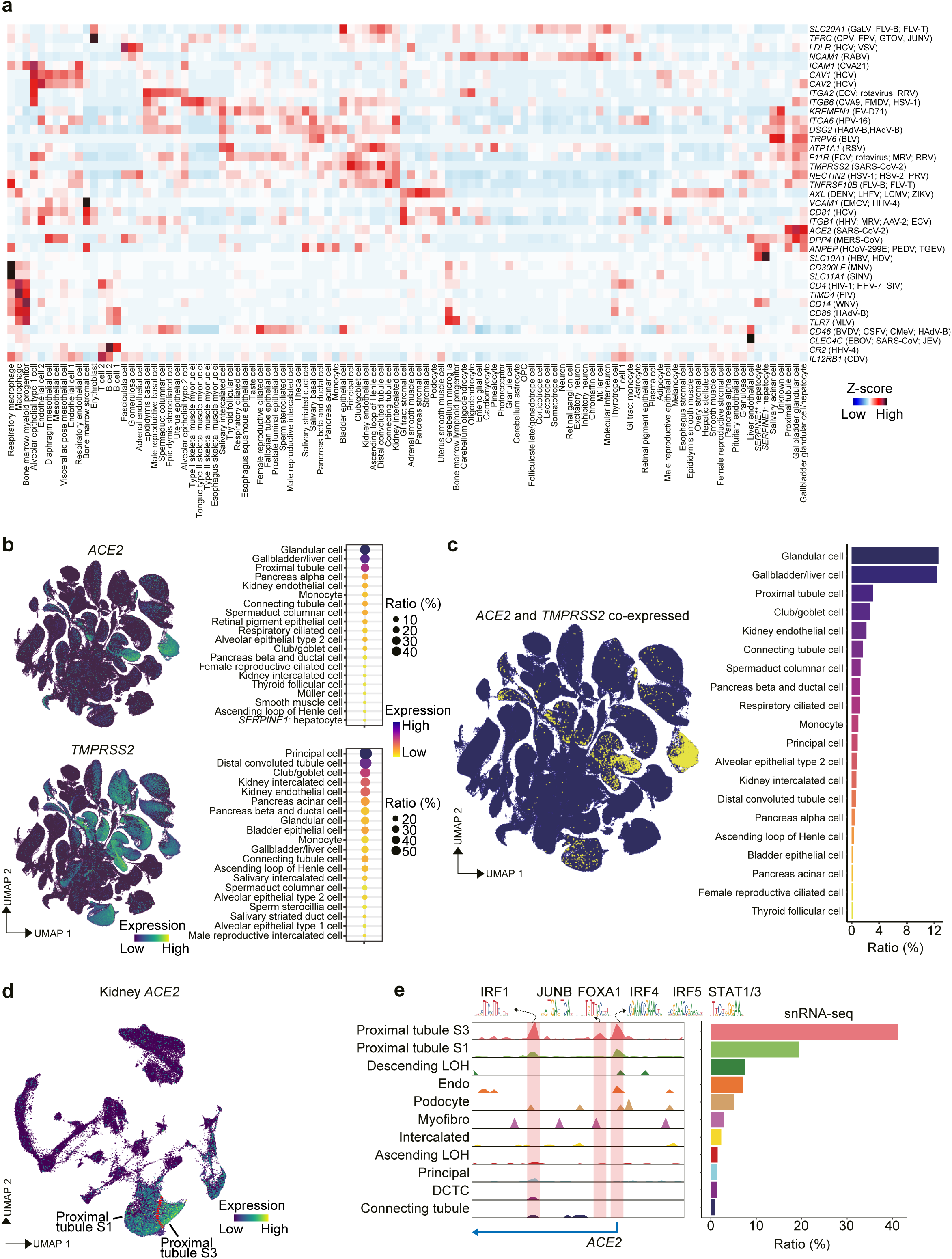
Global analysis of *ACE2* and *TMPRSS2* across monkey tissues. **(a)** Heatmap showing the expression of entry receptors for a selection of the most common viruses (indicated on the left) in all cell clusters annotated in our dataset (indicated at the bottom). **(b)** UMAP visualization of *ACE2* (top) and *TMPRSS2* (bottom) expression in all single cells from our dataset. The bubble plot next to each UMAP shows the expression levels of *ACE2* and *TMPRSS2* in the indicated cell types. The color of each bubble represents the levels of expression and the size indicates the proportion of expressing cells. **(c)** UMAP projection of *ACE2^+^*/*TMPRSS2^+^* cells (highlighted in yellow). The bar plot on the right represents the ratio of cells that co-express both genes. **(d)** UMAP visualization of *ACE2* in the integrated scATAC-seq and snRNA-seq from monkey kidney. **(e)** ArchR track visualization of aggregate scATAC-seq signals on the *ACE2* locus in each on the annotated cell types of the kidney. Predicted binding of human transcription factor predicted based on DNA sequence is shown in the corresponding open chromatin regions of *ACE2*. The bar plot on the right indicates the ratio (%) of *ACE2*^+^ cells in each annotated cell type of the monkey kidney.

As a representative tissue with high but heterogeneous *ACE2* expression and a significant proportion of *ACE2*^+^ *TMPRSS2*^+^ cells, we studied the kidney in more detail by looking at the integration of snRNA-seq and scATAC-seq data. Analysis of open chromatin regions revealed discrete peaks in the *ACE2* locus with the highest signal detected in a population of proximal tubule cells that also contains the highest proportion of *ACE2*-expressing cells (**Fig. 4d**). Motif analysis demonstrated that *ACE2* promoter and enhancer regions are enriched in *STAT1* and *3*, *FOXA1*, *JUNB* and several *IRF* (interferon response factor) binding sites (**Fig. 4e**). These transcription factors have important immune functions and are targets of tissue protective and innate immune responses such as those mediated by interleukin-6 (IL6), interleukin-1 (IL1) and interferons^65^. In this regard, dysregulation of both IL6 and IL1β has been implicated in the pathogenesis of severe COVID-19^66^. Thus, we investigated the co-expression of their receptors (*IL6R*, *IL1R1* and *IL1RAP*) with *ACE2* in monkey kidney, only observing good correlation with *ACE2* in proximal tubule cells for *IL6R* (**Extended Data Fig. 35b**). These observations imply a potential link between IL6, STAT transcription factors and enhanced *ACE2* expression in specific tissues such as the kidney that can either facilitate the existence of viral reservoirs or exacerbate COVID-19 disease progression due to increased viral dissemination (**Extended Data Fig. 35c)**. In addition to ACE2 and TMPRSS2, numerous other molecules have been implicated in facilitating SARS-CoV-2 binding to the cell surface or in COVID-19 pathogenesis^67,68^. Their expression or co-expression in monkey tissues, as well as other potential associations and other virus-host interactions can be explored using our NHPCA database.

### Investigation of common human traits and genetic diseases in monkey

We next assessed the effect of genetic variation linked to complex human traits and diseases by applying Genome Wide Association Studies (GWAS) to our monkey dataset. We linked human single-nucleotide polymorphisms from 163 GWAS taken from the UK Biobank to orthologous coordinates in the monkey single-cell transcriptome to calculate the enrichment of traits across the genes expressed in each cell cluster annotated in our dataset. As a general trend, we observed enriched heritability for neurological traits such as ‘schizophrenia’, ‘depression’ or ‘autism’ in clusters corresponding to neuronal and glial cells (**Fig. 5a, Extended Data Fig. 36 and Supplementary Table 5a**). Similarly, we observed enrichment of Alzheimer’s disease traits in immune cells, in line with the knowledge that immune dysfunction contributes to the pathogenesis of this disease^69^. Consistent with expectations, we also noticed enrichment of immunological-related traits (‘lymphocyte count’, ‘monocyte count’ and traits related to immune disorders) in myeloid cells and B and T lymphocytes. Likewise, blood related traits such as ‘mean sphered cell volume’ and ‘red blood cell distribution width’ were enriched in erythrocytes and bone marrow progenitor cells. Interestingly, however, we observed some unexpected trends for traits like ‘body mass index’ or ‘waste ratio’. Despite showing the expected highest enrichment in adipocytes, these trends additionally revealed an enrichment in smooth muscle cells, melanocytes and stromal cells. Similarly, type 2 diabetes and cholesterol-related traits revealed not only the expected association with hepatocytes but also with several kidney cell populations^70^. Our analysis also pointed at the enrichment of attention deficit and hyperactive disorder (ADHD) in skeletal muscle type I and type II myonuclei but not in neuronal cell types, suggesting an intriguing link between this pathology and motor abnormalities (**Fig. 5a**). In this regard for example, ocular muscle hyperactivity is an accompanying sign of ADHD and might be a major trigger for the disease rather than a consequence^71^.

**Figure 5.**
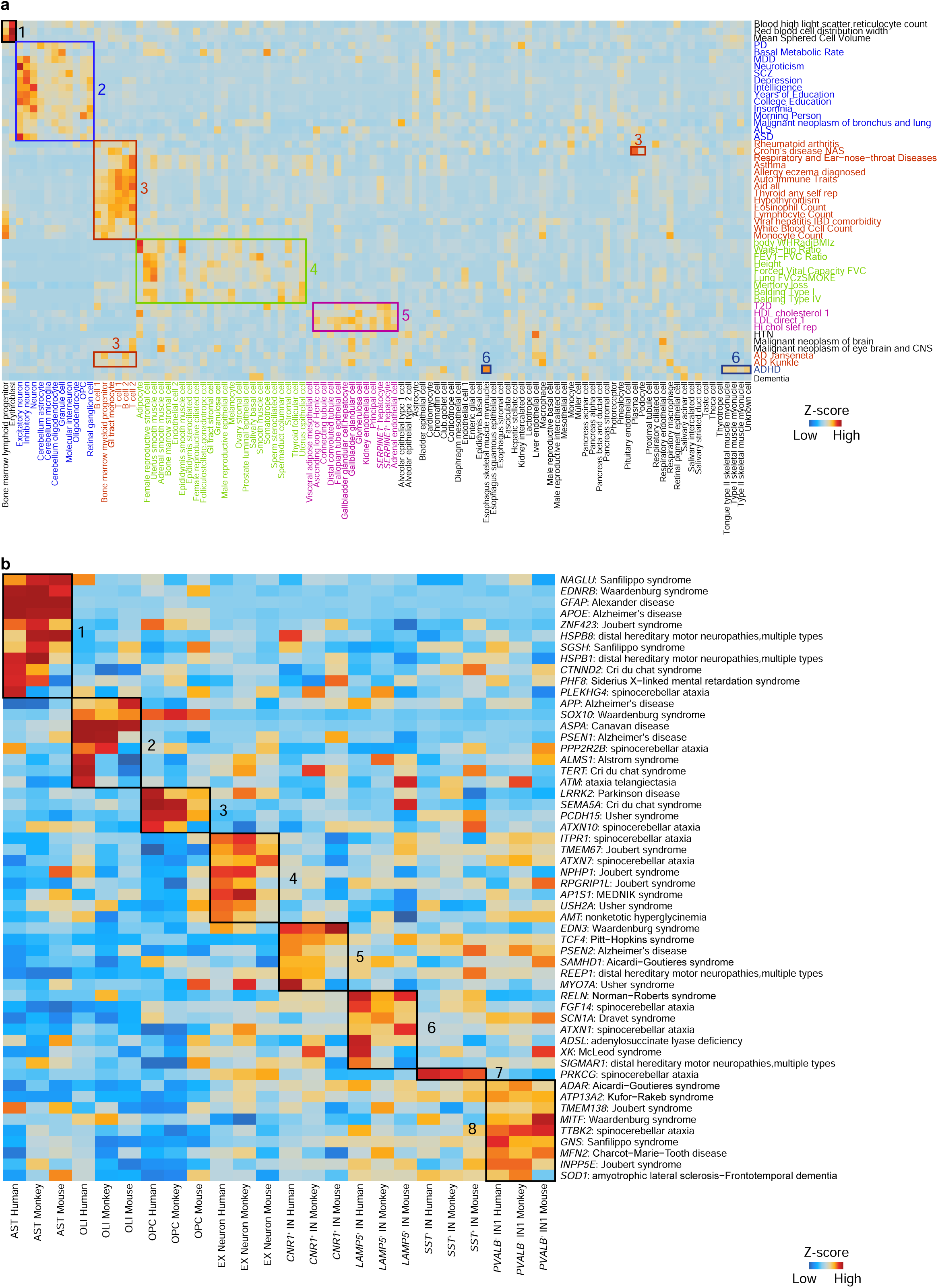
Association of monkey transcriptomic profiles with human common traits and genetic diseases. **(a)** Heatmap showing the association of selected common human traits and diseases (indicated on the right) with the cell types (indicated at the bottom) annotated in our dataset. The colored boxes indicate enriched specific patterns related to human traits/diseases subtypes. **(b)** Heatmap showing the enrichment of genetic diseases related to the central nervous system in human, monkey and mouse neocortex snRNA-seq datasets. The black boxes indicated specific patterns associated with cell types annotated in the neocortex dataset.

Besides the association of complex human traits to cell types stated above, we also generated a correlation map of mutant genes causing human genetic diseases with all cell types annotated in our monkey dataset (**Extended Data Fig. 37 and Supplementary Table 5b**). As expected, genes related to retinitis pigmentosa were specifically expressed in monkey photoreceptors, while genes related to porphyria were found associated to erythroblasts. This shows that our dataset can predict cell types that are directly affected in human genetic diseases. In addition, we compared the interspecies distribution of a panel of genes related to human neurological diseases using snRNA-seq data for mouse, monkey and human neocortex^46^. Notably, for most genes, we observed a generally higher correlation of the expression in specific cell types between human and monkey than between human and mouse (**Fig. 5b**). However, some diseases also appeared to be related to different cell types in monkey compared to human. For instance, distal neuropathy caused by mutations in *HSPB8*^72^ was enriched in *CNR1*^+^ inhibitory neurons in human while being enriched in astrocytes in monkey and mouse. Similarly, ataxia telangiectasia caused by mutations in *ATM* was mostly enriched in oligodendrocytes^73^ in human while in monkey and mouse it was enriched in *PVALB*^+^ and *LAMP5*^+^ inhibitory neurons, respectively.

Our analysis thus highlights the potential for modelling human diseases in species phylogenetically closer to humans and underlines that differences will still exist. Further scrutiny of GWAS datasets and gene mutations and wider comparisons between species will provide additional relevant observations.

## DISCUSSION

Despite the enormous potential, few NHP tissues have been profiled to date at the single-cell level and the use of different species, experimental conditions and platforms makes comparisons challenging^20,74,75^. To address this, we have generated the first version of a large single-cell transcriptomic atlas for a NHP widely used in research studies, *Macaca fascicularis*, and an expandable and interactive database (https://db.cngb.org/nhpca/) to facilitate its exploration. The current version of our atlas provides a comprehensive and integrated overview of gene expression in 106 cell types extracted from 43 tissue types. Specialized tissues such as skin, thymus, testis and some parts of the gastrointestinal tract, as well as increased cell numbers for some of the already profiled ones, will be added in future releases. Cell type identification relied on previously reported markers and gene expression profiles. Therefore, although we identified most (if not all) known cell types in these tissues, our current annotations are likely to benefit from deeper sub-clustering and further revision.

We provide a detailed description of individual tissue single-cell composition and a comparison of common cell types across all sequenced tissues. This information will be particularly valuable for understanding tissues that have either not been profiled at all at the single-cell level in human (e.g., diaphragm, tongue and salivary gland) or lack enough cell numbers (e.g., liver, gallbladder and substantia nigra), and for prediction of human disease susceptibilities. Regarding the latter, we have identified an unexpected link between ADHD and muscle function. ADHD is a polygenic and multifactorial disorder associated with hyperactivity and motor coordination abnormalities that are thought to have a neurological origin^76^. Our data support the possibility that skeletal muscle rather than the nervous system may be a direct driver of ADHD pathogenesis^77^. Similarly, as part of the analysis for virus receptors and co-receptors, we provide a comprehensive map of *ACE2*^+^/*TMPRSS2*^+^ double positive cells throughout the monkey body that may be useful to understand COVID-19 pathogenesis in human^59,61^. In particular, the link between IL6, STAT transcription factors and ACE2 expression could explain the reported positive effects of tocilizumab, a humanized monoclonal antibody against IL6R for the treatment of patients with severe COVID-19^78^. On the other hand, our study shows significant interspecies differences in cell type-specific gene expression with potentially important functional consequences. For example, the distribution of *ACE2* and *TMPRSS2* across different cell types is not identical between monkey and human and this could influence the disease course. Moreover, in the context of the survey of Wnt pathway components we have identified *LGR5*^+^ renal cells with progenitor characteristics that are seemingly absent in human and mouse based on analysis of reported datasets. This is relevant because the kidney has limited regenerative capacity in mammals^79^. During embryonic development *LGR5*^+^ cells located at the junction between the ureteric bud (source of the collecting tubule and connecting tubule) and the metanephric blastema are responsible for nephrogenesis, but they quickly disappear after birth^45^. Their persistence in adult monkey kidney suggests a higher regenerative capacity compared to other species, which if true raises the hope of activating a similar mechanism in human^80^. Similarly, *LGR5*^+^ cells in the neocortex correspond mainly to OPC in monkey and to oligodendrocytes and to a lesser extent OPC in human, whereas in mouse inhibitory neurons are more highly enriched. This finding is consistent with the knowledge that Wnt activity regulates OPC and oligodendrocyte function and differentiation^81^ but suggest interspecies differences in the mode of action. Likewise, the expression of *LGR5* in skeletal slow-twitch myofibers, and *LGR6* in the pituitary gland and heart, is intriguing. During development, Wnt activity regulates skeletal myogenesis and myofiber typing^82^, cardiomyocyte proliferation^83^ and pituitary gland growth^84^, but little is known about the adult. The functional implications of these and other related findings and the extent to which the patterns differ between monkey and other mammalian species will require further study. Finally, interspecies comparison of single-cell gene expression in neocortex highlights the problems associated with modelling neurological diseases in rodents and suggests that a cautious approach should also be taken when studying NHP. Additional comparisons with other human and mouse single-cell/nuclei datasets will provide a more comprehensive, body-wide picture of differences in disease vulnerability among the three species.

In the future, with efforts from us and scientists worldwide, the NHPCA database will be extended with additional single-cell datasets generated from disease modelling studies, spontaneously developed diseases (e.g., diabetes or cardiomyopathy) and aging. Adding other layers of single-cell -omics studies, in particular scATAC-seq and spatially resolved transcriptomics^85^ for all tissues presented here, will help characterize cell states and the interactions between different cell types more accurately. Proof of principle is the kidney scATAC-seq dataset included here. In addition, it will be important to compare our *Macaca fascicularis* atlas with datasets from other non-endangered NHP species such as *Macaca mulatta* (rhesus monkey), *Callithrix jacchus* (marmoset monkey)^86^ and *Microcebus murinus* (mouse lemur)^10,13^. Altogether, this information will be instrumental for understanding primate evolution and human disease.

**Supplementary Table 1. Description of all tissues profiled, cell types and markers used for cluster annotation**

**Supplementary Table 2. Global analysis of common cell types and tissue-specific signatures**

**Supplementary Table 3. Global distribution of LGR5, LGR6 and MKI67 expression**

**Supplementary Table 4. Analysis of the expression of common virus and SARS-Cov-2 receptors**

**Supplementary Table 5. Correlation of GWAS traits and human genetic diseases with monkey cell types**

## METHODS

### Ethics statement

This study was approved by the Institutional Review Board on Ethics Committee of BGI (permit no. BGI-IRB19125).

### Collection of monkey tissues

A total of three females and three males, approximately 6-year-old, cynomolgus monkeys were obtained from Huazhen Laboratory Animal Breeding Centre and Hubei Topgene Biotechnology (Guangzhou, China). Animals were anesthetized with ketamine hydrochloride (10 mg/kg) and sodium pantabarbital (40 mg/kg) injection before being euthanized by exsanguination. Tissues were isolated and placed on the ice-cold board for dissection. A total of 43 whole tissues were isolated: abdominal wall, adrenal gland, aorta and carotid arteries, bladder, bone marrow, bronchia, cerebellum, colon, diaphragm, duodenum, epididymis, esophagus, fallopian tube, gallbladder, heart, kidney, liver, lung, lymph node, neocortex, ovary, pancreas, PBMC, pigmentary epithelium choroid plexus, pineal gland, pituitary gland, prostate, retina, salivary gland, spermaduct, spinal cord, spleen, stomach, subcutaneous adipose tissue, substantia nigra, thyroid gland, tongue, tonsil, trachea, uterus, vagina and visceral adipose tissue. Each tissue (except for bone marrow, peripheral blood and tissues on which enzymatic digestion was performed) was cut into 5-10 pieces of roughly 50-200 mg each. Samples were transferred to cryogenic vials (Corning, #430488), then quickly frozen in liquid nitrogen and finally stored until nuclear extraction was performed. PBMC and bone marrow cells were isolated from heparinized venous blood using a Lymphoprep^TM^ medium (STEMCELL Technologies, #07851) according to standard density gradient centrifugation methods. Cells from those two tissues were resuspended in 90% FBS, 10% DMSO (Sigma Aldrich, #D2650) freezing media and frozen using a Nalgene® Mr. Frosty® Cryo 1°C Freezing Container (Thermo Fisher Scientific, #5100-0001) in a -80°C freezer for 24 hours before being transferred to liquid nitrogen for long-term storage.

### Single-nucleus/cell suspension preparation

Single nucleus isolation was performed as described previously^87^. Briefly, tissues were thawed, minced and transferred to a 1 ml Dounce homogenizer (TIANDZ) with 1 ml of homogenization buffer A containing 250 mM sucrose (Ambion), 10 mg/ml BSA (Ambion), 5mM MgCl_2_ (Ambion), 0.12 U/μl RNasin Plus (Promega, #N2115), 0.12 U/μl RNasein (Promega, #N2115) and 1x Protease Inhibitor (Roche, #11697498001). Tissues were kept in an ice box and homogenized by 25-50 strokes of the loose pestle (Pestle A) after which the mixture was filtered using a 100 µm cell strainer in to a 1.5 ml tube (Eppendorf). The mixture was then transferred to a clean 1 ml dounce homogenizer to which 750 ul of buffer A containing 1% Igepal (Sigma, #CA630) was added and the tissue was further homogenized by 25 strokes of the tight pestle (Pestle B). After this, the mixture was filtered through a 40 µm strainer in a 1.5 ml tube and centrifuged at 500 g for five minutes at 4°C to pellet nuclei. At this stage, the pellet was resuspended in 1 ml of buffer B containing 320 mM Sucrose, 10 mg/ml BSA, 3 mM CaCl_2_, 2 mM MgAc_2_, 0.1 mM EDTA, 10 mM Tris-HCl, 1 mM DTT, 1x Protease Inhibitor and 0.12 U/μl RNasein. This was followed by a centrifugation at 500 g for five minutes at 4°C to pellet nuclei. Nuclei were then resuspended with cell resuspension buffer at a concentration of 1,000 nuclei/μl for single-nucleus library preparation. Cells from lymph node, spleen, duodenum, stomach and colon were obtained from fresh tissues by enzymatic digestion. Briefly, tissues were rinsed in PBS, minced into small pieces by mechanical dissociation and incubated for 1 hour in 10 ml of DS-LT buffer (0.2 mg/ml CaCl_2_, 5 μM MgCl_2_, 0.2% BSA and 0.2 mg/ml Liberase in HBSS) at 37°C. After this, the tissue digestion was stopped by addition of 3 ml of FBS, followed by filtration through a 100 µm cell strainer and centrifugation for 5 minutes at 500 g at 4°C. Samples were then filtered through a 40 µm cell strainer and centrifuged for five minutes at 500 g at 4°C. Pellets were then resuspended in cell resuspension buffer at 1,000 cells/μl for single-cell library preparation.

### Single-cell/single-nucleus RNA-seq (sc/snRNA-seq)

DNBelab C Series Single-Cell Library Prep Set was utilized as previously described^14^. In brief, single-nucleus/cell suspensions were used for droplet generation, emulsion breakage, beads collection, reverse transcription and cDNA amplification to generate barcoded libraries. Indexed sc/snRNA-seq libraries were constructed according to the manufacturer’s protocol. The concentration of sc/snRNA-seq sequencing libraries was quantified by Qubit™ ssDNA Assay Kit (Thermo Fisher Scientific, #Q10212). The resulting libraries were sequenced using a DIPSEQ T1 or DIPSEQ T7 sequencers at the China National GeneBank (Shenzhen, China).

### Single-cell ATAC-seq (scATAC-seq)

ScATAC-seq libraries were prepared using DNBelab C Series Single-Cell ATAC Library Prep Set^14^. DNA nanoballs were loaded into the patterned Nano arrays and sequenced on a BGISEQ-500 sequencer using the following read length: 50 bp for read 1, 76 bp for read 2, inclusive of 50 bp insert DNA, 10 bp cell barcode 1, 6 bp constant sequence and 10 bp cell barcode 2.

### Immunofluorescence

Staining of monkey neocortex sample was conducted following standard protocol^88^. In brief, paraffin embedded sections were deparaffinized, incubated with primary antibodies for PDGFRα (Cell Signaling #3174S) and LGR5 (Abcam #ab273092) overnight at 4℃, followed by an incubation with a secondary antibody (Alexa Fluor 488 and Cy3, Jackson ImmunoResearch) for 30 minutes at room temperature. Slides were mounted with Slowfade Mountant+DAPI (Life Technologies, #S36964) and sealed.

### Single-molecule fluorescence *in situ* hybridization (smFISH)

SmFISH in monkey kidney, diaphragm and heart tissues was performed using RNAScope Fluorescent Multiplex and RNAScope Multiplex Fluorescent v2 (Advanced Cell Diagnostics) according to manufacturer’s instructions. The following alterations were added: the thickness of paraffin section was adjusted to 5 μm and target retrieval boiling time was adjusted to 15 minutes while the incubation time of Protease plus at 40℃ was adjusted to 30 minutes. RNA smFISH probes used: *LGR5* (C1), *LGR6* (C2), *MYH7* (C2).

### Sc/snRNA-seq data processing

Raw sequencing reads from DIPSEQ-T1 or DIPSEQ-T7 were filtered and demultiplexed using PISA (version 0.2) (https://github.com/shiquan/PISA). Reads were aligned to Macaca_fascicularis_5.0 genome using STAR (version 2.7.4a)^89^ and sorted by sambamba (version 0.7.0)^90^. For tissues sequenced with scRNA-seq, reads were aligned to the exon of mRNA as normal. For tissues sequenced with snRNA-seq, a custom ‘pre-mRNA’ reference was created for alignment of count reads to introns as well as to exons because of large amount of unspliced pre-mRNA and mature mRNA in the cell nucleus. Thus, each gene’s transcripts in snRNA-seq was counted out by including exon and intron reads together^91^. In the end, cell/nucleus versus gene UMI count matrix was generated with PISA.

### Doublet removal

For each library, we performed doublet removal using DoubletFinder^92^. DoubletFinder first averages the transcriptional profile of randomly chosen cell pairs to create pseudo doublets and then predicts doublets according to each real cell’s similarity in gene expression to the pseudo doublets. The doublet removal was performed according to the default parameter of DoubletFinder and the top 5% of cells most similar to the “pseudo doublets” were excluded.

### Cell clustering and identification of cell types

Clustering analysis of the complete cynomolgus monkey tissue dataset was performed using Scanpy (version 1.6.0)^93^ in a Python environment (version 3.6). Parameters used in each function were manually curated to portray the optimal clustering of cells. In the preprocessing, cells or nuclei were filtered based on the criteria of expressing a minimum of 500 genes and genes expressed by at least three cells or nuclei were kept for the following analysis. In addition, cells or nuclei with more than 10% mitochondrial gene counts were removed. Filtered data were ln (counts per million (CPM)/100 + 1) transformed. 3,000 highly variable genes were selected according to their average expression and dispersion. The number of UMI and the percentage of mitochondrial genes were regressed out and each gene was scaled by default options. Dimension reduction starts with principal component analysis and the number of principal components used for UMAP depended on the importance of embeddings. Louvain method is then used to detect subgroups of cells. Distinguishing differential genes among clusters were ranked (Benjamini-Hochberg, Wilcoxon rank-sum test). Cell types were manually and iteratively assigned based on overlap of literature, curated and statistically ranked genes. Each tissue dataset was portrayed using the Seurat package (version 3.2.2)^94^ in R environment (version 3.6). Data from different replicates were integrated following the standard integrated pipeline by default parameters for filtering, data normalization, dimensionality reduction, clustering and gene differential expression analysis. Finally, we annotated each cell type by extensive literature reading and searching for the specific gene expression patterns.

### Differentially expressed gene (DEG) and gene ontology (GO) term enrichment analysis

In the global clustering, we performed DEG analysis using the sc.pl.rank_genes_groups function in Scanpy (V1.6.0). In other studies, we used the FindMarker or FindAllMarker function in the Seurat R package (V3.2.2). Analysis of DEG comparing specific populations was performed by calculating the fold-change of the mean expression level of genes between the selected populations. DEG were defined as those with a fold-change > 2 and adjusted *P* < 0.01. GO enrichment analysis was performed using the CompareCluster function fun = “enrichGO”, pvalueCutoff = 0.1, pAdjustMethod = “BH”, OrgDb = org.Hs.eg.db,ontBP”) of ChIPseeker R package (v.1.22.1)^95^. Only GO terms with adjusted *P* < 0.05 were retained.

### Analysis of inter-species differences

For tissue inter-species analysis, in order to get more accurate comparisons, we specifically chose three tissues with snRNA-seq data, namely kidney, neocortex and heart, and processed the raw sequencing data using our pipeline described below in the ‘Sc/snRNA-seq data processing’ section. Kidney^43,44^, neocortex^46^ and heart^49,50^ data were downloaded from NCBI Gene expression omnibus (human kidney: GSE121862, mouse kidney: GSE119531, human neocortex: GSE97942, human heart: ERP123138, mouse heart: E-MTAB-7869). For each tissue we preprocessed the UMI matrix of the three species following three steps: 1. only orthologs genes among three species were kept. 2. only genes expressed in at least one cell in one species were kept. 3. the gene names of the human and mouse UMI matrix were converted into orthologs in *Macaca fascicularis*. After preprocessing, the UMI matrices of the three species were integrated together and the clustering was performed following the standard integrated pipeline using Seurat (V3.2.2) with the addition of one additional criterion for which only cells expressing more than 500 genes were kept. Also, we downsampled the cells of human and macaque neocortex to 10,000 to get a better clustering result. The Seurat clusters were then annotated into different cell types using cell type-specific markers as described above. In addition, for the comparison presented in Extended Data Figure 35 we retrieved the publicly available single-cell data for gallbladder, liver and lung from GEO GSE134355^3^, GEO GSE108098^6^ and GSE124395^96^, respectively. Data from the three species were integrated, clustered and annotated in the same way as described.

### Common cell analysis

We performed common cell analysis for 7 cell types across all the 43 tissues, those being stromal cells, macrophages/microglia, endothelial cells, smooth muscle cells, skeletal muscle cells, mesothelial cells and adipocytes. For each cell type, we extracted those cells from all tissues in our dataset according to the cell type annotation presented in Extended Data Figure 7-10. For the downstream analysis, we excluded cell types with numbers lower than 200. Data from different replicates were integrated following the standard integrated pipeline using Seurat (V3.2.2).

### Single-cell trajectory analysis

Cell lineage trajectory was inferred using Monocle2^97^ following the tutorial. We used the “differentialGeneTest” function to derive DEG from each cluster and genes with *q* < 0.01 were used to order the cells in a pseudotime analysis. After the cell trajectories was constructed, DDRtree was used to visualize it in a two-dimensional space.

### Cell-cell interaction network

To assess the cellular crosstalk between different cell types in each tissue, we used CellPhoneDB, a public repository of ligand-receptor interactions^98^. Cell type-specific receptor-ligand interactions between cell types were identified based on specific expression of a receptor by one cell type and a ligand by another cell type. The interaction score refers to the mean total of all individual ligand-receptor partner average expression values in the corresponding interacting pairs of cell types. For this analysis, we applied a statistical method to ensure that only receptors or ligands expressed in more than 10% of the cells in the given cluster were considered. The total mean of the individual partner average expression values in the corresponding interacting pairs of cell types was calculated. For the cell-cell interaction analysis in Extended Data Figure 27-31, we plot the figure based on the indicated genes related to *LGR5* and *LGR6*.

### Association of GWAS summary data of human diseases and traits with monkey cell types

To test for the enrichment of human diseases and traits in DEG for each cluster of cells based on global clustering, we applied LD (linkage disequilibrium) score regression analysis. For this, we only considered genes with an adjusted *P* < 0.05 and fold-change > 2 in the tested cell types. For accuracy, cell types identified in a number lower than 100 were excluded from this analysis. We converted the gene coordinates of *Macaca fascicularis* into hg19 genome coordinates by downloading from Ensembl the homologous gene list. Single nucleotide polymorphisms located in gene regions of the most specific genes in each cell type were added to the baseline model independently for each cell type (one file for each cell type). We then selected the coefficient *z*-score *P* value as a measure of the association of the cell type with the traits. All plots show the −log_10_ *P* value of partitioned LDscore regression.

### ScATAC-seq data processing, clustering and cell type identification

Raw sequencing reads from BGISEQ-500 were filtered and demultiplexed using PISA (version 0.2) (https://github.com/shiquan/PISA). The fragment file of each scATAC-seq library was used for downstream analysis. TSS (transcription start site) enrichment score and fragment number of each nuclei was calculated by using ArchR software^99^. Nuclei with TSS enrichment score lower than five and fragment number lower than 1,000 were removed. Then, we calculated the doublet score with *addDoubletScores* function in ArchR package and filtered doublets by *filterDoublets* function with parameter filterRatio = 2. ScATAC–seq clustering analysis was performed using ArchR software by first identifying a robust set of peak regions followed by iterative LSI (latent semantic indexing) clustering. Briefly, we created 500 bp windows tiled across the genome and determined whether each cell was accessible within each window. Next, we performed an LSI dimensionality reduction on these windows with *addIterativeLSI* function in ArchR packages. We then performed Seurat clustering (*FindClusters*) on the LSI dimensions at resolutions of 0.8. Anchors between scATAC-seq and sc/snRNA-seq datasets were identified and used to transfer cell type labels identified from the sc/snRNA-seq data. We embedded the data by the TransferData function of Seurat (version 3.2.2).

### Transcription factor motif enrichment analysis

To predict the motif footprint in peaks within the *ACE2* promoter and enhancer sequences, we extracted genome sequences in the peak region with Seqkit (version 0.7.0)^100^. The sequences were imported into R and were matched with all *Homo sapiens* motifs form JASPAR2018 using matchMotifs function in motifmatchr packages version 1.8.0 with default parameter.

### Data availability

All raw data have been deposited to CNGB Nucleotide Sequence Archive (accession code: CNP0001469; https://db.cngb.org/cnsa/project/CNP0001469/reviewlink/).

## ACKNOWLEDGMENTS

We would like to thank Wei Liu and Liangzhi Xu from Huazhen Laboratory Animal Breeding Centre for helping in the collection of monkey tissues, Dahai Zhu and Hu Li from Bioland Laboratory (Guangzhou Regenerative Medicine and Health Guangdong Laboratory) for technical help, Guoji Guo and Huiyu Sun from Zhejiang University for providing analytical advice, Guoyi Dong and Chao Liu from BGI research, Xiao Zhang, Peng Li, and Chen Qi from the Guangzhou Institutes of Biomedicine and Health for experimental advice. This work was supported by the Shenzhen Key Laboratory of Single-Cell Omics (ZDSYS20190902093613831), Shenzhen Bay Laboratory (SZBL2019062801012) and the Guangdong Provincial Key Laboratory of Genome Read and Write (2017B030301011). Additionally, Longqi Liu was supported by the National Natural Science Foundation of China (31900466) and Shenzhen Basic Research Project for Excellent Young Scholars (2020251518), Miguel A. Esteban’s was supported by a Changbai Mountain Scholar award (419020201252), the Strategic Priority Research Program of the Chinese Academy of Sciences (XDA16030502), and a Chinese Academy of Sciences-Japan Society for the Promotion of Science joint research project (GJHZ2093), Giacomo Volpe was supported by a Chinese Academy of Sciences President’s International Fellowship for Foreign Experts (2020FSB0002), and Mingyuan Liu was supported by the National Key Research and Development Program of China (2021YFC2600200).

## AUTHOR CONTRIBUTIONS

L.H., Y.H, X.X., M.A.E. and L.L. conceived the idea; Y.H, X.X., M.A.E. and L.L. supervised the work; L.H., X.W., Y.Y., M.A.E. and L.L designed the experiments; L.H., X.W., G.V., Y.Y., X.Zhang., P.F., P.G., X.L., F.Y., S.S., G.L., J.A., Y.Lei., Y. Lai, M.C., C.W., X.G., S.L. and J.M. collected tissue samples; C.L., G.V., Z.W., Y.Y., X. Zhang., P.F., Q.D., Ya. Liu, Y.Huang, H.L., B.W., M.C., J.X., M.W., C. Wang, Y.Z., Y. Yu, H.Z., Y.W. and S.X. performed the experiments. L.H., X.W., G.V., Z.Z., X. Zou, T.P., Y. Lai, L.W., Q.S., H.Y., Yang Liu, D.X., F.H., Z.Zhu and C.Ward performed data analysis. L.H., X.W., C.L., G.V., Z.Z., X.Zou, Z.Wang, T.P., Y.Yang, J.L. and L.L. prepared the figures. H.Y., X.F.W., F.C., T.Y., W.D. and J.C. prepared the website. Zong.W., Z.P., C.W.W., B.Q, A.S., J.I., L.F., Yan Liu, Z.L., X. Liu, H. Zhang, M.L., Q.S., P.M., N.B., P.M.C., Y.G., J.M., M.U., T.T., S.L., H.Y. and J.W. provided relevant advice and reviewed the manuscript. L.H., G.V., M.A.E. and L.L. wrote the manuscript with input from all authors. All other authors contributed to the work. All authors read and approved the manuscript for submission.

---

**Extended Data Figure 1.**
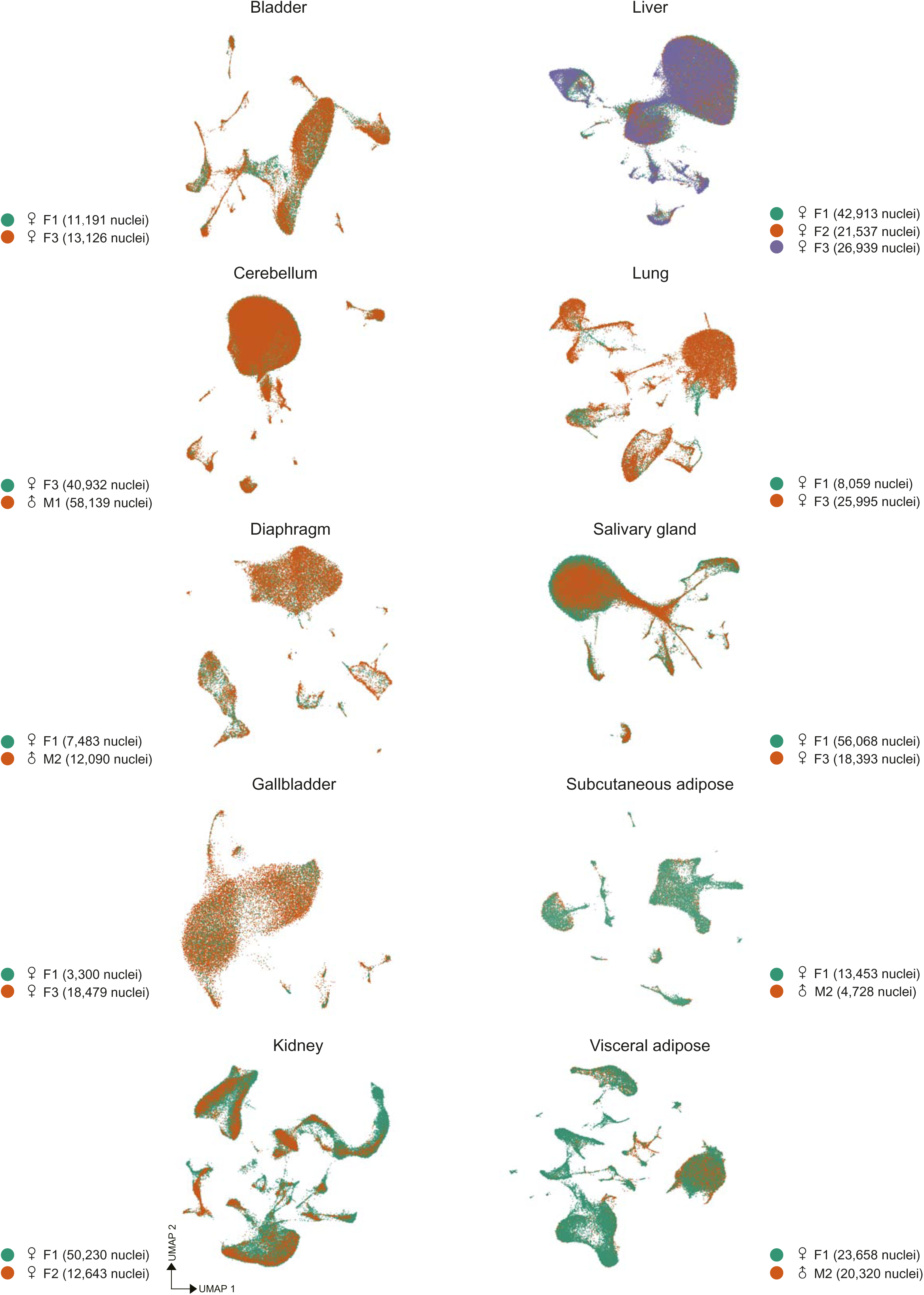
Quality control analysis of gender and individual effect. UMAP visualization of single-cell profiles for selected tissues to calculate the batch effect between tissues from different individuals and genders. Two individuals were analyzed for bladder (F1 and F3), cerebellum (F3 and M1), diaphragm (F1 and M2), gallbladder (F1 and F3), kidney (F1 and F2), lung (F1 and F3), salivary gland (F1 and F3), subcutaneous (F1 and M2) and visceral adipose (F1 and M2) tissues, and three for liver (F1, F2 and F3).

**Extended Data Figure 2.**
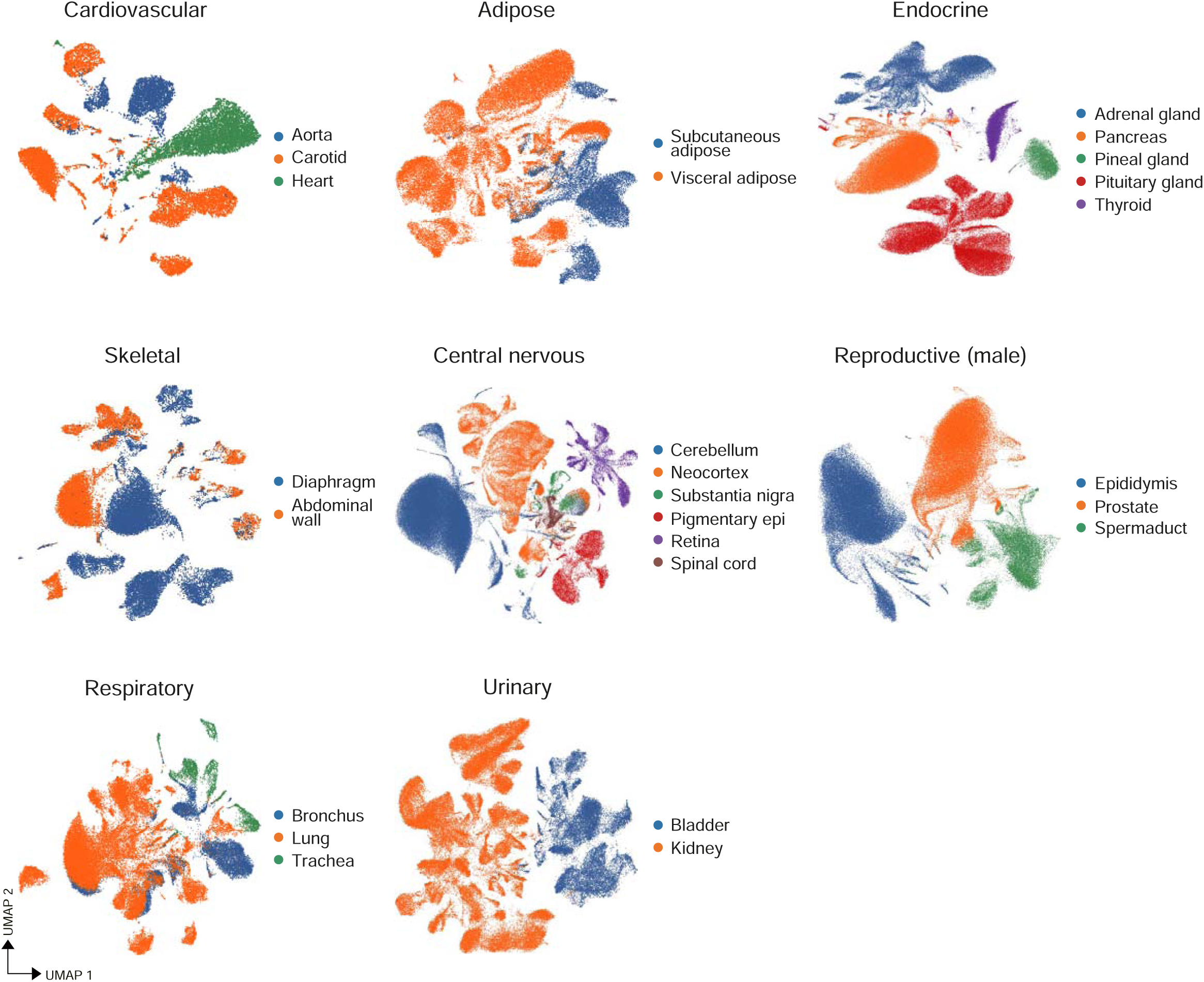
Global clustering of different systems. UMAP visualization of cell clusters in selected tissues grouped by system: cardiovascular (aorta, carotid and heart), endocrine (adrenal, pancreas, pineal, pituitary and thyroid glands), skeletal (abdominal wall and diaphragm), central nervous (cerebellum, neocortex, pigmentary epithelium choroid plexus, retina and spinal cord), respiratory (bronchus, lung and trachea) and urinary (bladder and kidney). Adipose tissues (subcutaneous and visceral) are also shown grouped. Clusters shown in every plot are colored by tissue. Abbreviation: pigmentary epi, pigmentary epithelium and choroid plexus.

**Extended Data Figure 3.**
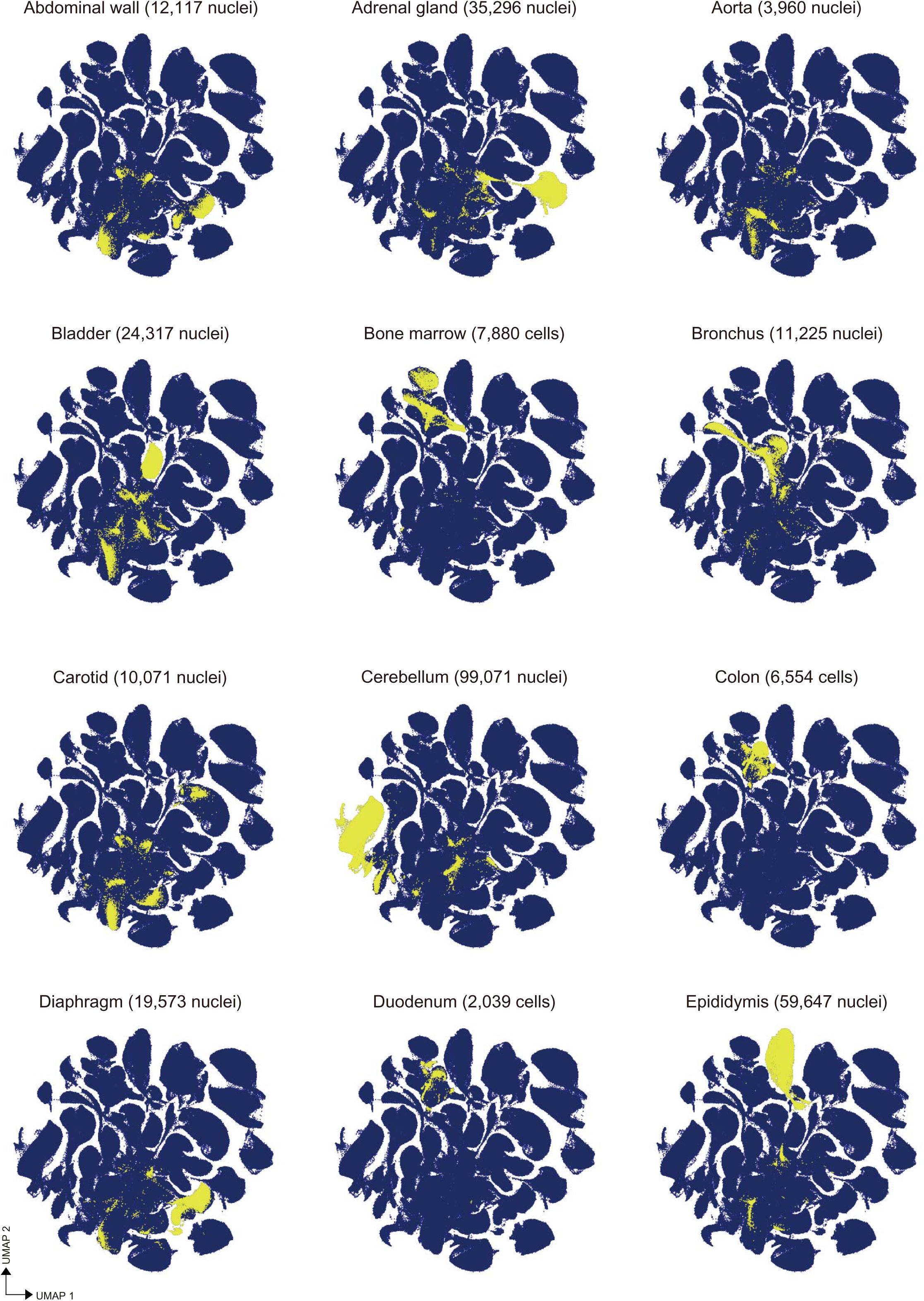
Global profiling of individual monkey tissues – 1. UMAP projection of the global clustering indicating the distribution of all single cells (highlighted in yellow) from individual tissues for abdominal wall, adrenal gland, aorta, bladder, bone marrow, bronchus, carotid, cerebellum, colon, diaphragm, duodenum and epididymis.

**Extended Data Figure 4.**
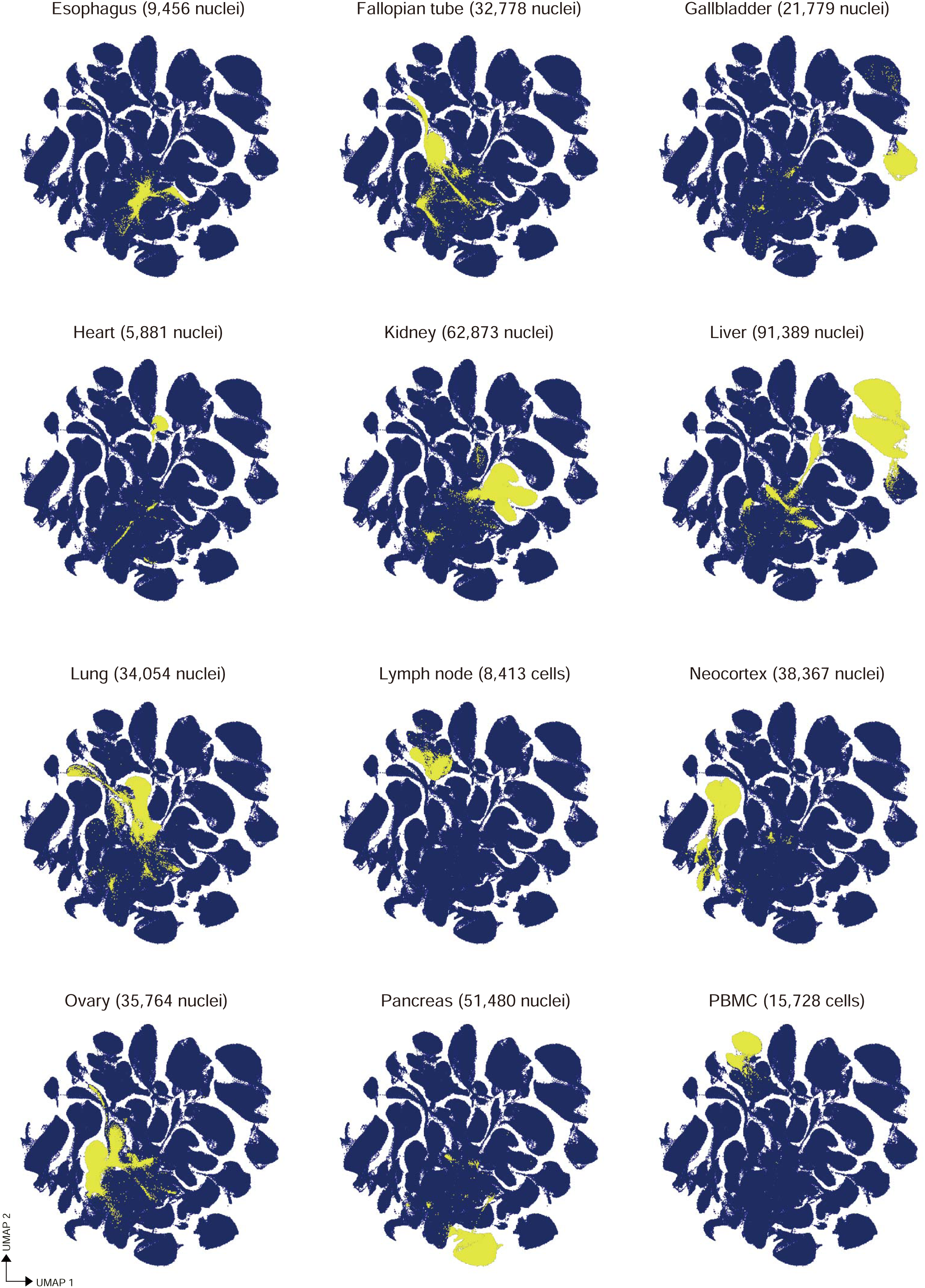
Global profiling of individual monkey tissues – 2. UMAP projection of the global clustering indicating the distribution of all single cells (highlighted in yellow) from individual tissues for esophagus, fallopian tube, gallbladder, heart, kidney, liver, lung, lymph node, neocortex, ovary, pancreas and PBMC.

**Extended Data Figure 5.**
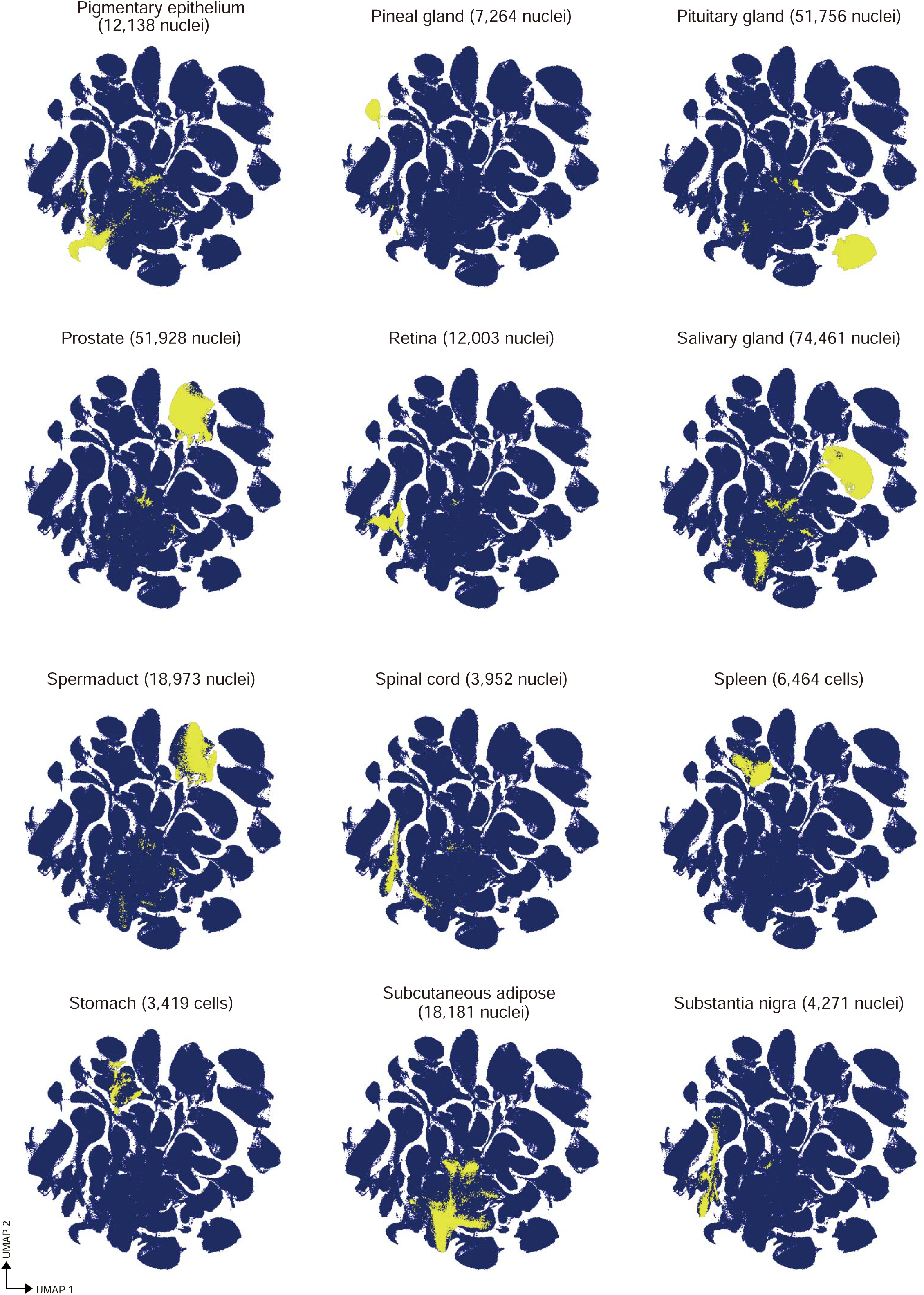
Global profiling of individual monkey tissues – 3. UMAP projection of the global clustering indicating the distribution of all single cells (highlighted in yellow) from individual tissues for pigmentary epithelium choroid plexus, pineal gland, pituitary gland, prostate, retina, salivary gland, spermaduct, spinal cord, spleen, stomach, subcutaneous adipose tissue and substantia nigra.

**Extended Data Figure 6.**
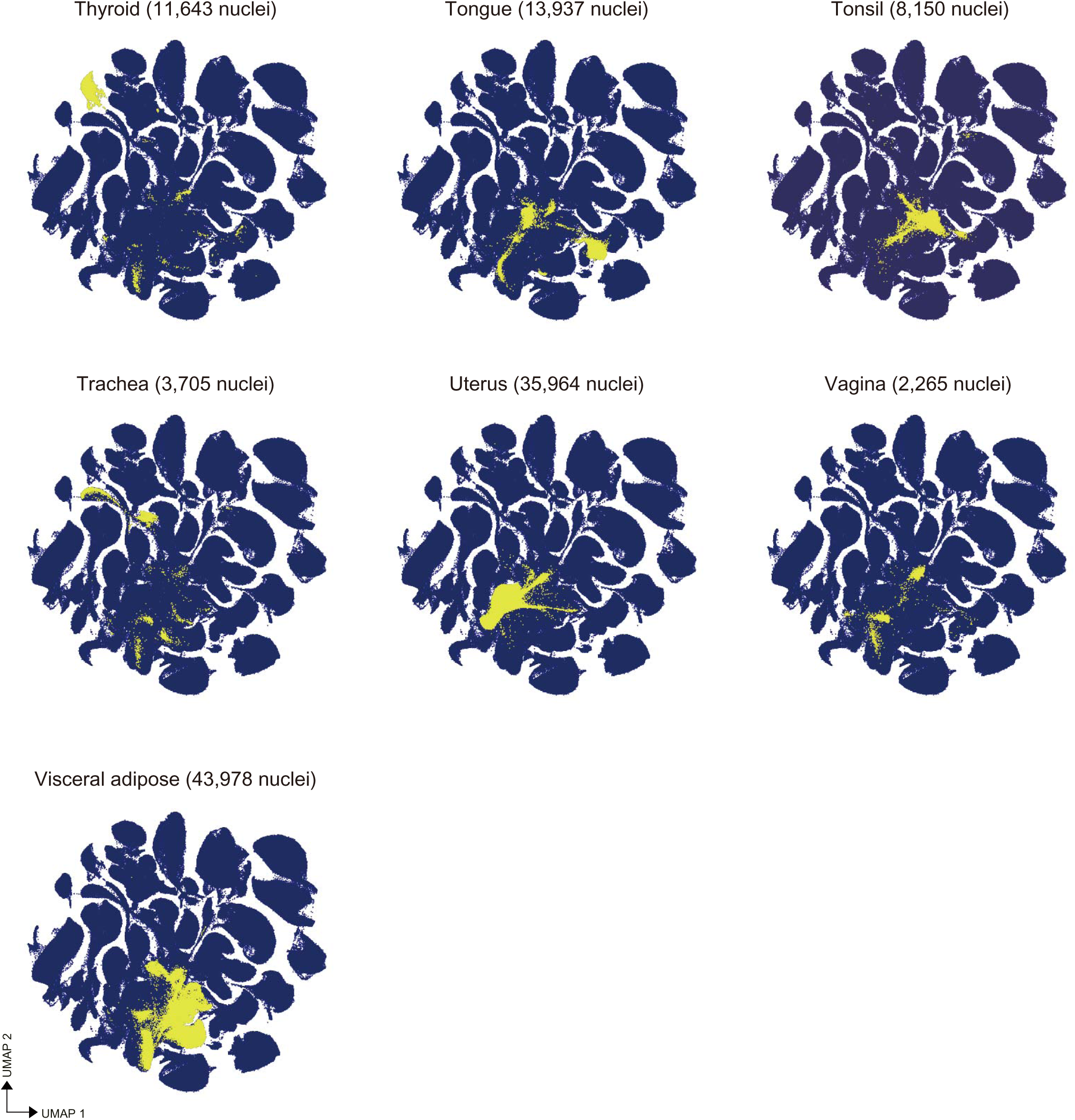
Global profiling of individual monkey tissues – 4. UMAP projection of the global clustering indicating the distribution of all single cells (highlighted in yellow) from individual tissues for thyroid, tongue, tonsil, trachea, uterus, vagina and visceral adipose tissue.

**Extended Data Figure 7.**
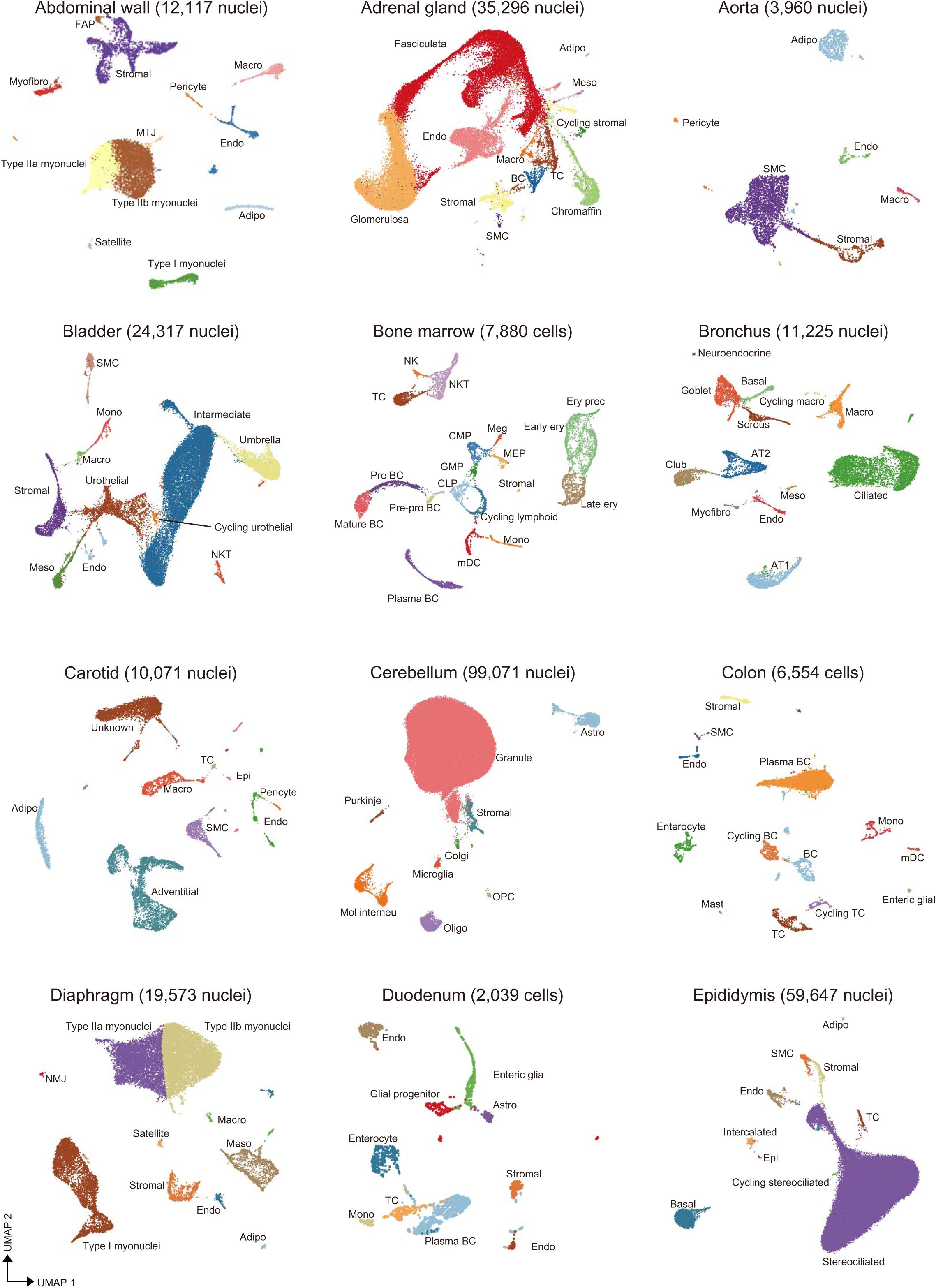
Cluster annotations – 1. UMAP visualization of cell clusters in the abdominal wall, adrenal gland, aorta, bladder, bone marrow, bronchus, carotid, cerebellum, colon, diaphragm, duodenum and epididymis. The name of the population in each cluster and the total number of cells profiled for every tissue are indicated in every plot. Abbreviations: Adipo, adipocytes; Astro, astrocytes; AT1, alveolar type 1 cells; AT2, alveolar type 2 cells; BC, B cells; CLP, common lymphoid progenitors; CMP, common myeloid progenitors; Endo, endothelial cells; Epi, epithelial cells; Ery, erythroblasts; FAP, fibroadipogenic progenitors; GMP, granulocyte monocyte progenitors; Macro, macrophages; mDC, myeloid derived dendritic cells; MEP, megakaryocyte erythrocyte progenitors; Meso, mesothelial cells; Mol interneu, molecular interneurons; Mono, monocytes; MTJ, myotendinous junction; Myofibro, myofibroblasts; NK, natural killers; NKT, natural killer T cells; NMJ, neuromuscular junction; Oligo, oligodendrocytes; OPC, oligodendrocyte progenitor cells; SMC, smooth muscle cells; TC, T cells.

**Extended Data Figure 8.**
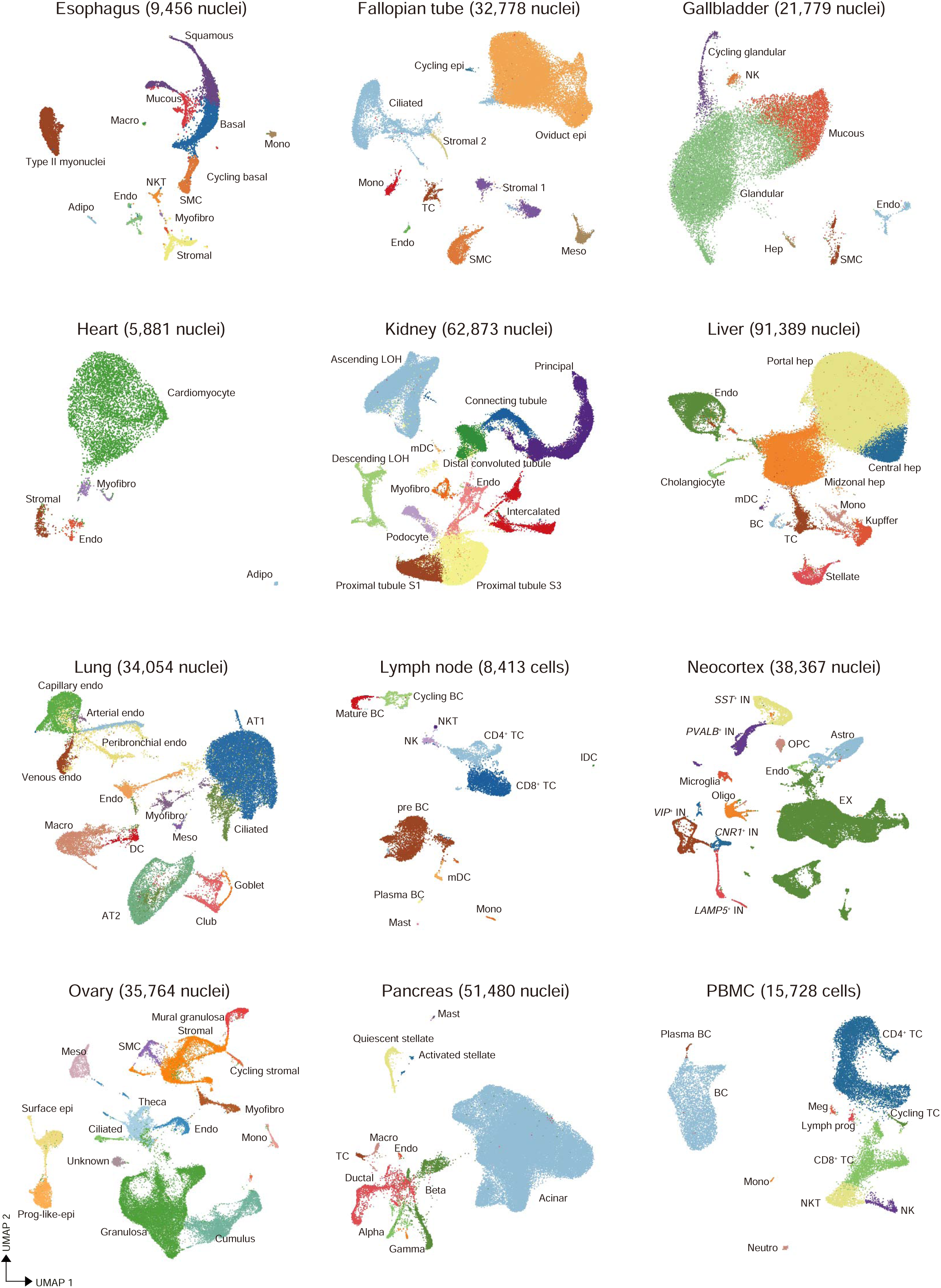
Cluster annotations – 2. UMAP visualization of cell clusters in the esophagus, fallopian tube, gallbladder, heart, kidney, liver, lung, lymph node, neocortex, ovary, pancreas and PBMC. The name of the population in each cluster and the total number of cells profiled for every tissue are indicated in every plot. Abbreviations: Adipo, adipocytes; Astro, astrocytes; AT1, alveolar type 1 cells; AT2, alveolar type 2 cells; BC, B cells; Endo, endothelial cells; Epi, epithelial cells; EX, excitatory neurons; Hep, hepatocytes; IN, inhibitory neurons; lDC, lymphoid derived dendritic cells; LOH, loop of Henle cells; Lymph prog, lymphoid progenitors; Macro, macrophages; mDC, myeloid derived dendritic cells; Meg, megakaryocytes; Meso, mesothelial cells; Mono, monocytes; Myofibro, myofibroblasts; NK, natural killers; NKT, natural killer T cells; NMJ, neuromuscular junction; Oligo, oligodendrocytes; OPC, oligodendrocyte progenitor cells; Prog-like epi, progenitor-like epithelial cells; SMC, smooth muscle cells; TC, T cells.

**Extended Data Figure 9.**
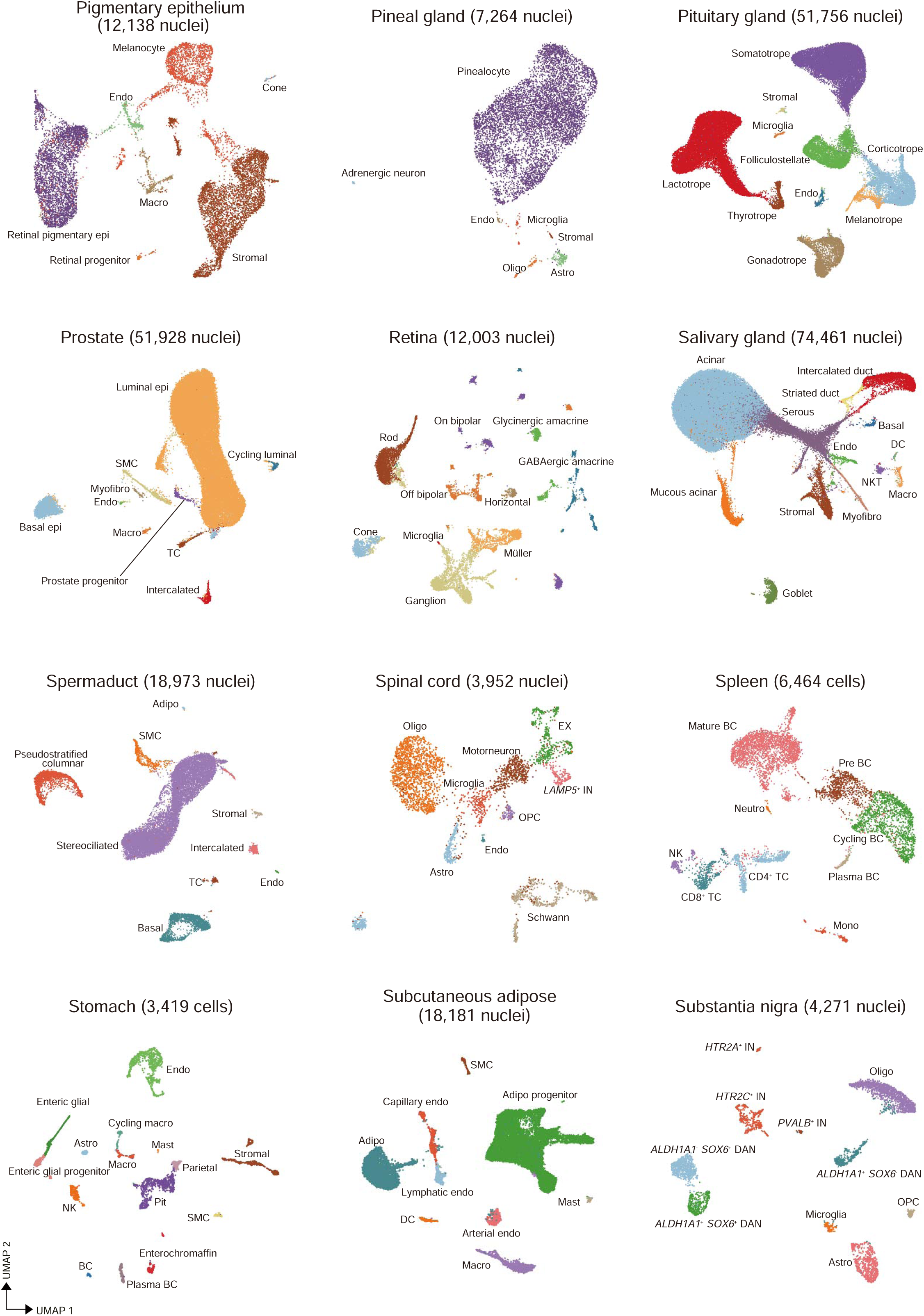
Cluster annotations – 3. UMAP visualization of cell clusters in the pigmentary epithelium choroid plexus, pineal gland, pituitary gland, prostate, retina, salivary gland, spermaduct, spinal cord, spleen, stomach, subcutaneous adipose tissue and substantia nigra. The name of the population in each cluster and the total number of cells profiled for every tissue are indicated in every plot. Abbreviations: Adipo, adipocytes; Astro, astrocytes; BC, B cells; DAN, dopaminergic neurons; DC, conventional dendritic cells; Endo, endothelial cells; Epi, epithelial cells; EX, excitatory neurons; IN, inhibitory neurons; Macro, macrophages; Mono, monocytes; Myofibro, myofibroblasts; Neutro, neutrophils; NK, natural killers; NKT, natural killer T cells; Oligo, oligodendrocytes; OPC, oligodendrocyte progenitor cells; SMC, smooth muscle cells; TC, T cells.

**Extended Data Figure 10.**
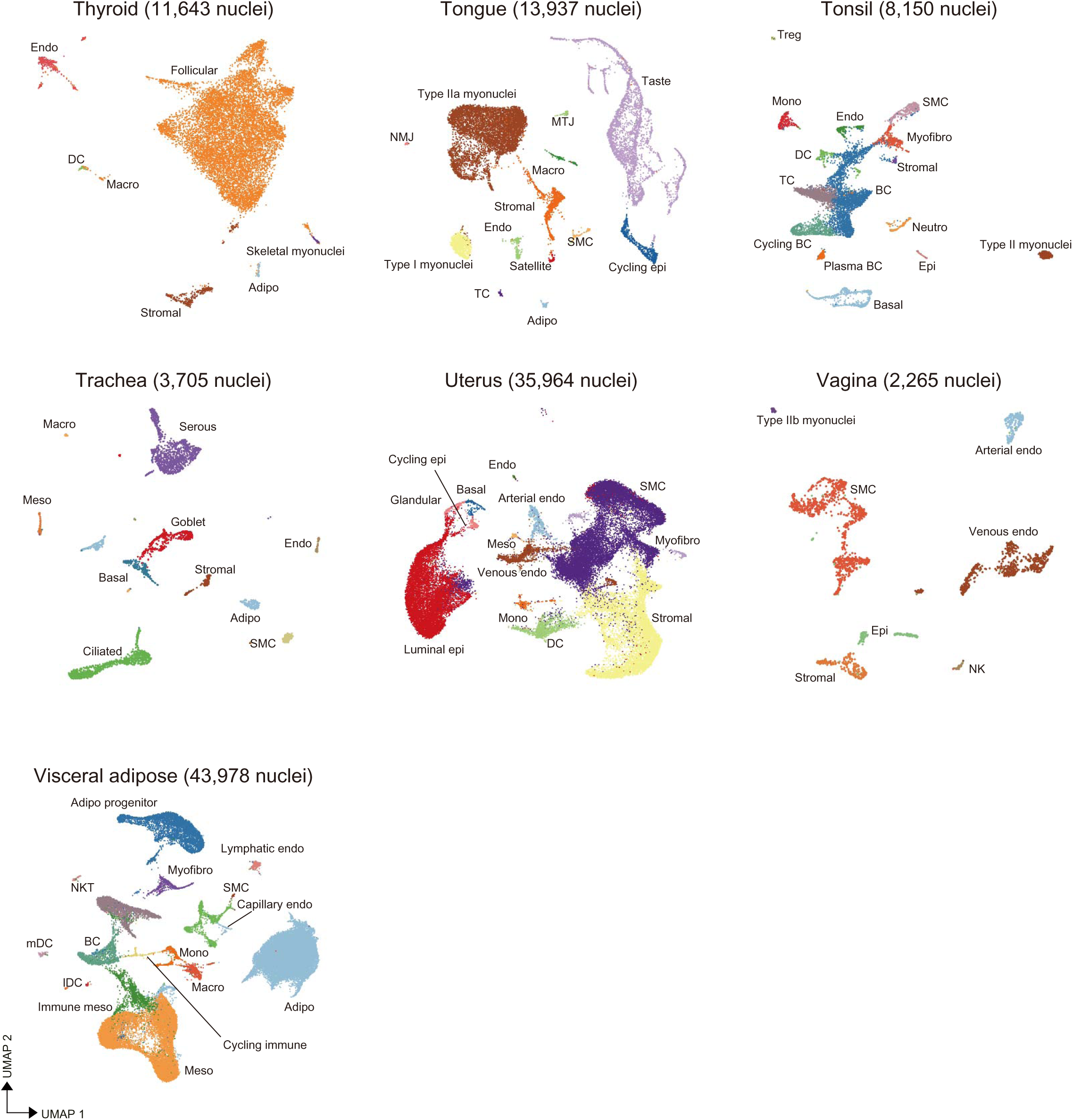
Cluster annotations – 4. UMAP visualization of cell clusters in the thyroid, tongue, tonsil, trachea, uterus, vagina and visceral adipose tissue. The name of the population in each cluster and the total number of cells profiled for every tissue are indicated in every plot. Abbreviations: Adipo, adipocytes; BC, B cells; Endo, endothelial cells; lDC, lymphoid derived dendritic cells; LOH, loop of Henle; Macro, macrophages; mDC, myeloid derived dendritic cells; Meso, mesothelial cells; Mono, monocytes; NK, natural killers; NMJ, neuromuscular junction; SMC, smooth muscle cells; TC, T cells.

**Extended Data Figure 11.**
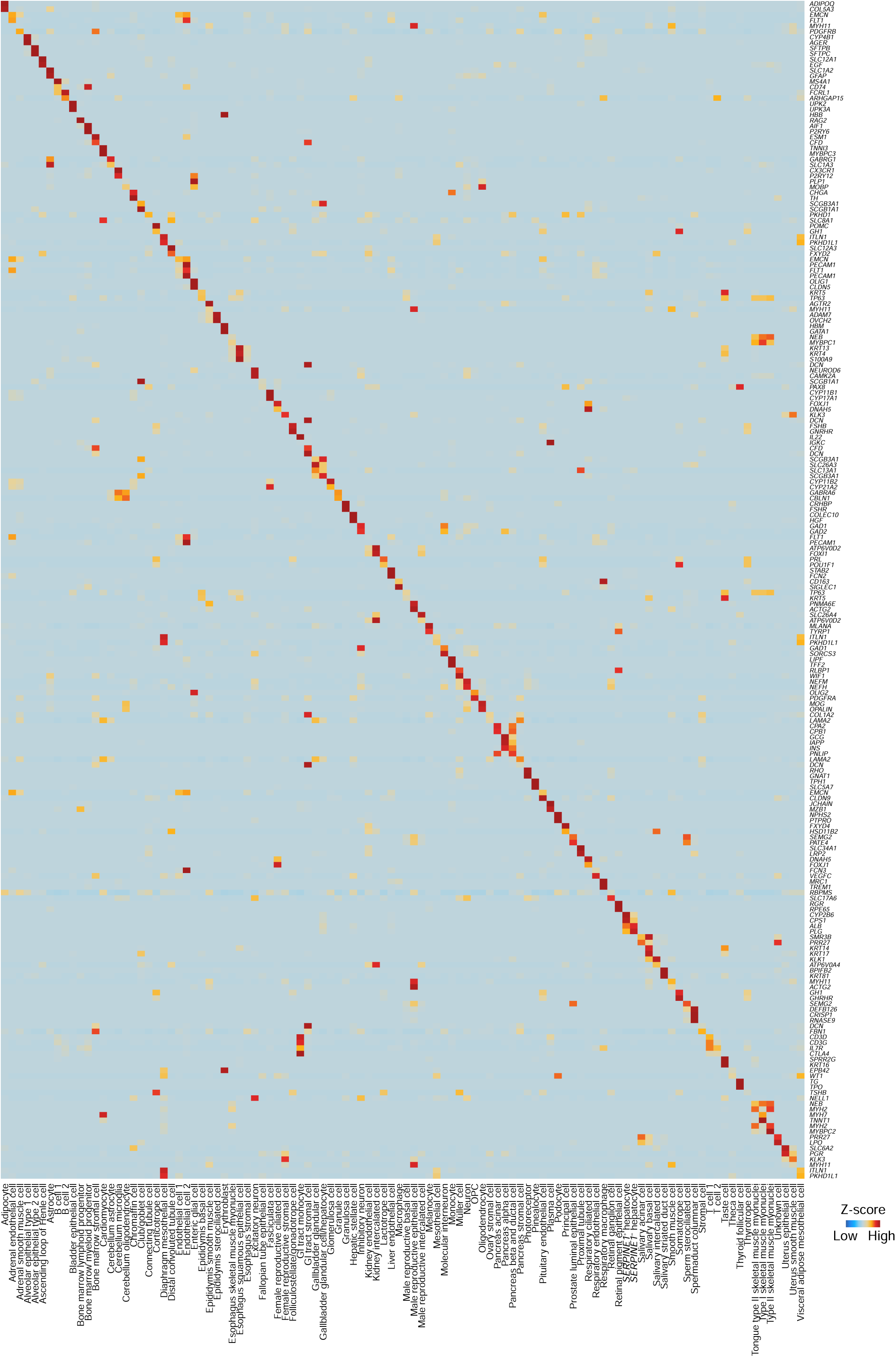
Selected markers for cell cluster annotations. Heatmap showing the expression of the marker genes used to manually annotate all cell clusters identified in every tissue of this dataset.

**Extended Data Figure 12.**
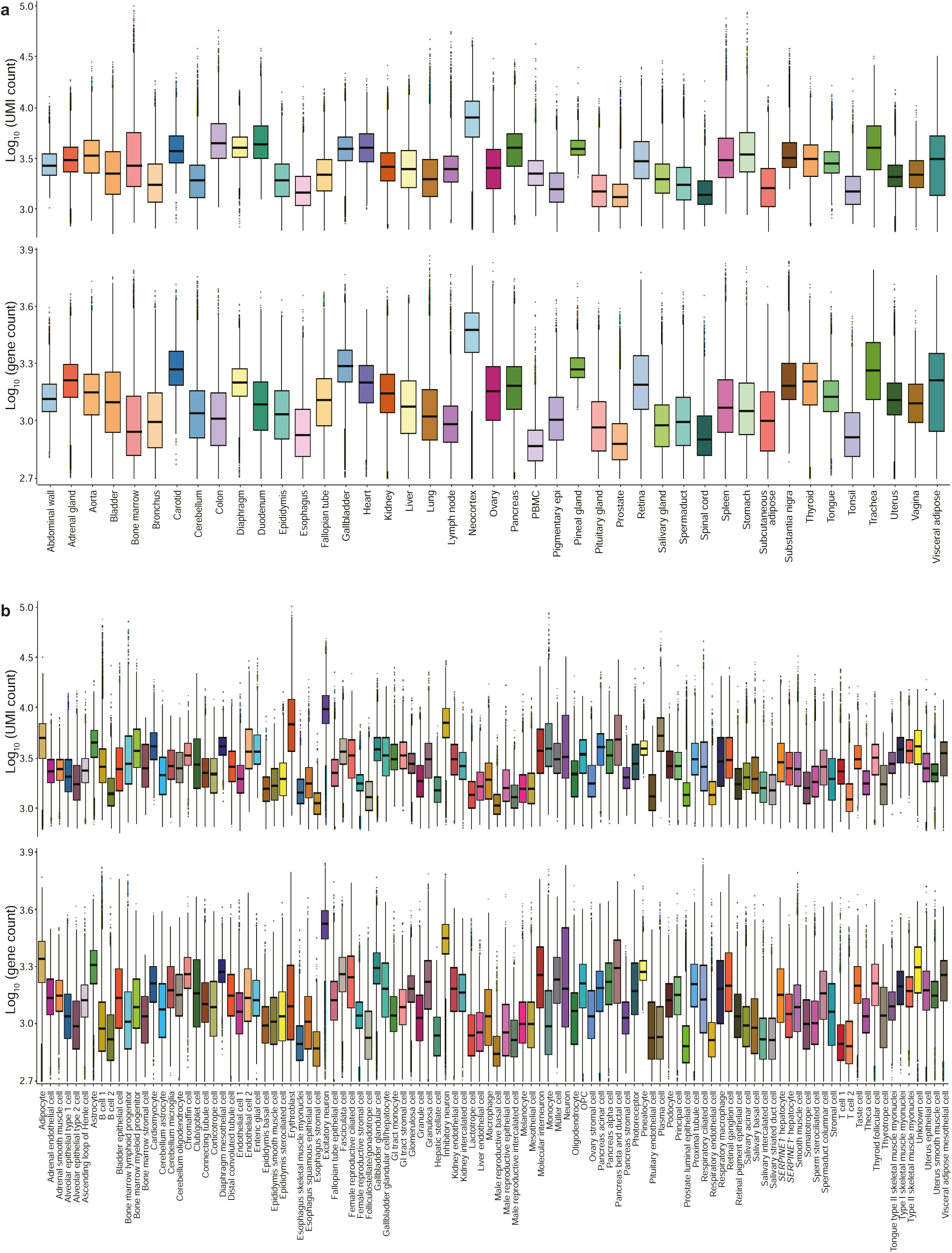
UMI and gene numbers of the sequenced tissues and annotated cell types. **(a)** Boxplot indicating the number of UMI (top) and genes (bottom) in each tissue of the dataset. **(b)** Boxplot indicating the number of UMI (top) and genes (bottom) detected in each of the major annotated cell types shown in Figure 1c.

**Extended Data Figure 13.**
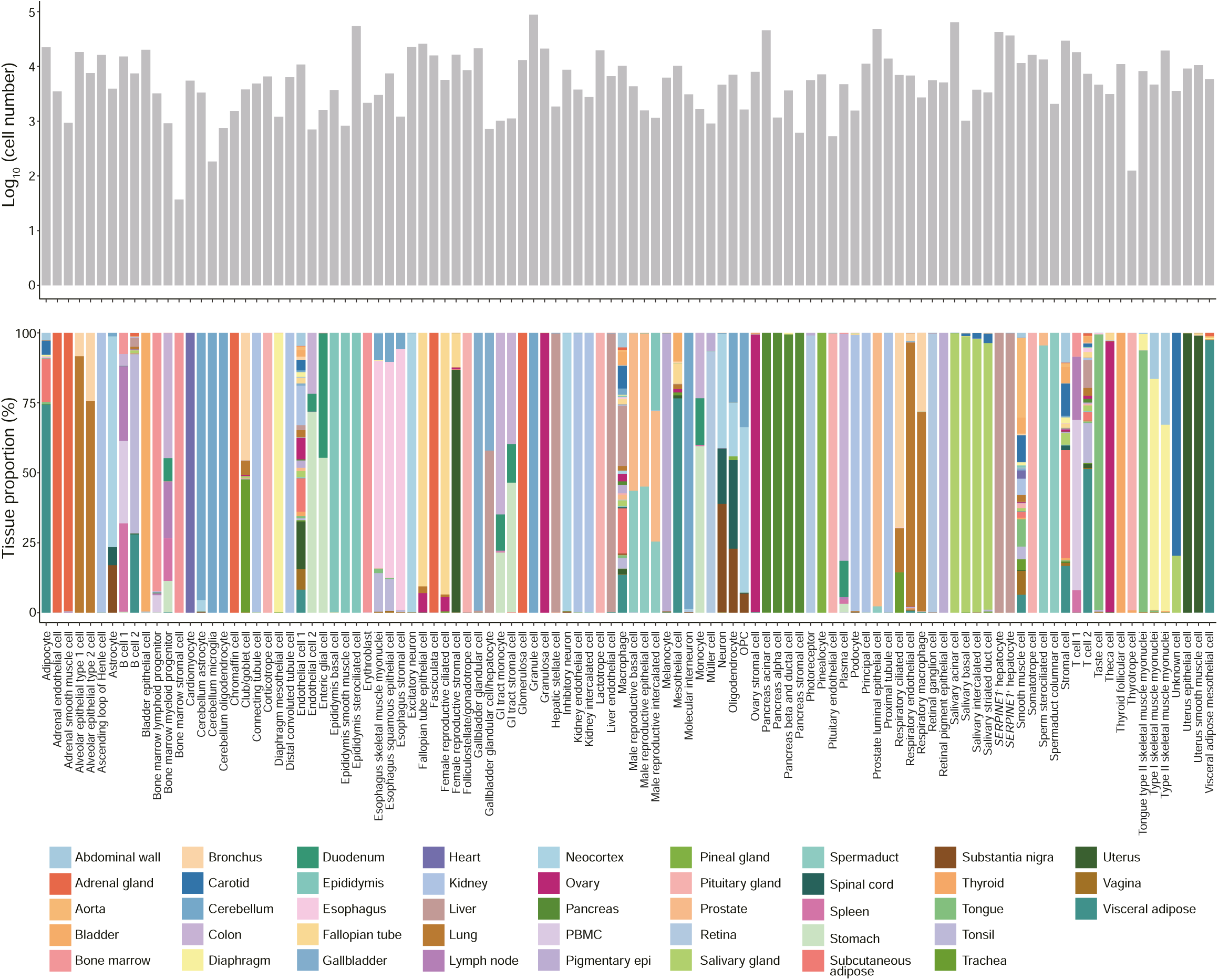
Cell numbers and proportions among the sequenced tissues. Bar plot representation of the number of cells analyzed for each cell type described in main Figure 1c. The stacked bar plot at the bottom indicates the ratio of each cell type detected in every tissue.

**Extended Data Figure 14.**
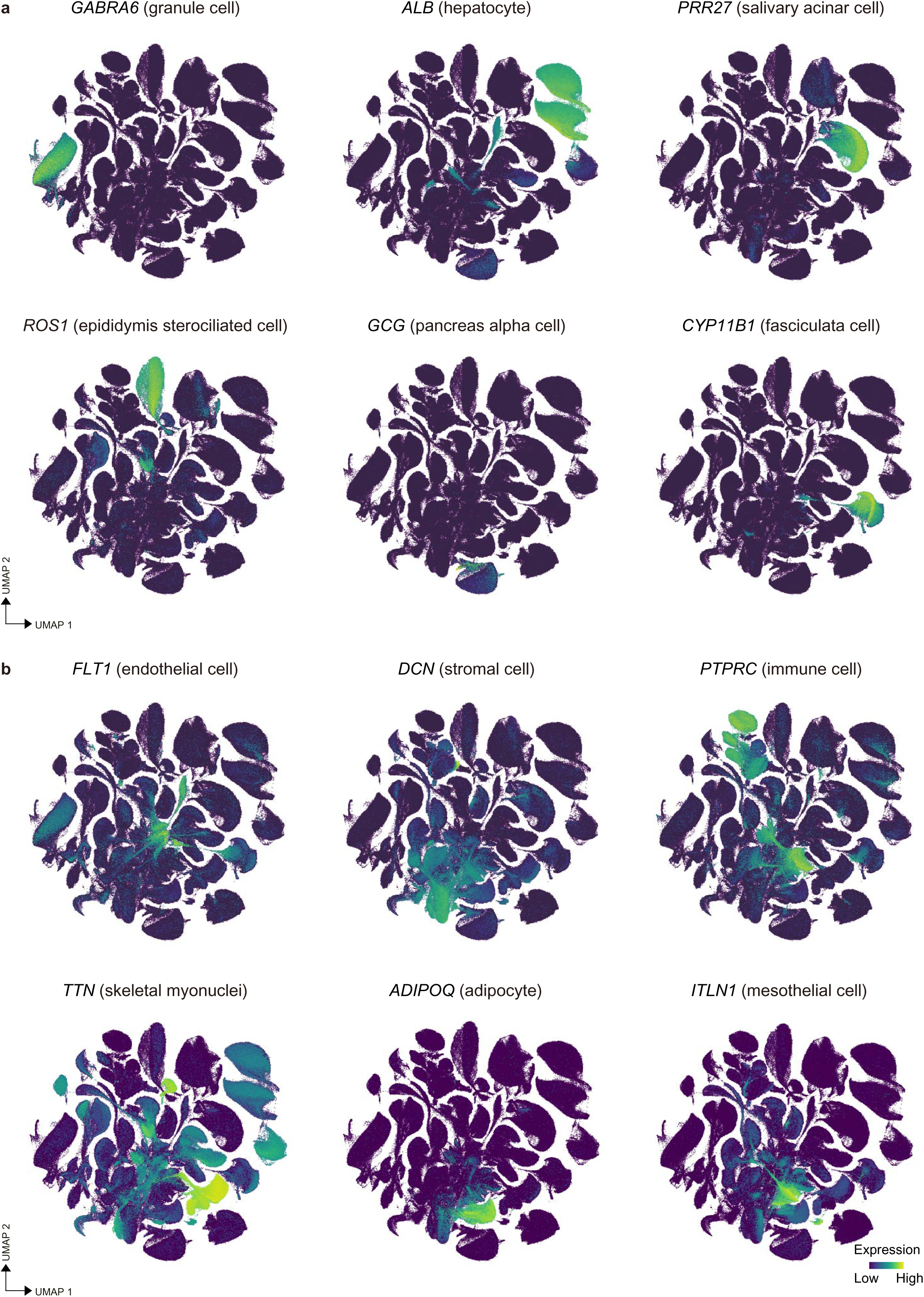
Unique and shared cell populations. **(a)** UMAP projection of the global clustering showing the expression of specific markers for cerebellum granule cells (*GABRA6*), hepatocytes (*ALB*), salivary gland acinar cells (*PRR27*), epididymis stereociliated cells (*ROS1*), pancreatic alpha cells (*GCG*) and fasciculata cells of the adrenal gland (*CYP11A1*). **(b)** UMAP projection of the global clustering showing the expression of pan-markers of endothelial (*FLT1*), stromal (*DCN*), immune (*PTPRC*), skeletal myonuclei (*TTN*), adipocytes (*ADIPOQ*) and mesothelial cells (*ITLN1*) that are shared across tissues.

**Extended Data Figure 15.**
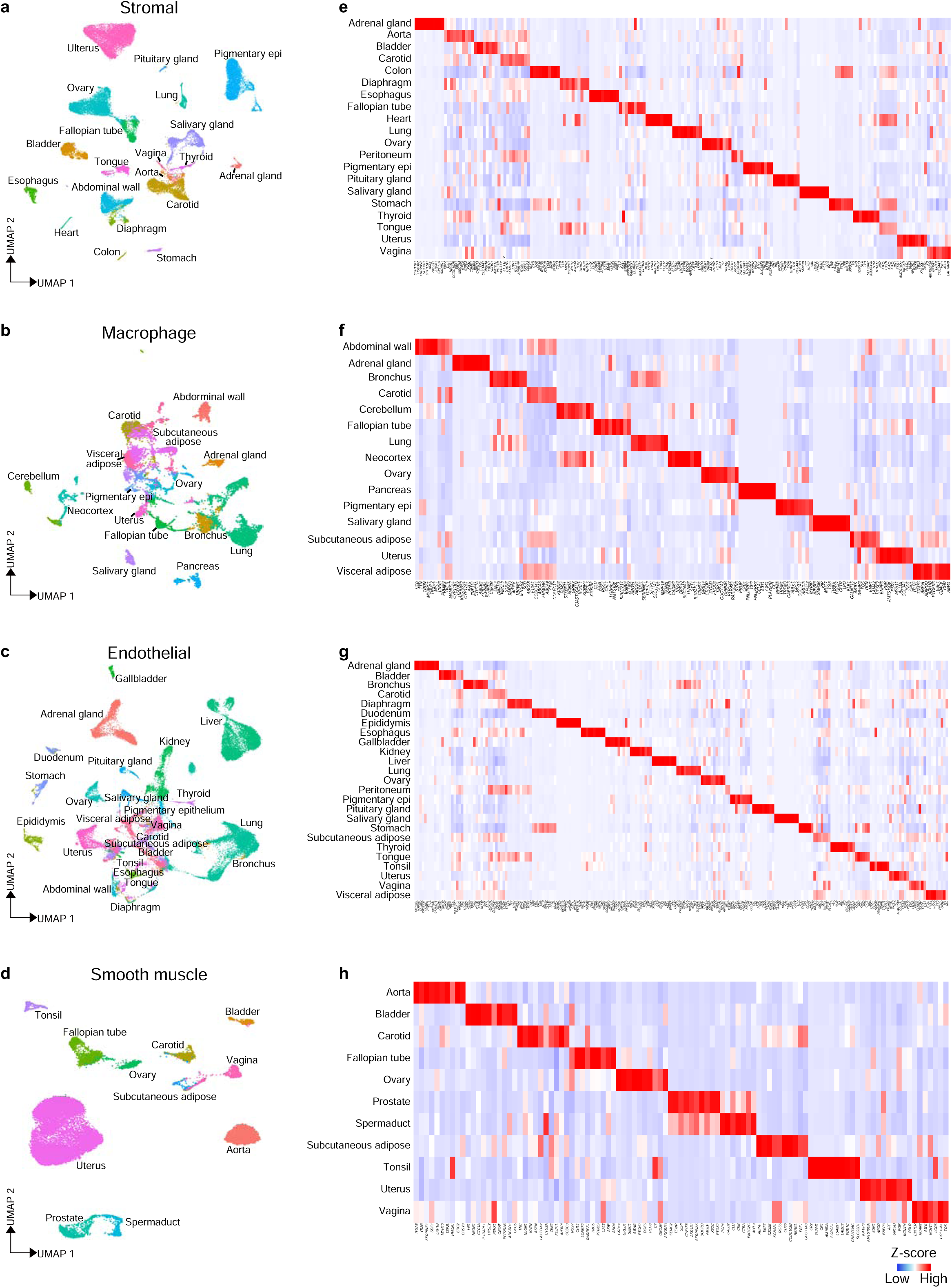
Global analysis of common cell types. UMAP visualization of (**a**) stromal cells (n = 35,415), **(b)** macrophages (n = 10,929), **(c)** endothelial cells (n = 37,640) and **(d)** smooth muscle cells (n = 24,175) from all analyzed monkey tissues. Tissues with low numbers of the selected cell types were excluded. Cell clusters are colored by tissue. The heatmap on the right shows tissue-specific DEG for **(e)** stromal cells, **(f)** macrophages, **(g)** endothelial cells and **(h)** smooth muscle cells.

**Extended Data Figure 16.**
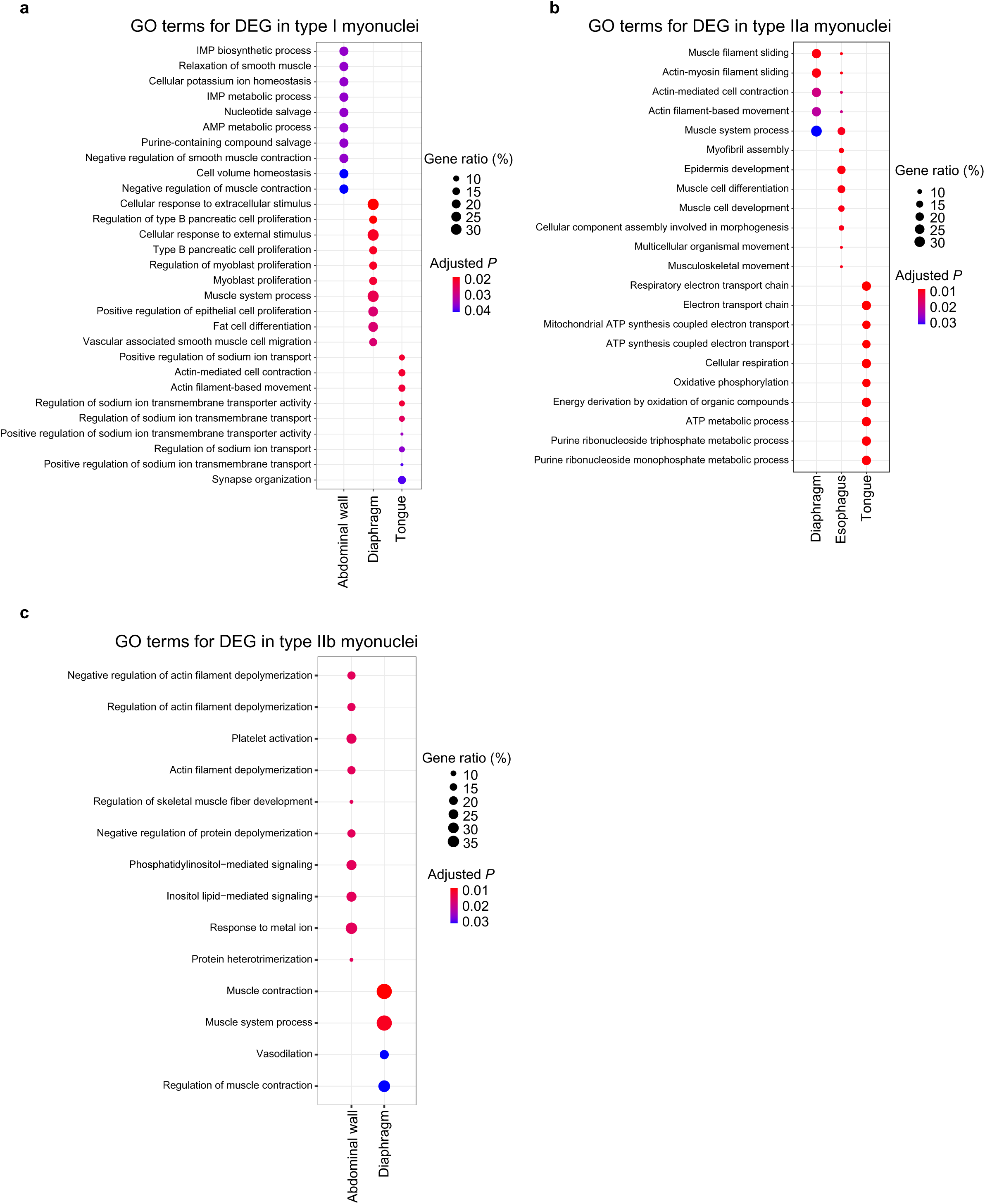
Analysis of skeletal myonuclei molecular signatures. **(a)** Bubble plot indicating tissue-specific enriched GO terms in type I myonuclei from abdominal wall, diaphragm and tongue. **(b)** Bubble plot indicating tissue-specific enriched GO terms in type IIa myonuclei from diaphragm, esophagus and tongue. **(c)** Bubble plot indicating tissue-specific enriched GO terms in type IIb myonuclei from abdominal wall and diaphragm.

**Extended Data Figure 17.**
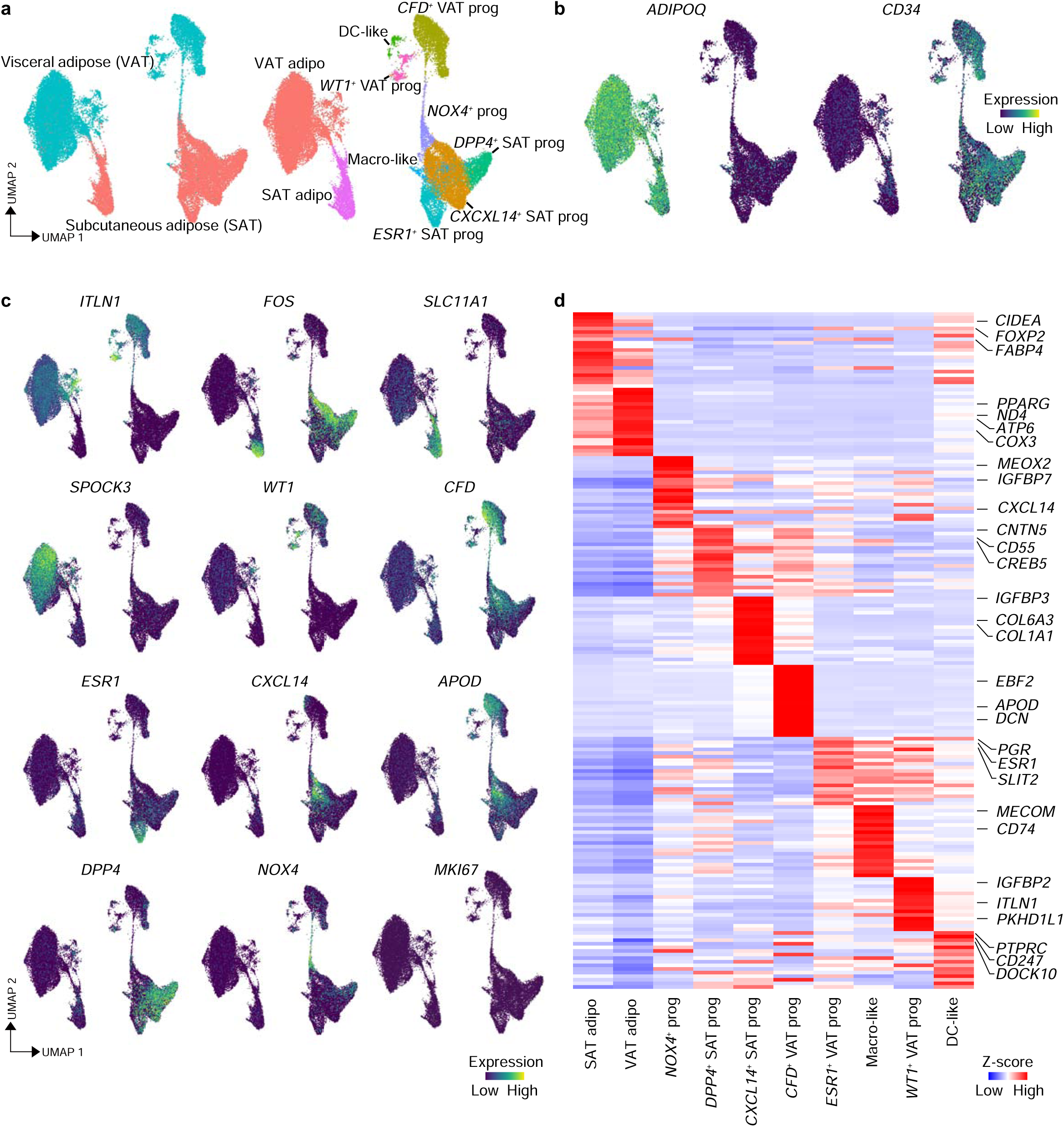
Global analysis of adipocyte populations. **(a)** UMAP visualization of mature adipocyte and adipocyte progenitors from visceral (VAT) and subcutaneous (SAT) adipose tissues. Data were grouped together and re-clustered either by tissue type (on the left) or by cell type (on the right). **(b)** UMAP visualization of specific markers for mature adipocytes (*ADIPOQ*) or adipocyte progenitors (*CD34*). **(c)** UMAP visualization of markers for tissue-specific (*ITLN1* and *FOS*), cell-type specific (*SLC11A1*, *SPOCK3*, *WT1, ESR1, CXCL14, APOD, CFD, DPP4* and *NOX4*) or cycling markers (*MKI67*). **(d)** Heatmap indicating the DEG in all clusters identified in **a**.

**Extended Data Figure 18.**
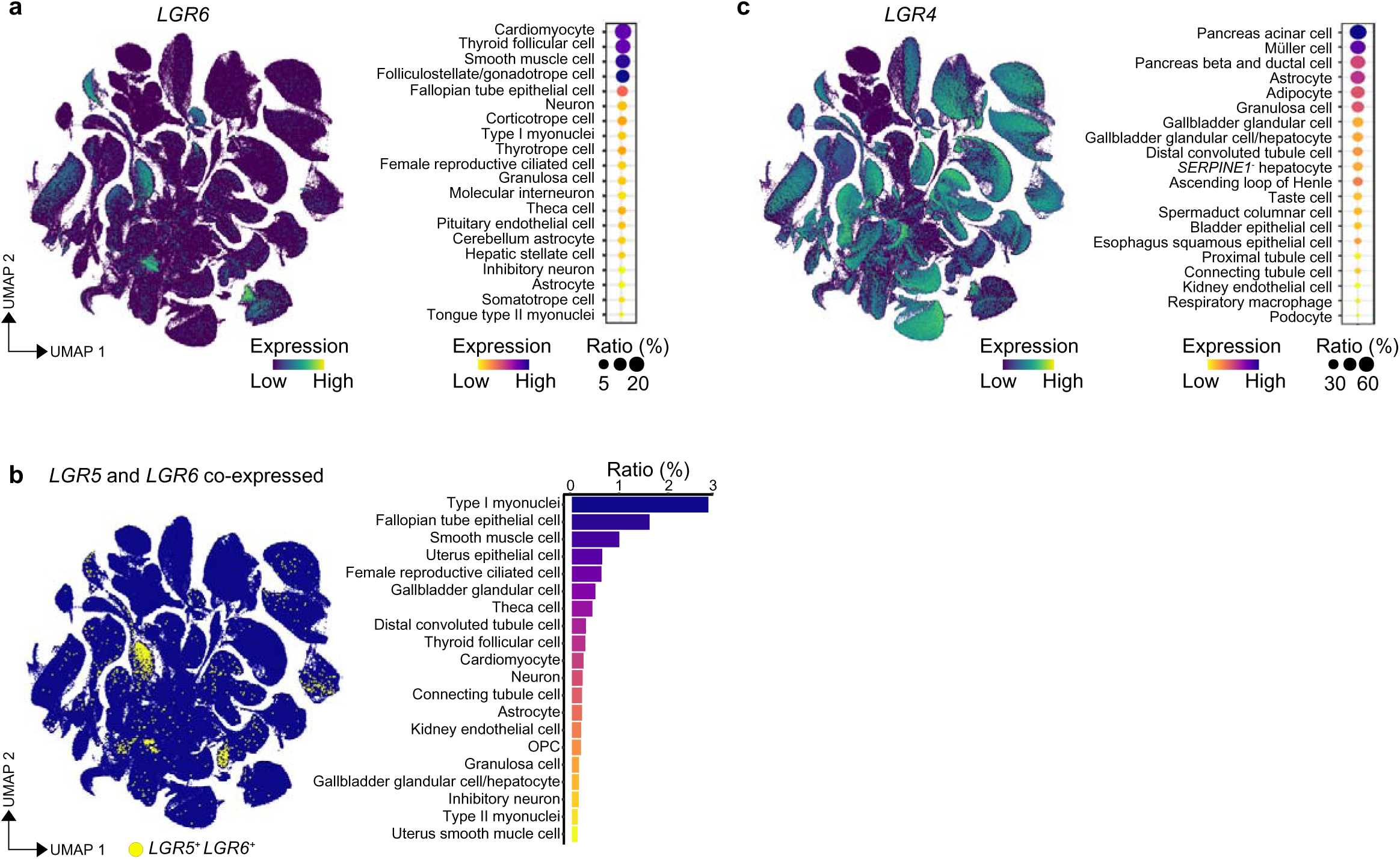
Global analysis of *LGR4, LGR6* and *LGR5/LGR6* co-expression across monkey tissues. **(a)** UMAP visualization of *LGR6* across all tissues profiled in this study. The bubble plot on the right shows the *LGR6* expression ratio in the indicated cell types. **(b)** UMAP visualization of *LGR5* and *LGR6* co-expression across all tissues profiled in this study. The barplot on the right shows the co-expression ratio in the indicated cell types. **(c)** UMAP visualization of *LGR4* across all tissues profiled in this study. The bubble plot on the right shows the *LGR4* expression ratio in the indicated cell types.

**Extended Data Figure 19.**
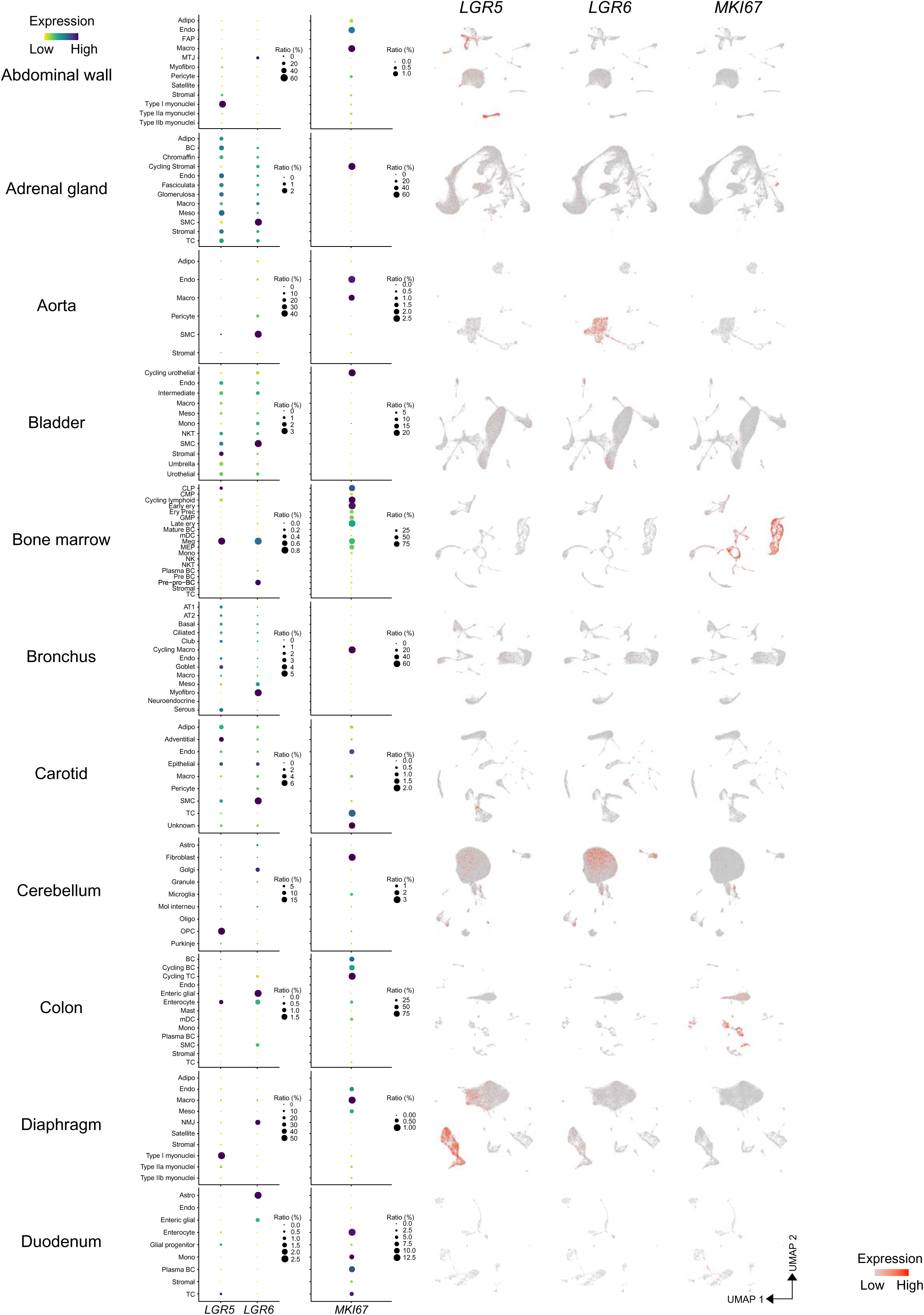
Global analysis of *LGR5* and *LGR6* across monkey tissues – 1. Bubble plot (left) showing the ratio of *LGR5*^+^, *LGR6*^+^ and *MKI67*^+^ cells in the annotated cell types for each tissue and UMAP visualization (right) of *LGR5*, *LGR6* and *MKI67* in abdominal wall, adrenal gland, aorta, bladder, bone marrow, bronchus, carotid, cerebellum, colon, diaphragm and duodenum.

**Extended Data Figure 20.**
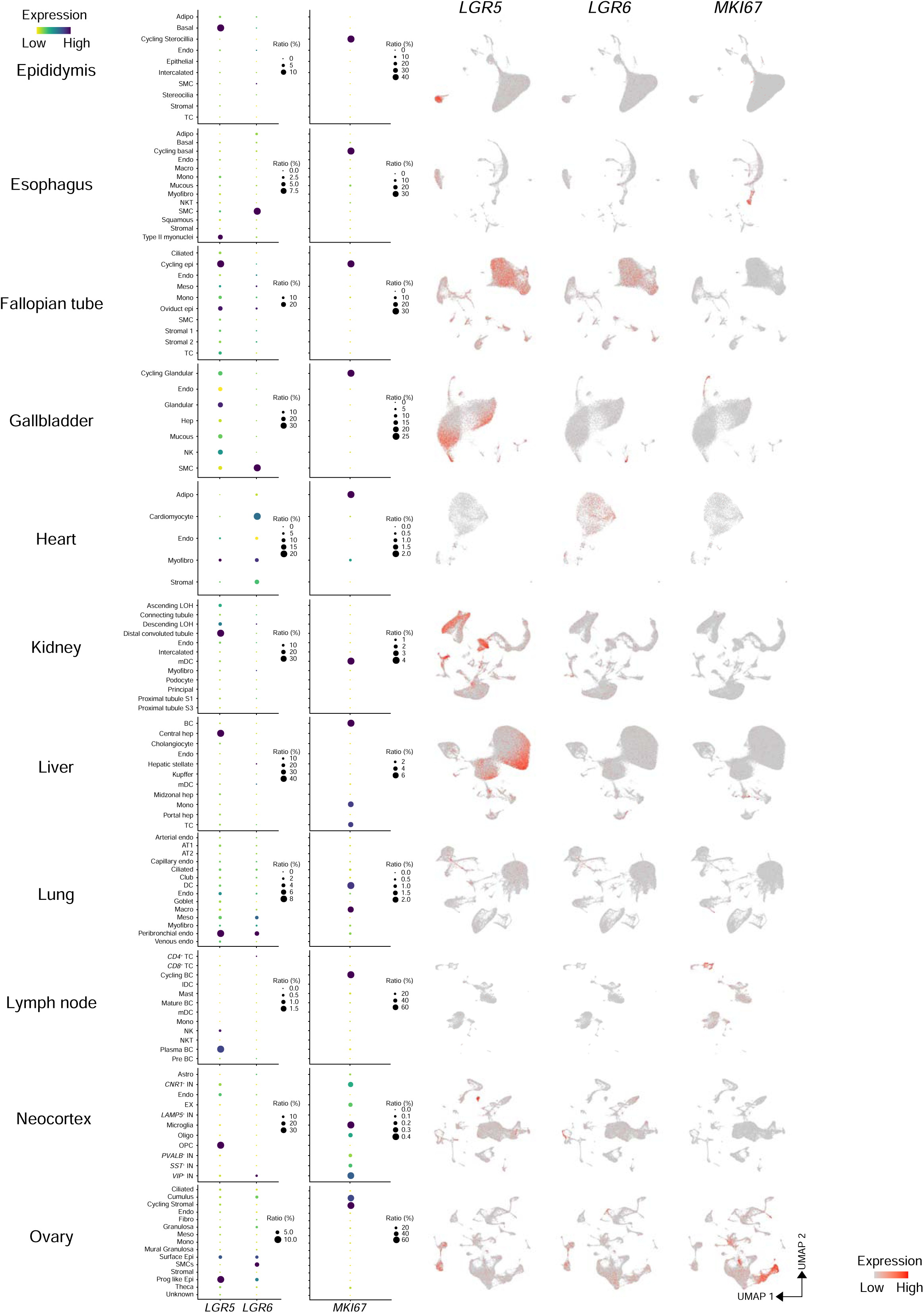
Global analysis of *LGR5* and *LGR6* across monkey tissues – 2. Bubble plot (left) showing the ratio of *LGR5*^+^, *LGR6*^+^ and *MKI67*^+^ cells in the annotated cell types for each tissue and UMAP visualization (right) of *LGR5*, *LGR6* and *MKI67* in epididymis, esophagus, fallopian tube, gallbladder, heart, kidney, liver, lung, lymph node and ovary.

**Extended Data Figure 21.**
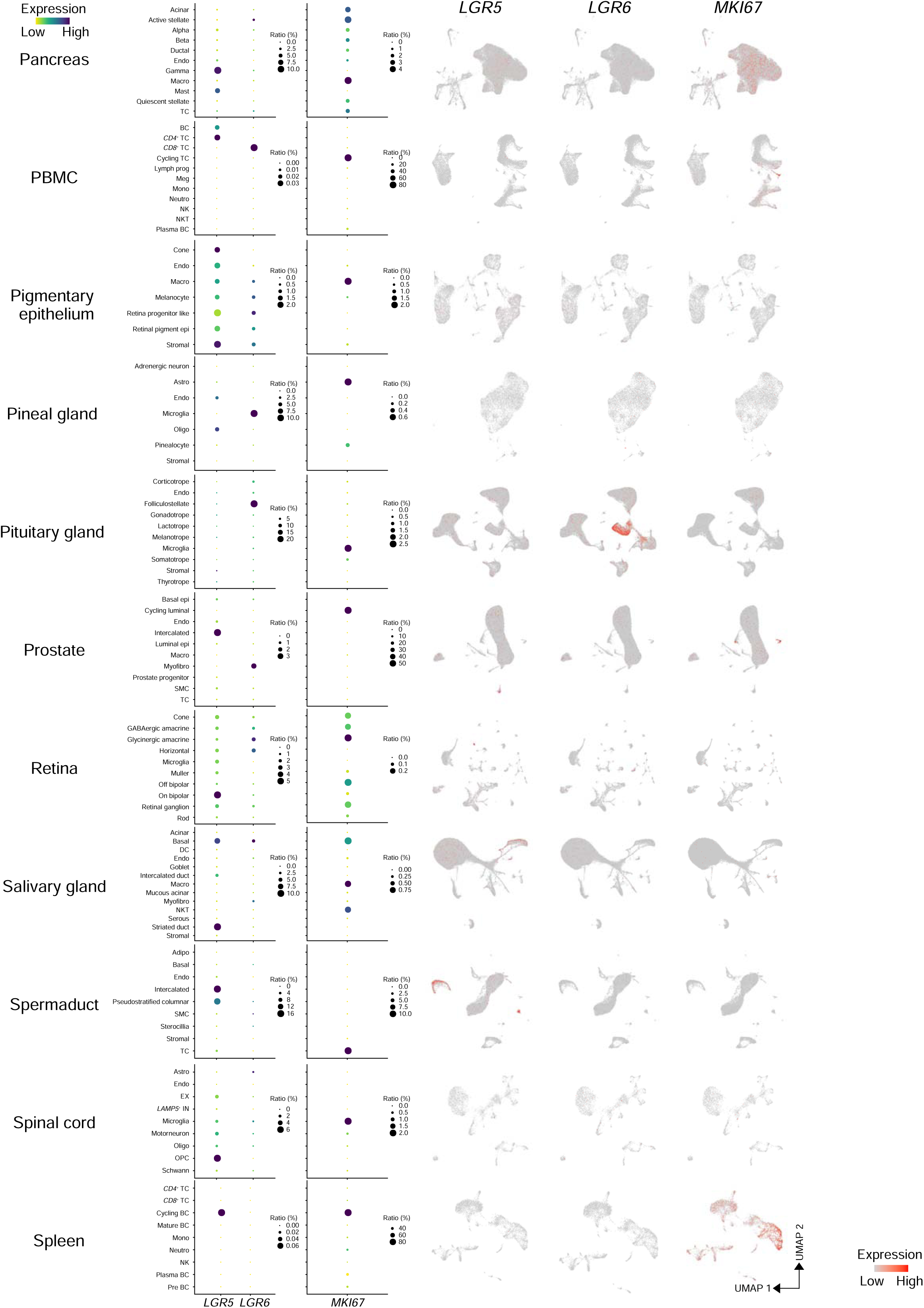
Global analysis of *LGR5* and *LGR6* across monkey tissues – 3. Bubble plot (left) showing the ratio of *LGR5*^+^, *LGR6*^+^ and *MKI67*^+^ cells in the annotated cell types for each tissue and UMAP visualization (right) of *LGR5*, *LGR6* and *MKI67* in pancreas, PBMCs, pigmentary epithelium choroid plexus (indicated as pigmentary epi), pineal gland, pituitary gland, prostate, retina, salivary gland, spermaduct, spinal cord and spleen.

**Extended Data Figure 22.**
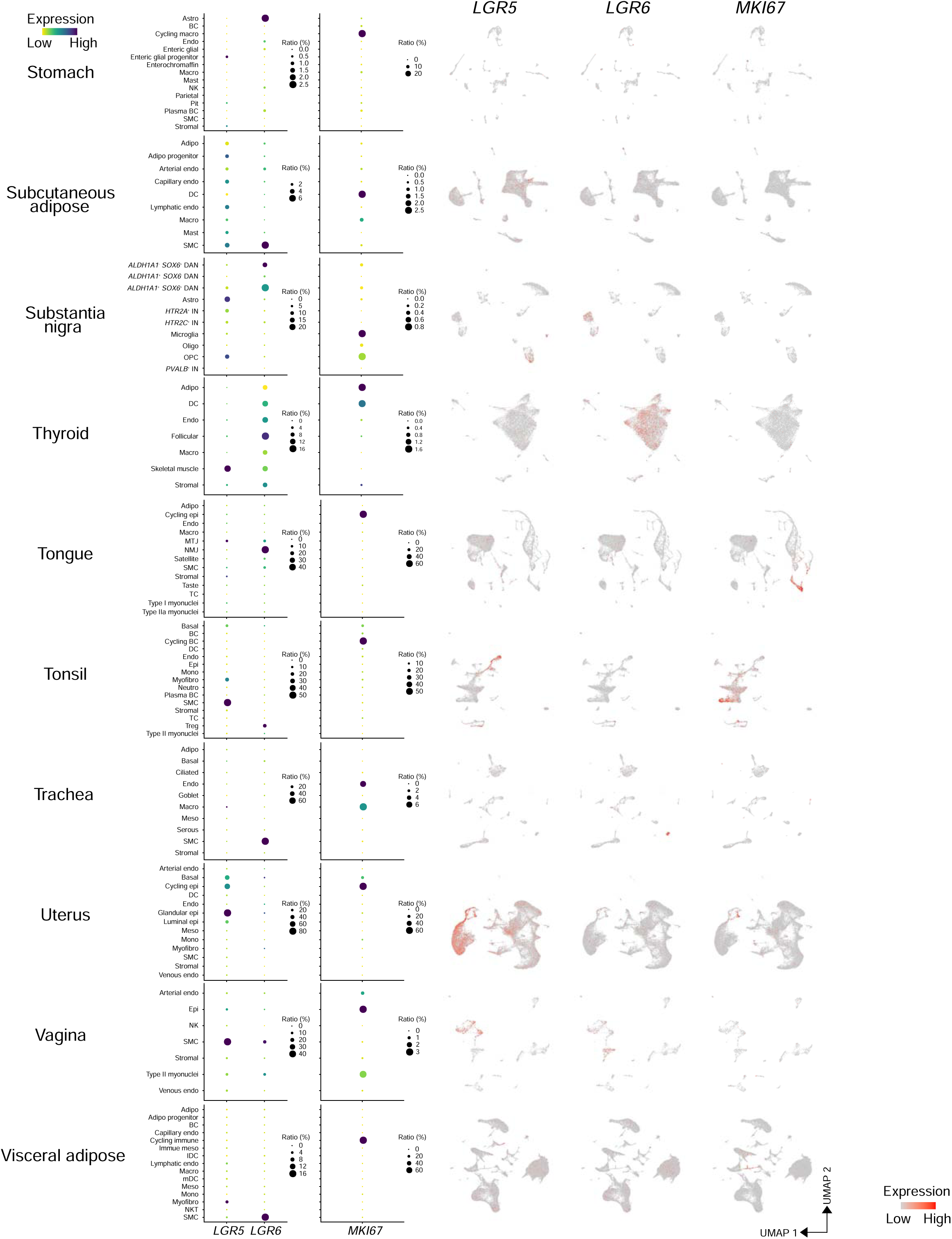
Global analysis of *LGR5* and *LGR6* across monkey tissues – 4. Bubble plot (left) showing the ratio of *LGR5*^+^, *LGR6*^+^ and *MKI67*^+^ cells in the annotated cell types for each tissue and UMAP visualization (right) of *LGR5*, *LGR6* and *MKI67* in stomach, subcutaneous adipose tissue, substantia nigra, thyroid, tongue, tonsil, trachea, uterus, vagina and visceral adipose tissue.

**Extended Data Figure 23.**
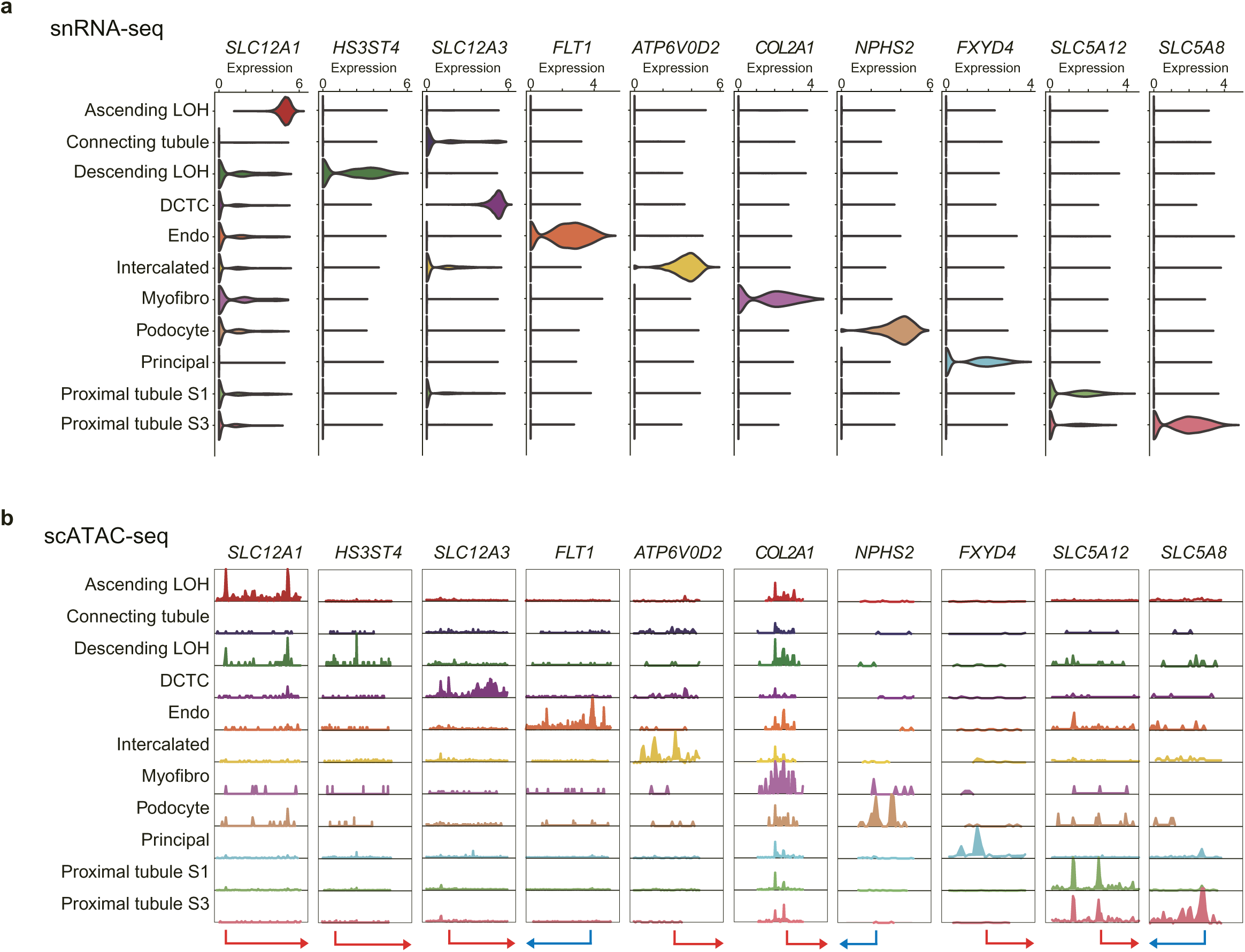
Kidney snRNA-seq and scATAC-seq dataset integration. **(a)** Violin plot showing the expression of selected markers used to annotate the kidney cell clusters from snRNA-seq data. **(b)** ArchR track visualization of aggregate scATAC-seq signals on the locus of the selected marker genes indicated in **a**. Abbreviations: DCTC, distal convoluted tubule cells; Endo, endothelial cells; LOH, loop of Henle; Myofibro, myofibroblasts.

**Extended Data Figure 24.**
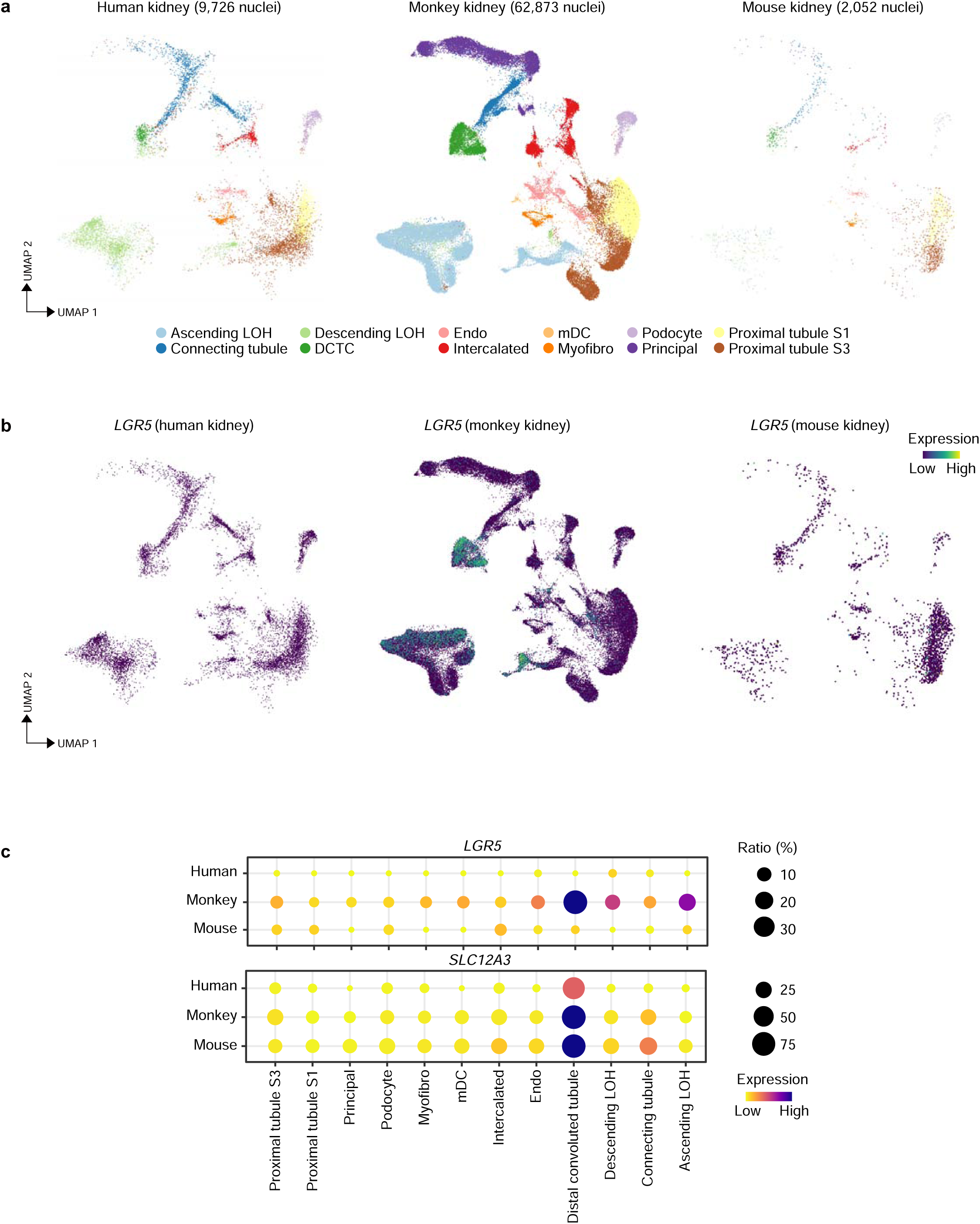
*LGR5* analysis in integrated human, monkey and mouse kidney data. **(a)** UMAP visualization of cell clusters in human (left), monkey (middle) and mouse (right) kidney snRNA-seq datasets. The annotation of each cluster in provided in the legend at the bottom. Abbreviations: Endo, endothelial cells; LOH, loop of Henle; mDC, myeloid dendritic cells; Myofibro, myofibroblasts. **(b)** UMAP visualization of *LGR5* in human (left), monkey (middle) and mouse (right) kidney. **(c)** Bubble plot showing the ratio and expression levels of *LGR5* and DCTC marker *SLC12A3* in human, monkey and mouse kidney datasets. The color of each bubble represents the level of expression and the size indicates the proportion of expressing cells.

**Extended Data Figure 25.**
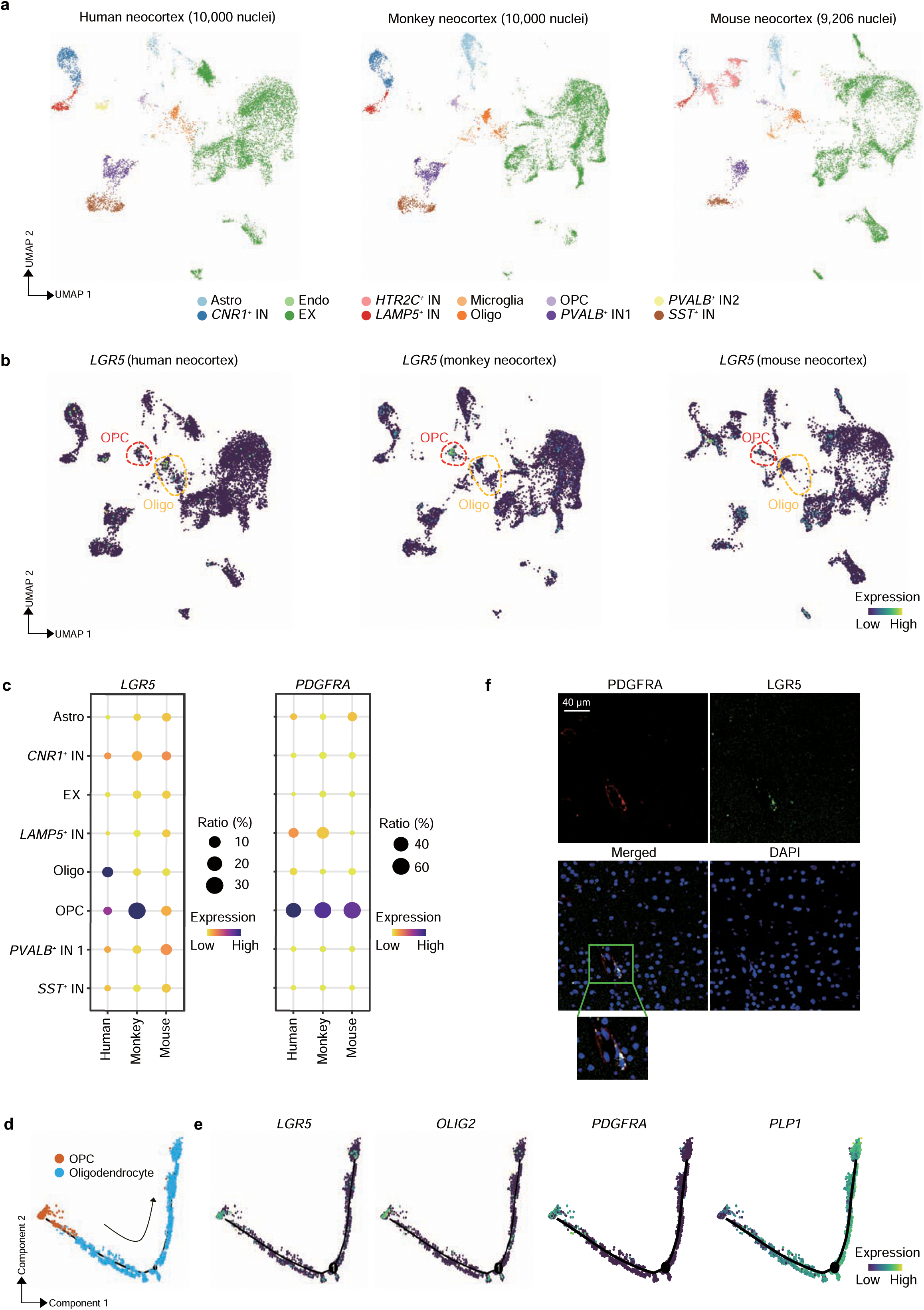
*LGR5* analysis in integrated human, monkey and mouse neocortex data. **(a)** UMAP visualization of cell clusters in human (left), monkey (middle) and mouse (right) neocortex snRNA-seq datasets. The annotation of each cluster is provided in the legend at the bottom. Abbreviations: Astro, astrocytes; Endo, endothelial cells; IN, inhibitory neurons; OPC, oligodendrocyte progenitor cells; EX, excitatory neurons; Oligo, oligodendrocytes. **(b)** UMAP visualization of *LGR5* in human (left), monkey (middle) and mouse (right) neocortex. OPC and oligodendrocytes are indicated by a red and yellow dotted circle, respectively. **(c)** Bubble plot showing the ratio and expression levels of *LGR5* and *PDGFRA* in human, monkey and mouse neocortex. The color of each bubble represents the level of expression and the size indicates the proportion of expressing cells. **(d)** Monocle 2 pseudotime-ordered trajectory of OPC (labelled in orange) maturation towards mature oligodendrocytes (labelled in blue). **(e)** Monocle 2 pseudotime analysis showing the expression of OPC markers (*LGR5*, *OLIG2* and *PDGFRA*) and the oligodendrocytes marker *PLP1*. **(f)** Representative image of immunofluorescence staining for PDGFRA (red) and LGR5 (green), respectively, and their co-expression in OPC of monkey neocortex (scale bar 20 μm). The smaller panel at the bottom is a magnification of the area indicated by the green box.

**Extended Data Figure 26.**
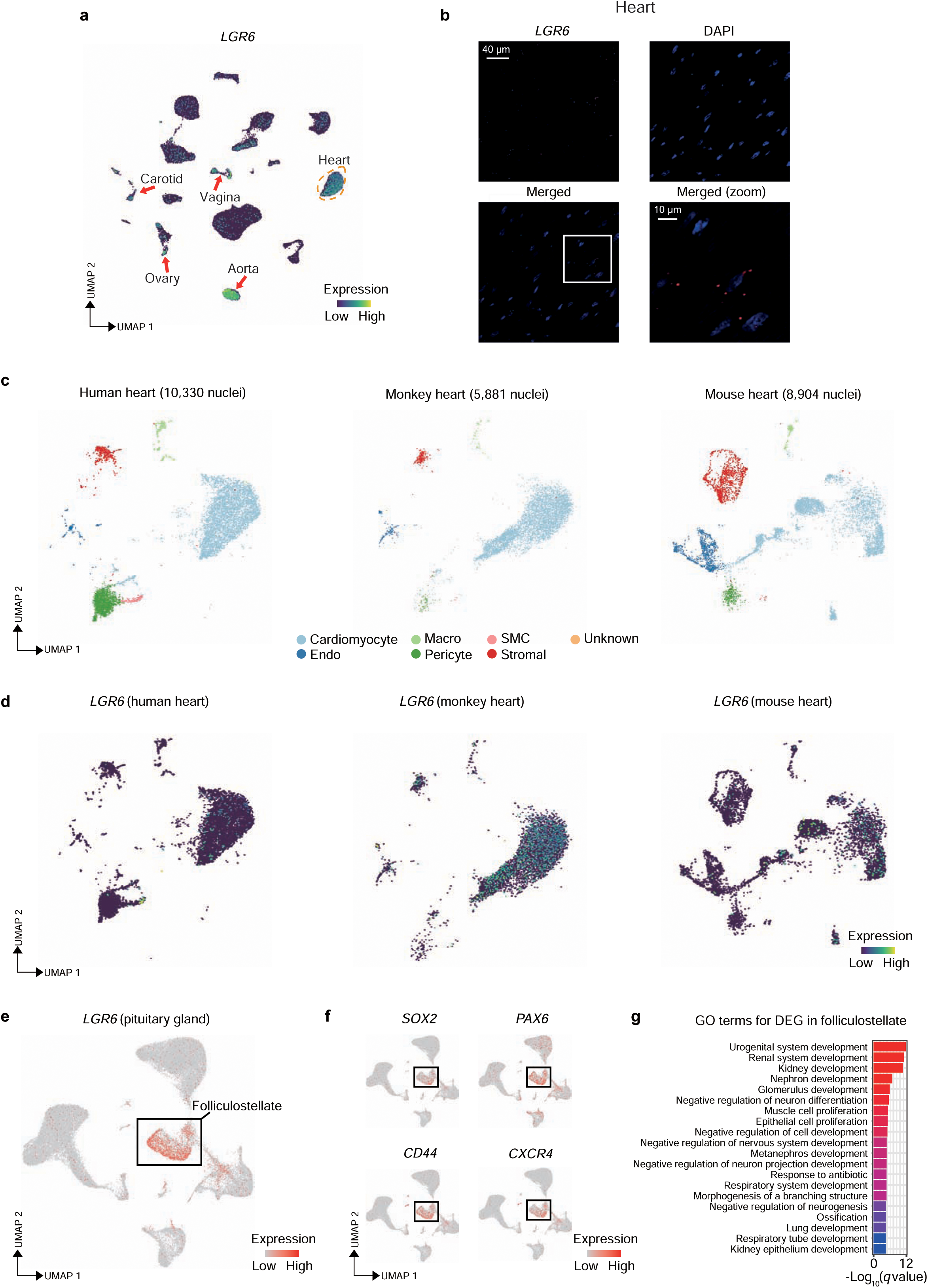
Analysis of *LGR6* expression in monkey heart and pituitary gland. **(a)** UMAP visualization of *LGR6* across all muscle cell types annotated in our dataset, as displayed in Figure 3g. The dotted red line indicates a cluster of muscle cells belonging to the heart. The red arrows indicate *LGR6*^+^ cells in aorta, carotid, ovary and vagina. **(b)** Representative image of smFISH detection for *LGR6* in heart myonuclei (scale bar 40 μm). The bottom right panel is a magnification of the area indicated by the white box. **(c)** UMAP visualization of cell clusters in human (left), monkey (middle) and mouse (right) heart snRNA-seq datasets. The annotation of each cluster is provided in the legend at the bottom. Abbreviations: Endo, endothelial cells; Macro, macrophages; SMC, smooth muscle cells. **(d)** UMAP visualization of *LGR6* in human (left), monkey (middle) and mouse (right) heart. **(e)** UMAP visualization of *LGR6* expression in pituitary gland highlighting the highest expression in folliculostellate cells. **(f)** UMAP visualization of *SOX2*, *PAX6*, *CD44* and *CXCR4* in folliculostellate cells as indicated by the black box. **(g)** Barplot showing GO terms associated to the DEGs in folliculostellate cells of pituitary gland.

**Extended Data Figure 27.**
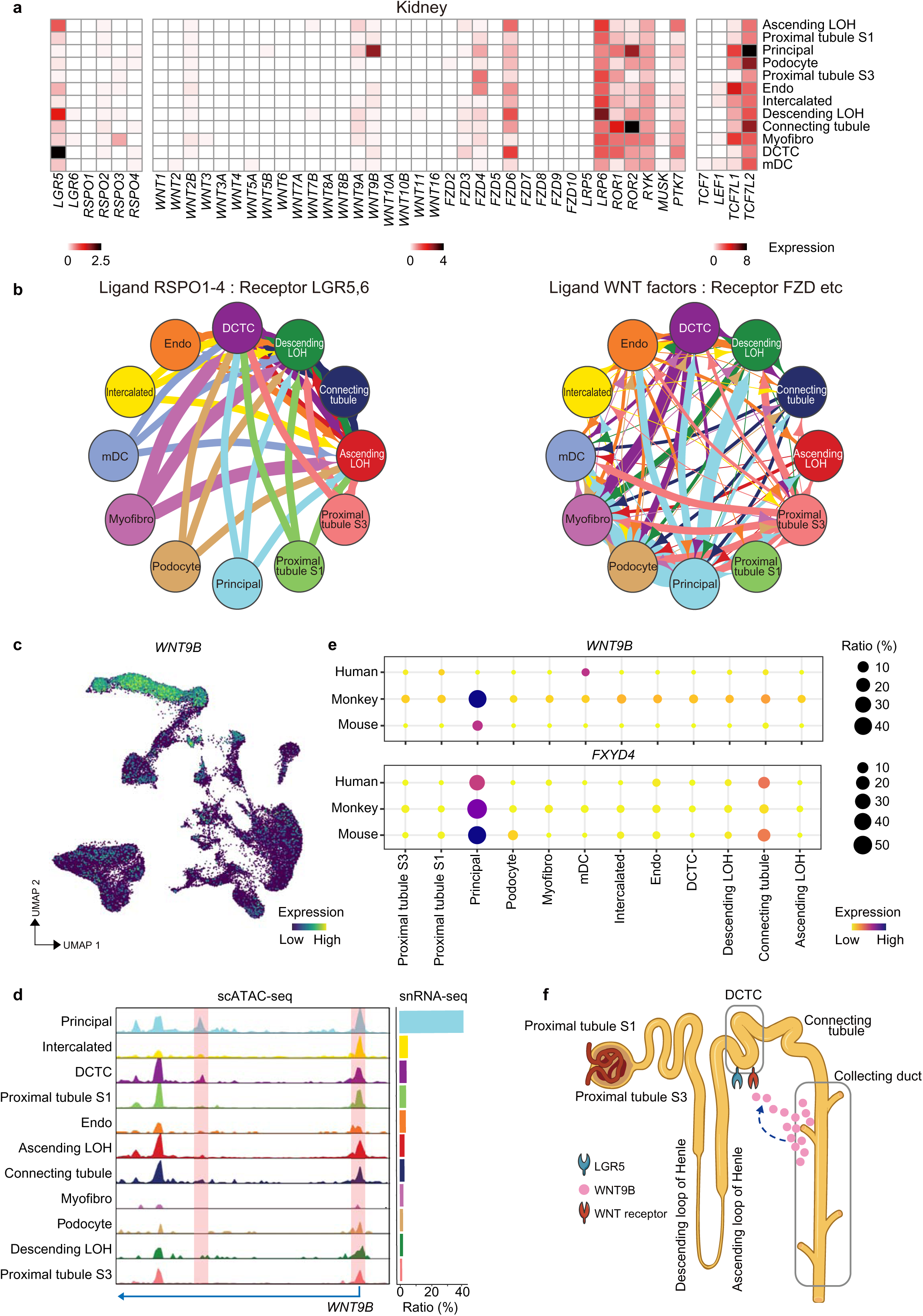
Analysis of *WNT9B* and Wnt pathway gene module in monkey kidney. **(a)** Heatmap showing the expression of all receptors and ligands of the Wnt pathway in the annotated cell populations of the kidney. **(b)** Network plots showing cell-cell communications based on ligand-receptor interactions calculated by CellphoneDB. **(c)** UMAP visualization of *WNT9B* expression in monkey kidney. **(d)** ArchR track visualization of aggregate scATAC-seq signals on the *WNT9B* locus in each on the annotated cell types. The bar plot at the bottom indicates the ratio (%) of *WNT9B*^+^ cells in each cell type of kidney. **(e)** Bubble plot showing the ratio and expression levels of *WNT9B* and principal tubule cell marker *FXYD4* in human, monkey and mouse kidney datasets. The color of each bubble represents the level of expression and the size indicates the proportion of expressing cells. **(f)** Schematic representation of a kidney nephron illustrating Wnt pathway ligand-receptor interactions.

**Extended Data Figure 28.**
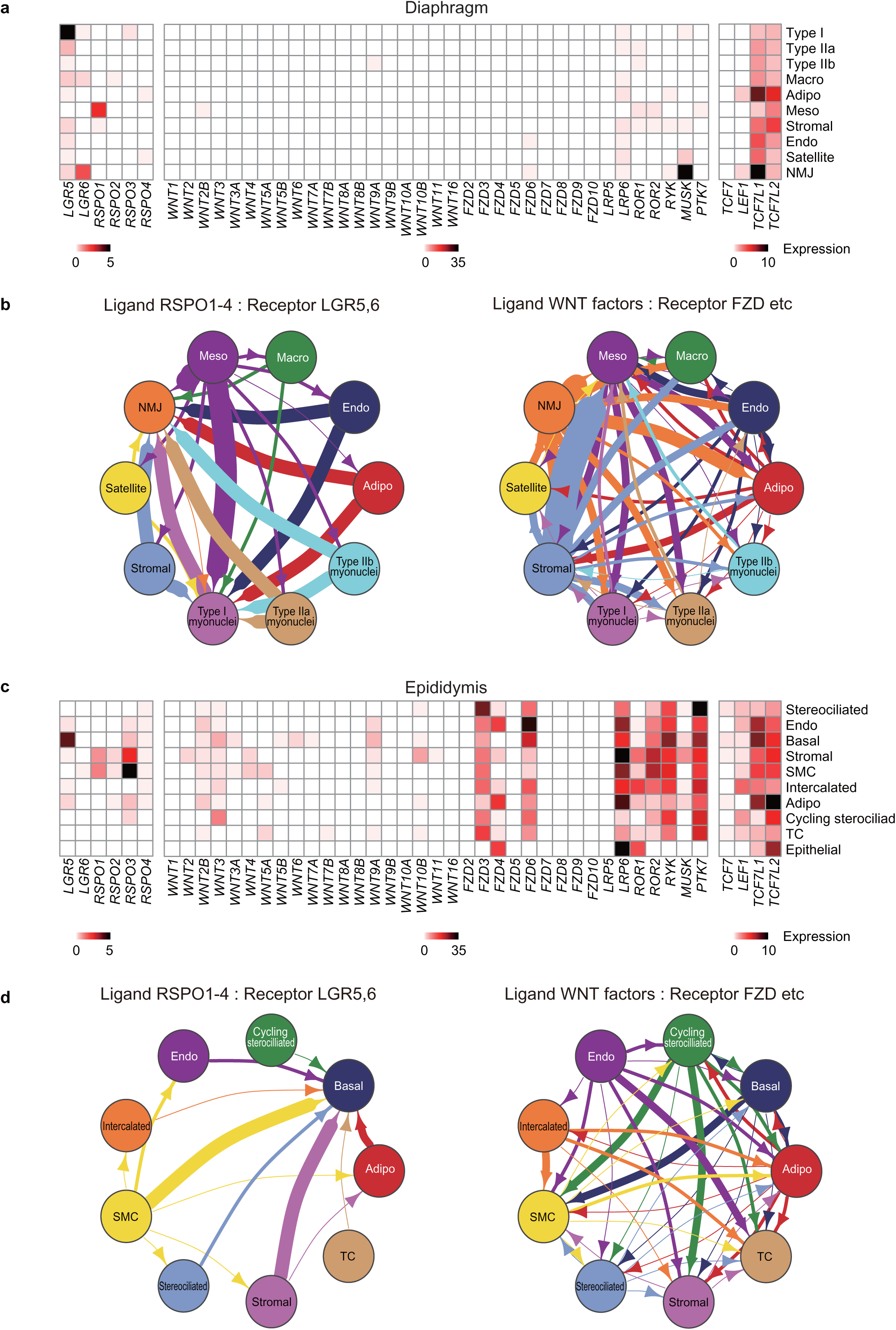
Global analysis of the Wnt pathway gene module in monkey diaphragm and epididymis. **(a)** Heatmap showing the expression of all receptors and ligands of the Wnt pathway in the annotated cell populations of the diaphragm. **(b)** Network plots showing cell-cell communication based on ligand-receptor interactions calculated by CellphoneDB in the diaphragm dataset. Abbreviations: Adipo, adipocytes; Endo, endothelial cells; Macro, macrophages; Meso, mesothelial cells; NMJ, neuromuscular junctions. **(c)** Heatmap showing the expression of all receptors and ligands of the Wnt pathway in the annotated cell populations of the epididymis. **(d)** Network plots showing cell-cell communication based on ligand-receptor interactions calculated by CellphoneDB in the epididymis dataset. Abbreviations: Adipo, adipocytes; Endo, endothelial cells; SMC, smooth muscle cells; TC, T cells.

**Extended Data Figure 29.**
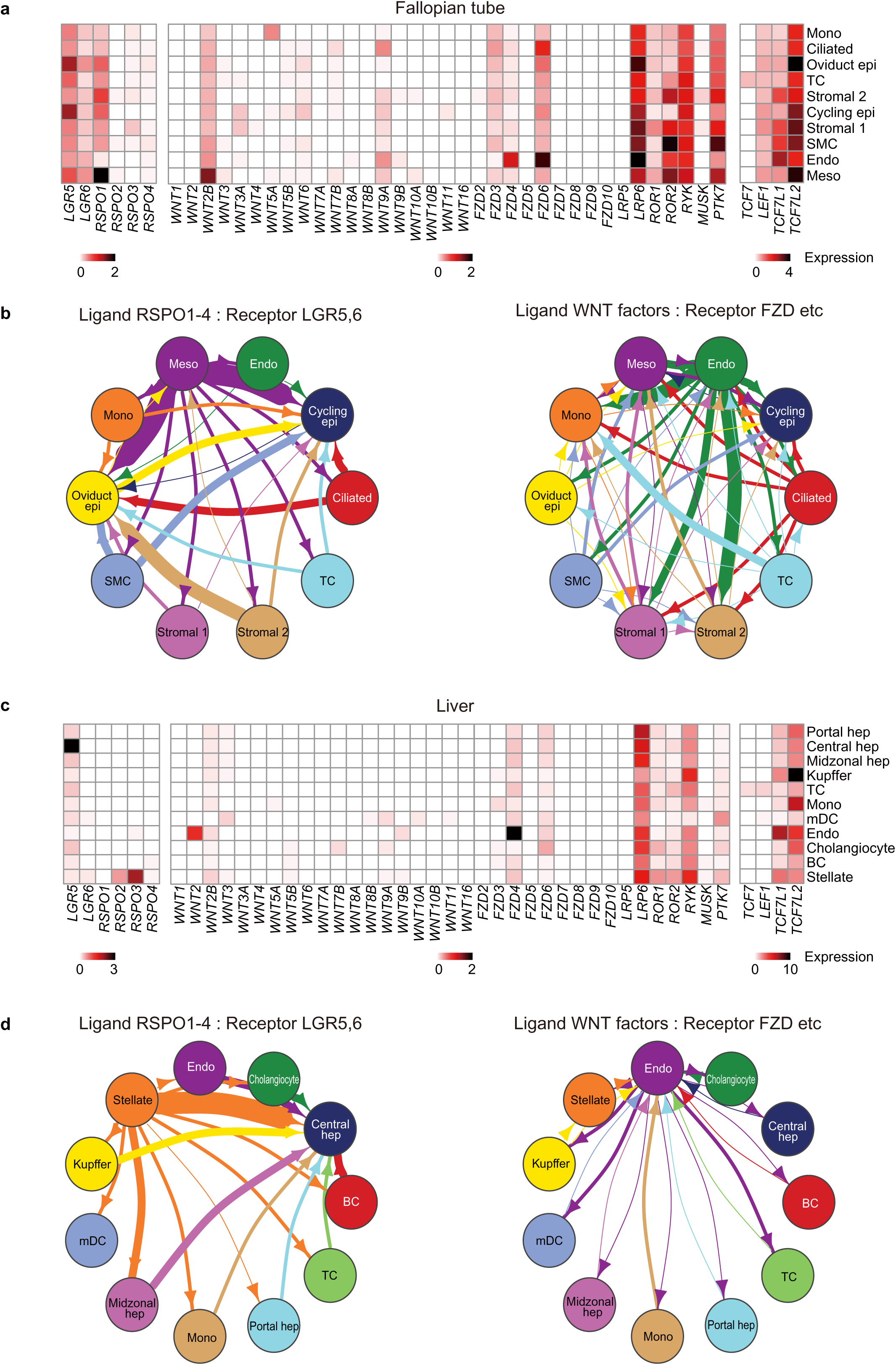
Global analysis of the Wnt pathway gene module in monkey fallopian tube and liver. **(a)** Heatmap showing the expression of all receptors and ligands of the Wnt pathway in the annotated cell populations of the fallopian tube. **(b)** Network plots showing cell-cell communication based on ligand-receptor interactions calculated by CellphoneDB in the fallopian tube dataset. Abbreviations: Endo, endothelial cells; epi, epithelial cells; Meso, mesothelial cells; Mono, monocytes; SMC, smooth muscle cells; TC, T cells. **(c)** Heatmap showing the expression of all receptors and ligands of the Wnt pathway in the annotated cell populations of the liver. **(d)** Network plots showing cell-cell communication based on ligand-receptor interactions calculated by CellphoneDB in the liver dataset. Abbreviations: BC, B cells; Endo, endothelial cells; hep, hepatocytes; mDC, myeloid derived dendritic cells; Mono, monocytes; TC, T cells.

**Extended Data Figure 30.**
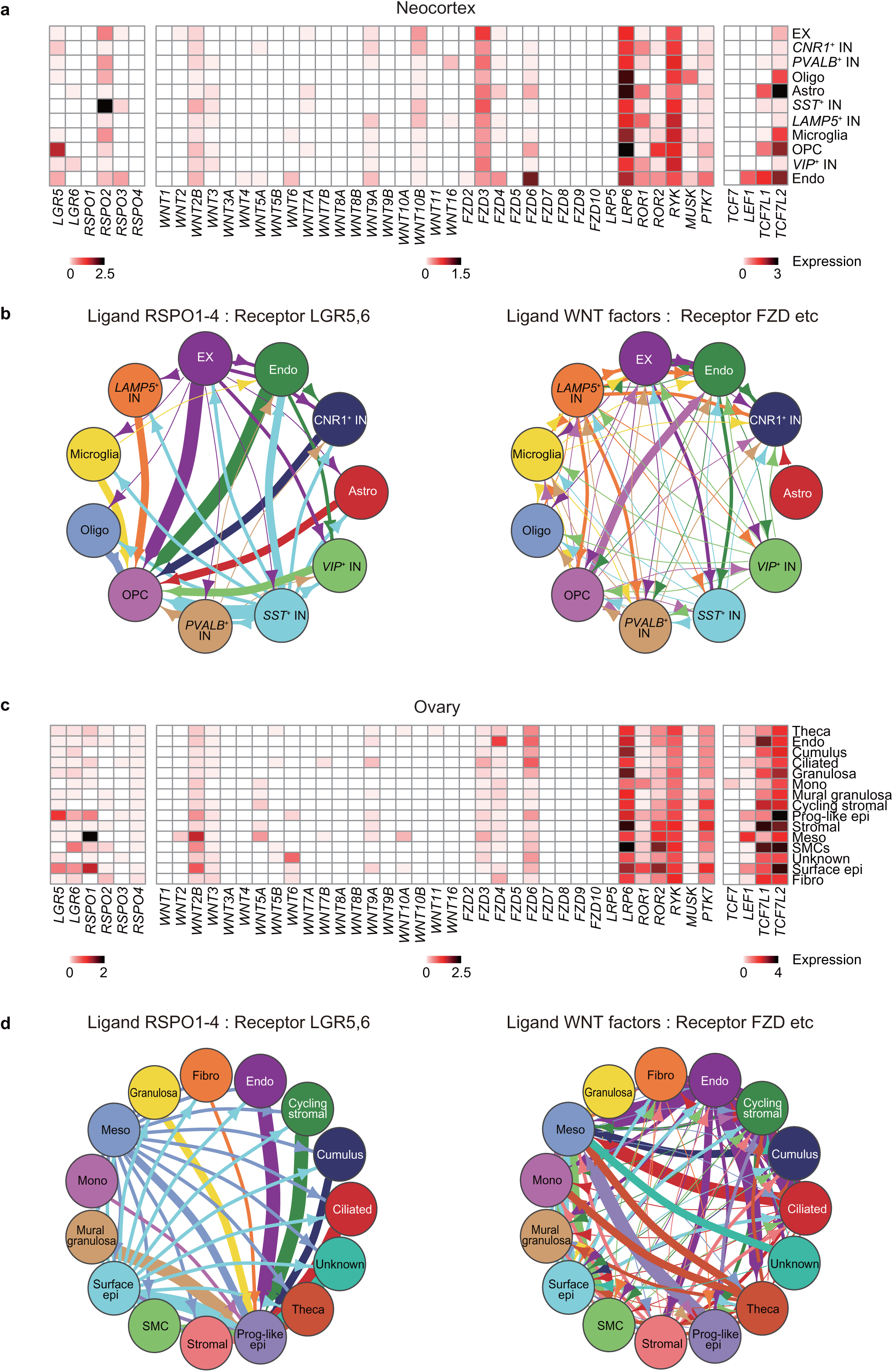
Global analysis of the Wnt pathway gene module in monkey neocortex and ovary. **(a)** Heatmap showing the expression of all receptors and ligands of the Wnt pathway in the annotated cell populations of the neocortex. **(b)** Network plots showing cell-cell communication based on ligand-receptor interactions calculated by CellphoneDB in the neocortex dataset. Abbreviations: Astro, astrocytes; Endo, endothelial cells; EX, excitatory neurons; IN, inhibitory neurons; Oligo, oligodendrocytes; OPC, oligodendrocyte progenitor cells. **(c)** Heatmap showing the expression of all receptors and ligands of the Wnt pathway in the annotated cell populations of the ovary. **(d)** Network plots showing cell-cell communication based on ligand-receptor interactions calculated by CellphoneDB in the ovary dataset. Abbreviations: Endo, endothelial cells; epi, epithelial cells; Meso, mesothelial cells; Mono, monocytes; Myofibro, myofibroblasts; Prog-like epi; progenitor-like epithelial cells; SMC, smooth muscle cells.

**Extended Data Figure 31.**
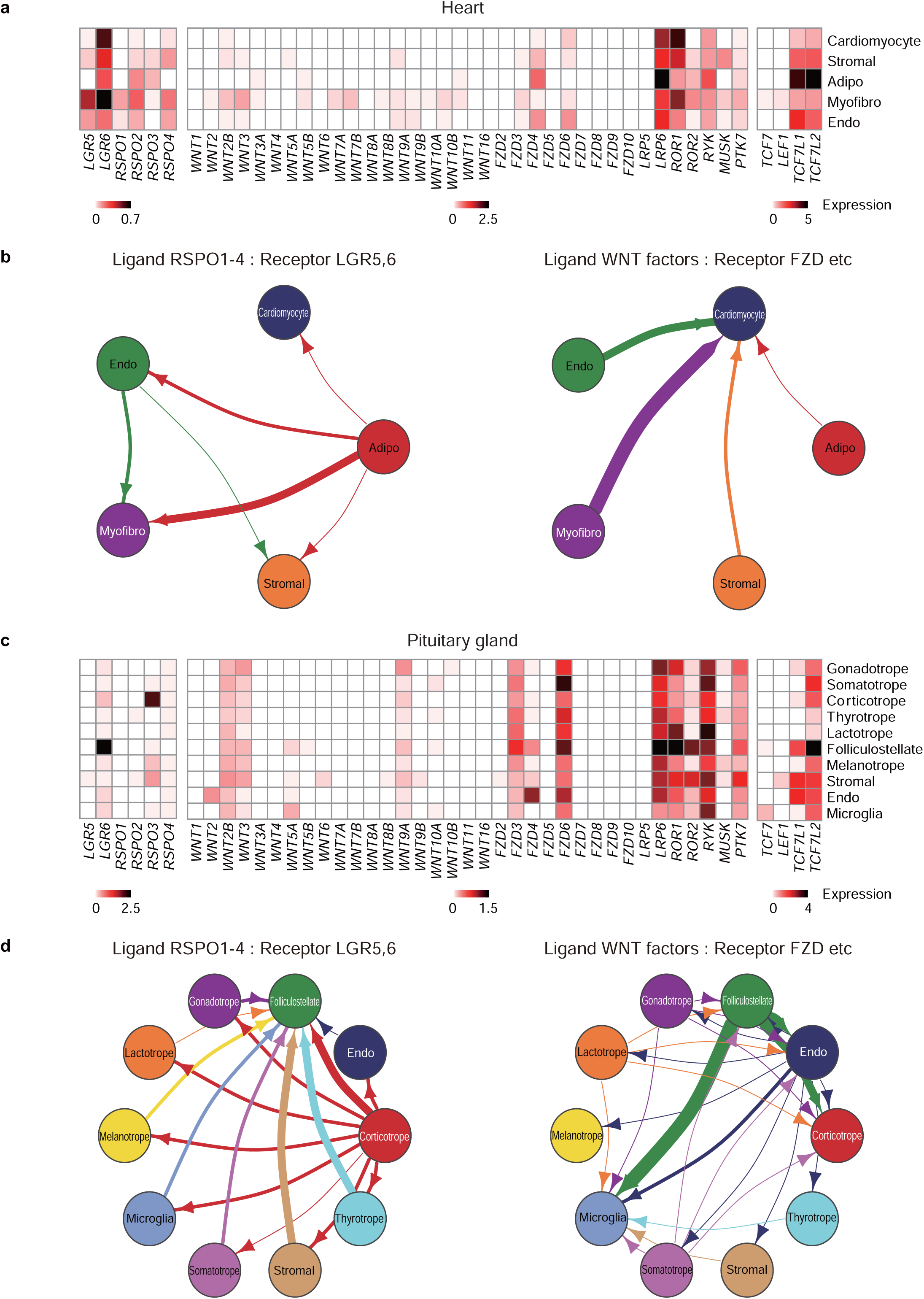
Global analysis of the Wnt pathway gene module in other monkey heart and pituitary gland. **(a)** Heatmap showing the expression of all receptors and ligands of the Wnt pathway in the annotated cell populations of the heart. **(b)** Network plots showing cell-cell communication based on ligand-receptor interactions calculated by CellphoneDB in the pituitary gland dataset. Abbreviations: Endo, endothelial cells; Myofibro, myofibroblasts. **(c)** Heatmap showing the expression of all receptors and ligands of the Wnt pathway in the annotated cell populations of the pituitary gland. **(d)** Network plots showing cell-cell communication based on ligand-receptor interactions calculated by CellphoneDB in the pituitary gland dataset. Abbreviations: Endo, endothelial cells.

**Extended Data Figure 32.**
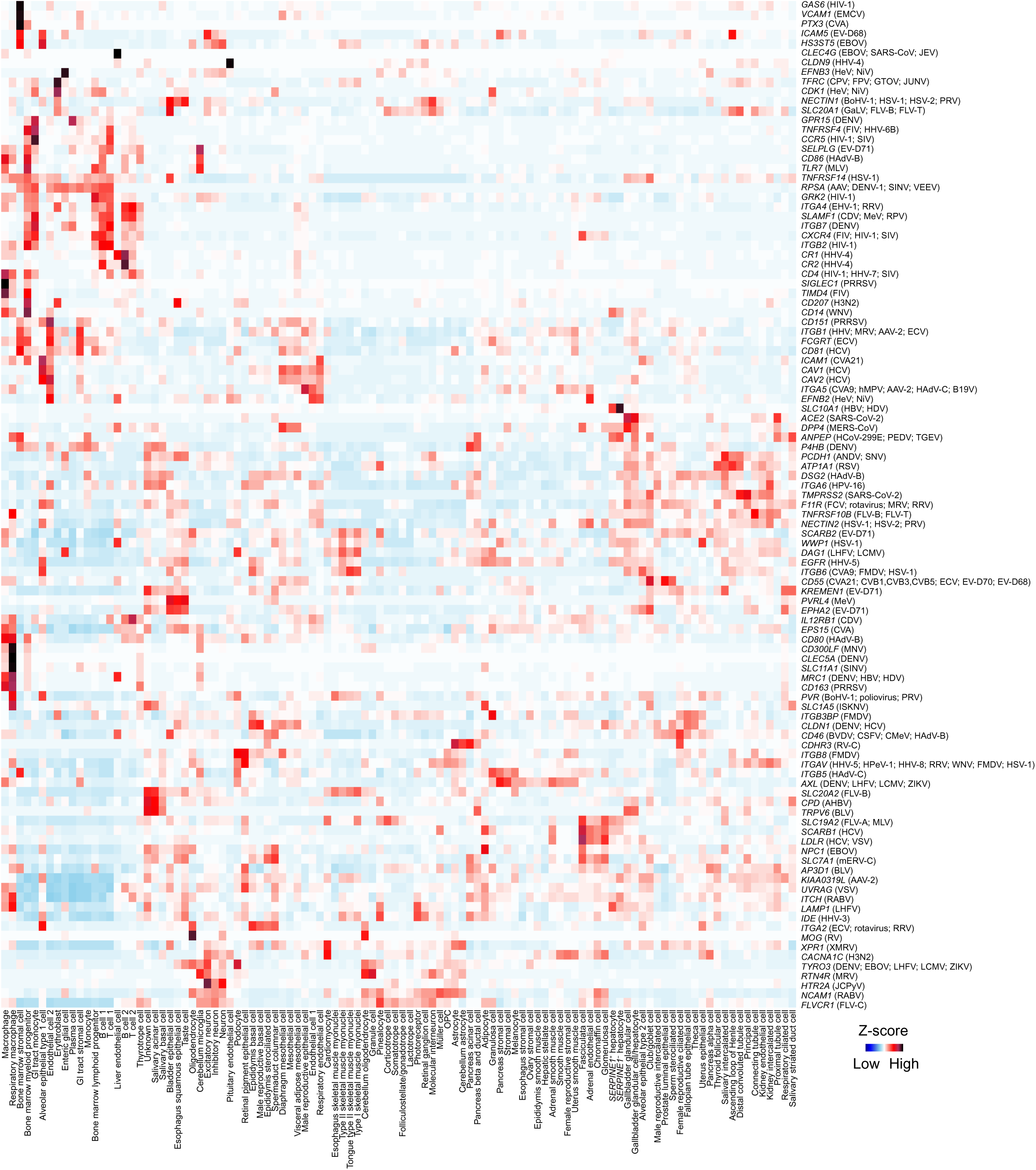
Global analysis of virus entry receptors across monkey tissues. Heatmap showing the expression of entry receptor for most common viruses (shown on the right) in the indicated cell types (shown at the bottom).

**Extended Data Figure 33.**
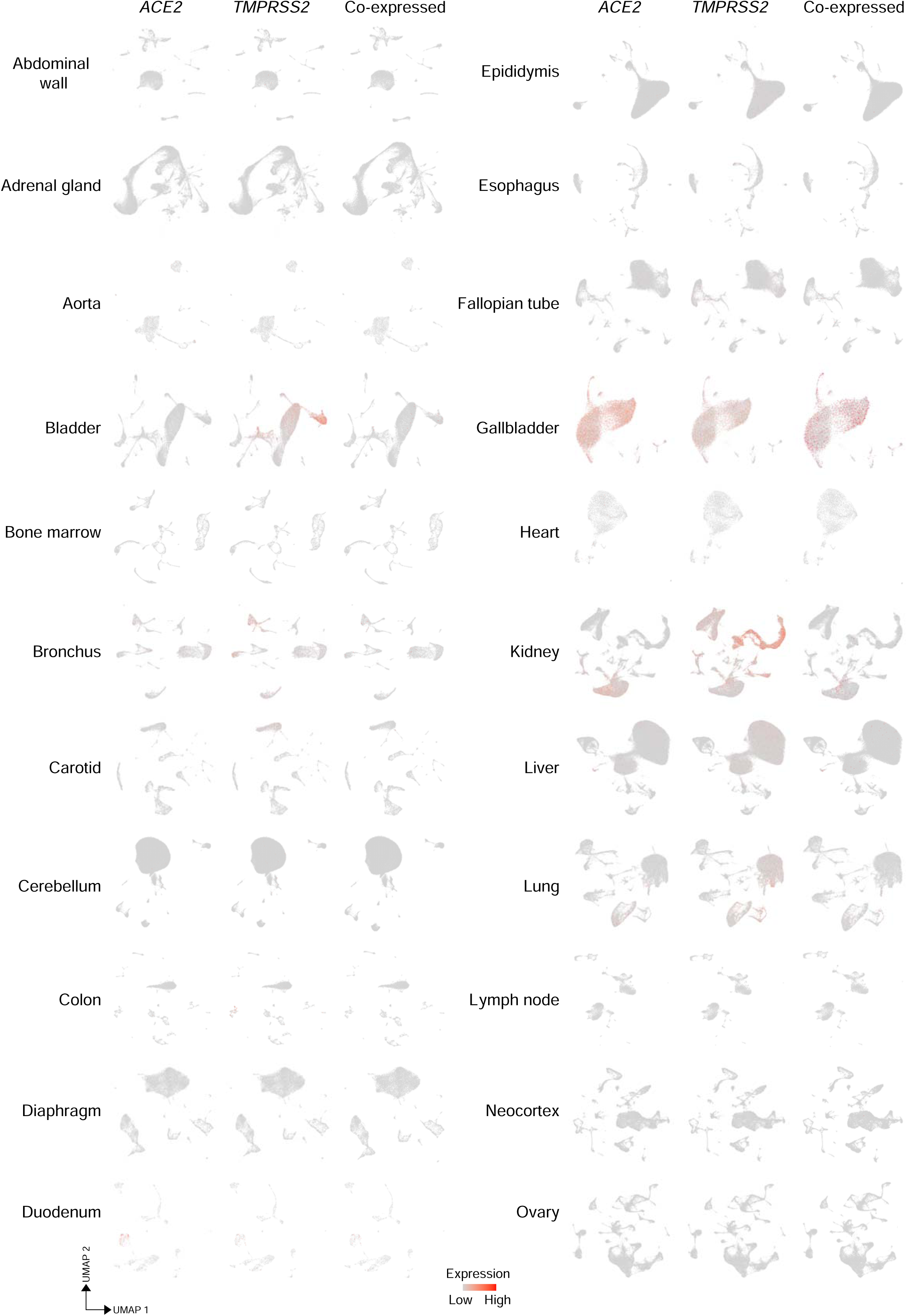
Analysis of *ACE2* and *TMPRSS2* expression across monkey tissues – 1. UMAP visualization of *ACE2* (left), *TMPRSS2* (middle) and *ACE2^+^*/*TMPRSS2^+^* (right) in abdominal wall, adrenal gland, aorta, bladder, bone marrow, bronchus, carotid, cerebellum, colon, diaphragm, duodenum, epididymis, esophagus, fallopian tube, gallbladder, heart, kidney, liver, lung, lymph node and ovary.

**Extended Data Figure 34.**
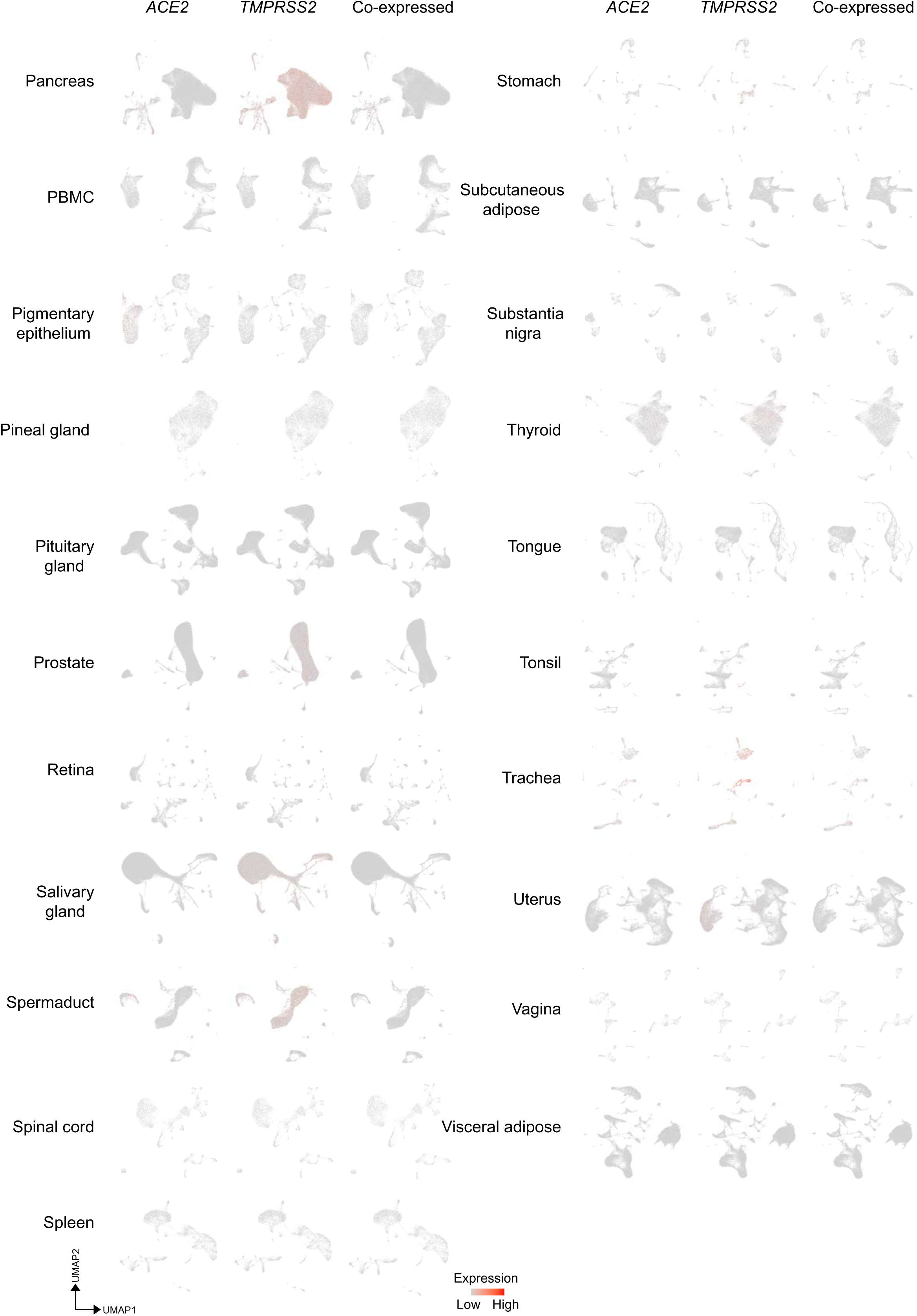
Analysis of *ACE2* and *TMPRSS2* expression across monkey tissues – 2. UMAP visualization of *ACE2* (left), *TMPRSS2* (middle) and *ACE2^+^*/*TMPRSS2^+^* (right) in pancreas, PBMC, pigmentary epithelium choroid plexus (indicated as pigmentary epi), pineal gland, pituitary gland, prostate, retina, salivary gland, spermaduct, spinal cord, spleen, stomach, subcutaneous adipose tissue, substantia nigra, thyroid, tongue, tonsil, trachea, uterus, vagina and visceral adipose tissue.

**Extended Data Figure 35.**
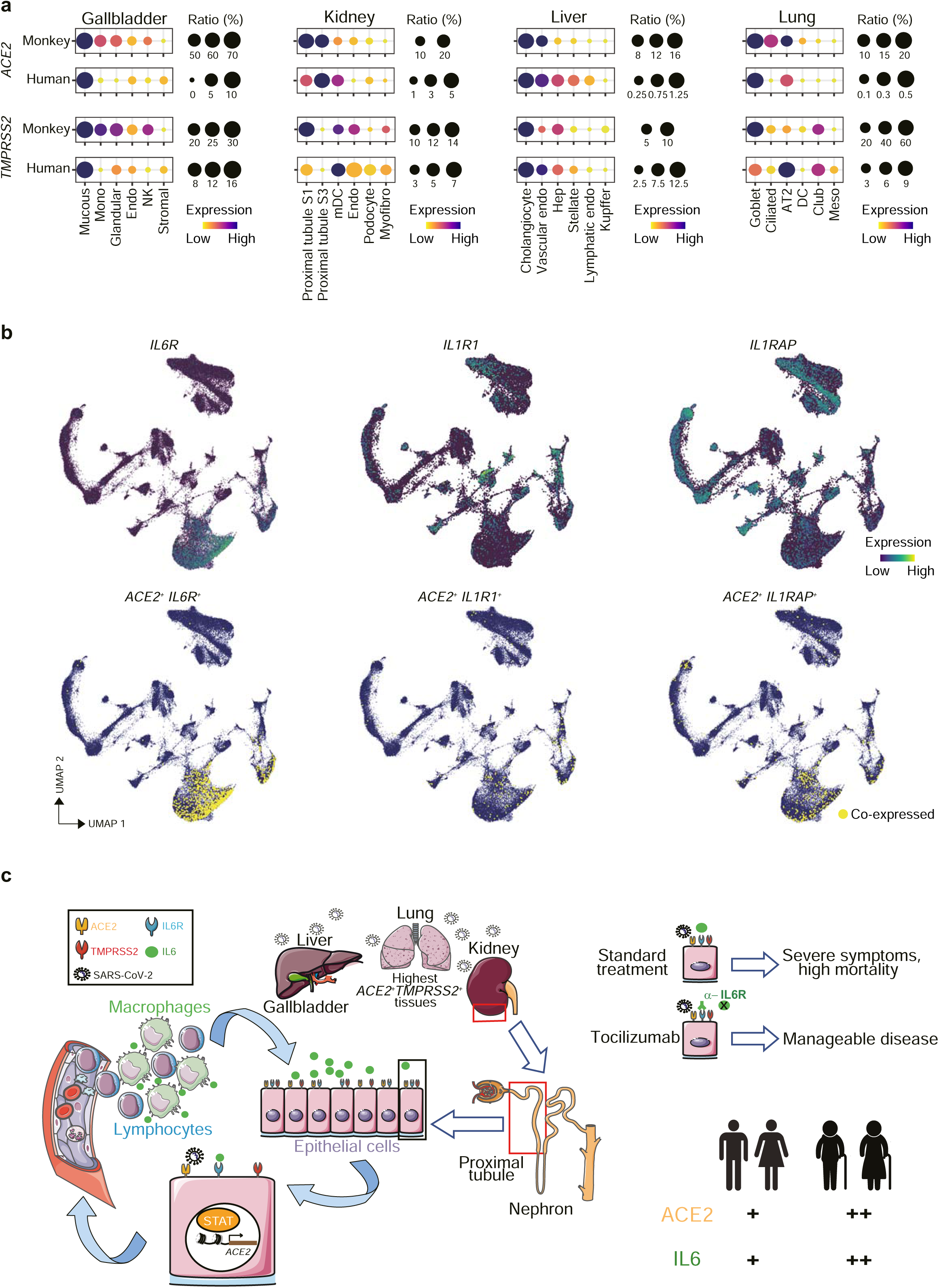
Comparative analysis of *ACE2* and *TMPRSS2* expression in monkey and human. **(a)** Bubble plot showing the ratio and expression levels of *ACE2* and *TMPRSS2* in gallbladder, kidney, liver and lung in monkey and human. The color of each bubble represents the level of expression and the size indicates the proportion of expressing cells. **(b)** UMAP visualization of *IL6R*, *IL1R1* and *IL1RAP* expressing in monkey kidney (top). The UMAP in the bottom represent the co-expression of *ACE2* and *IL6R*, *IL1R1* and *IL1RAP* in monkey kidney. Double positive cells are indicated in yellow. **(c)** Schematic diagram of the potential mechanism for SARS-CoV-2 spreading through gallbladder, kidney, liver and lung. Kidney proximal tubule cells within the nephron are among the highest ACE2 expressing cells. After virus contact, IL6R stimulates an immune response that, through the activation of STAT transcription factors, potentiates a paracrine positive feedback loop that enhances ACE2 expression and facilitates virus spreading. IL6 expression, which is higher in elderly patients and those with inflammatory conditions, is effectively targeted by anti-IL6R monoclonal antibodies leading to a more favourable disease course.

**Extended Data Figure 36.**
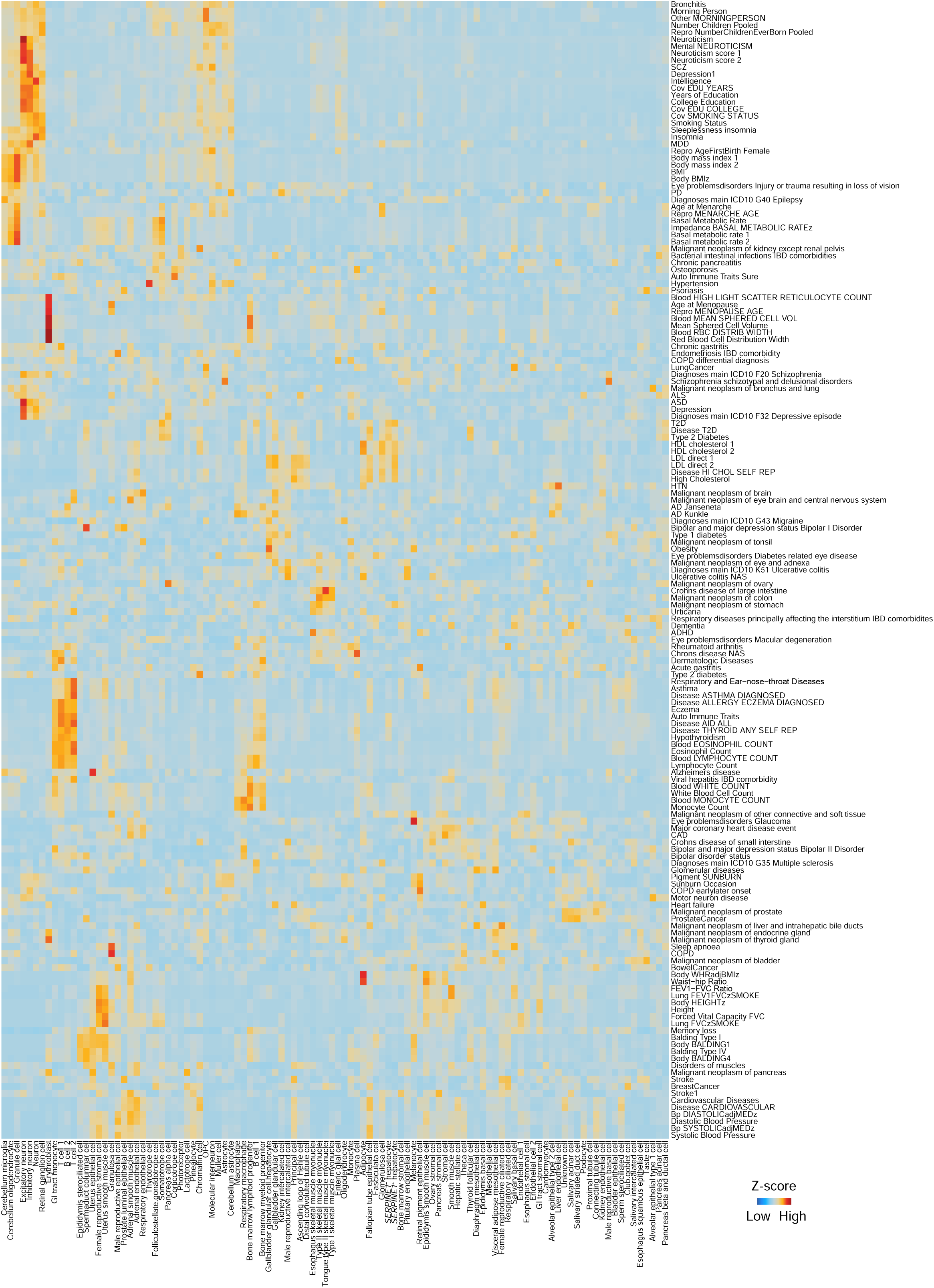
Expression of genes associated with human common traits in monkey cell types. Heatmap showing the association of common human traits and diseases from the UK Biobank (indicated on the right) with the cell types (indicated at the bottom) annotated in our dataset.

**Extended Data Figure 37.**
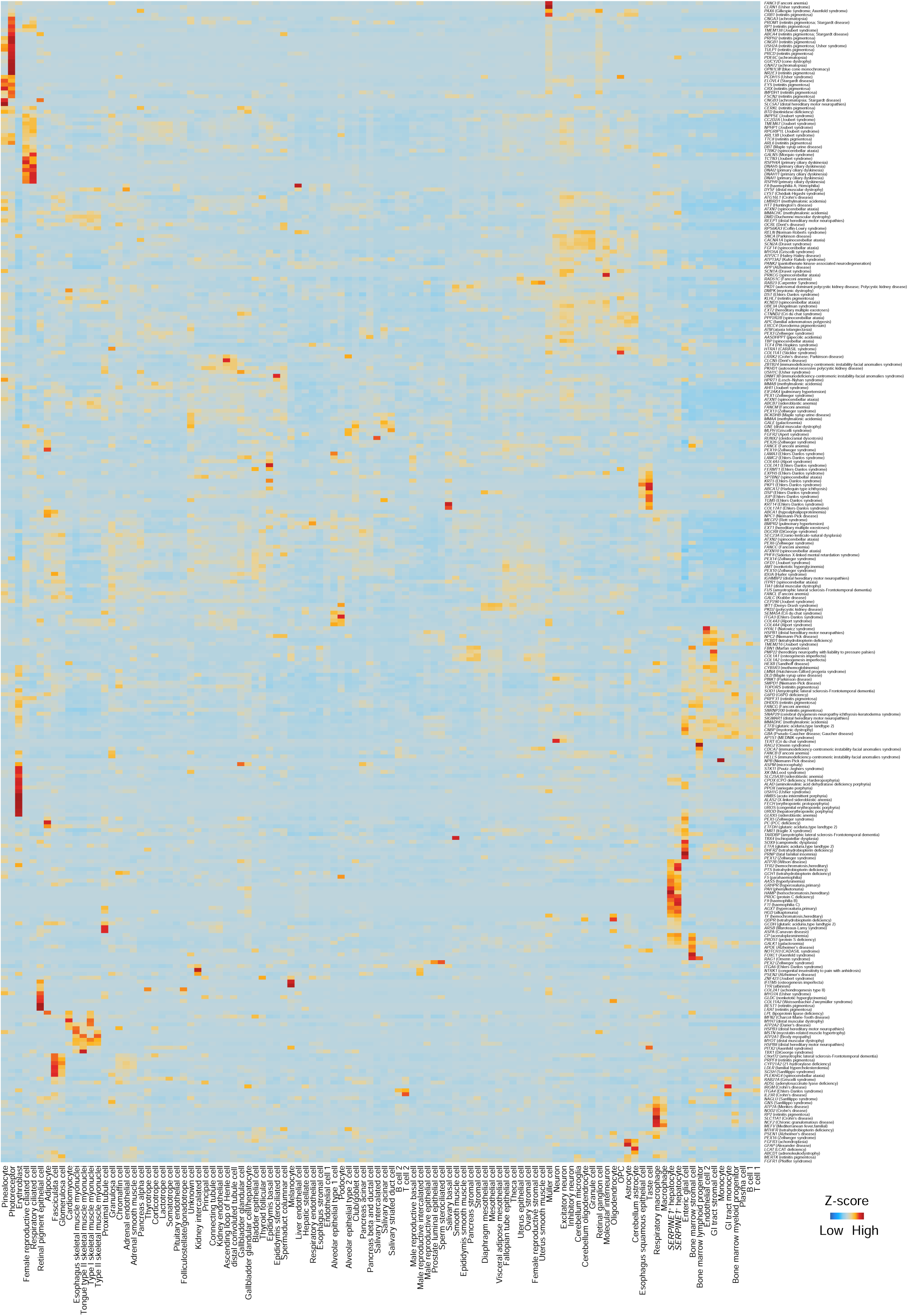
Association of monkey cell type-specific transcriptomic profiles with human genetic diseases. Heatmap showing the association of human genetic diseases (indicated on the right) with the cell types (indicated at the bottom) annotated in our dataset.

